# Extracellular matrix composition is associated with tissue-specific decellularization susceptibility and mechanical remodeling across human urogenital tissues

**DOI:** 10.64898/2026.08.28.747607

**Authors:** Stéphane Bolduc, Stéphane Chabaud, Arnaud Droit, Victor Fourcassié, Florence Roux-Dalvai, Yudaï Sahuc, Jayson Sueters

**Affiliations:** Centre de Recherche en Organogénèse Expérimentale/LOEX, Centre de Recherche du CHU de Québec-Université Laval, Axe Médecine Régénératrice, Québec, QC G1J 1Z4, Canada; Department of Surgery, Faculty of Medicine, Université Laval, Québec, QC G1V 0A6, Canada; Département de médecine moléculaire, Université Laval, Québec, QC G1V 0A6, Canada; Axe Endo-Nephro, Centre de recherche du CHU de Québec-Université Laval, Québec, QC G1V 0A6, Canada; Inria, Maasai team-Université Côte d’Azur, Nice, France

**Author notes:** Current address: Department of Urology, Faculty of Medicine, University Medical Centre Utrecht, Utrecht, the Netherlands (Jayson Sueters).

**Keywords:** tissue engineering, decellularized matrix, extracellular matrix, recellularization, biomaterial optimization, urethral reconstruction, proteomics, regenerative medicine

## Abstract

Decellularized extracellular matrices (ECMs) are widely used in regenerative medicine, yet current evaluation criteria prioritize cellular removal rather than preservation of the ECM characteristics that govern tissue behavior. Here, we demonstrate that efficient decellularization is achieved across a broad range of chemical conditions, whereas preservation of structurally and biologically relevant ECM components is confined to narrow, tissue-specific windows defined by coupled detergent interactions. Quantitative proteomics revealed that intrinsic ECM composition is strongly associated with tissue-specific susceptibility to decellularization-induced damage and provided molecular context for the distinct preservation responses between tissues. Optimized matrices retained major structural ECM components and supported tissue-specific cellular organization and cell-mediated mechanical reinforcement following cellular repopulation despite uniformly low residual DNA across protocols. Together, these findings support a shift in decellularization quality assessment from DNA-based evaluation toward preservation of biologically relevant ECM and establish a composition-driven strategy for the rational design of regenerative biomaterials with tissue-relevant biological and mechanical properties.

## 1. Introduction

Urethral anomalies, including hypospadias, urethral strictures, traumatic injury, spongiofibrosis, and gender incongruence requiring gender-affirming reconstruction, represent a major clinical challenge by compromising urinary function, sexual health, fertility and quality of life for millions of individuals worldwide.^1–5^ Despite more than 300 described urethroplasty techniques, reconstruction continues to rely predominantly on autologous tissues such as oral mucosa, skin, or *tunica vaginalis*. Although these grafts remain the clinical standard, they do not fully recapitulate the structural and biological characteristics of native urethral tissue and are therefore associated with stenosis, fibrosis, contracture, donor-site morbidity and variable long-term functional outcomes.^6–8^ These limitations have driven the development of tissue-engineered alternatives capable of restoring native tissue structure while reducing surgical complications. Early clinical studies have demonstrated the feasibility of engineered urethral substitutes, although widespread clinical translation remains limited.^9^

Decellularized matrices (DMs) represent one of the most promising biomaterials for tissue reconstruction because they retain key structural and biochemical features of the native extracellular matrix (ECM),^10,11^ providing structural support together with tissue-specific biochemical signals that regulate cell behavior,^12^ tissue organization and remodeling.^13–15^ Their translational potential has been demonstrated across numerous tissues, including vascular,^16^ bone,^17,18^ dermal^19^ and cardiac applications.^20^ In urogenital reconstruction, encouraging preclinical and early clinical studies have further highlighted the promise of biologically derived scaffolds for urethral repair, although outcomes remain variable.^9,21,22^ Despite these advances, clinical outcomes remain inconsistent, highlighting a fundamental limitation in the way DMs are currently evaluated.

The current definition of successful decellularization is primarily based on residual DNA content and histological absence of cellular material. These criteria were originally developed to minimize immunogenicity and standardize cellular removal rather than to evaluate functional matrix preservation.^23^ As a result, current standards effectively quantify cellular removal but provide little information about preservation of the ECM itself.^24^ Consequently, matrices that satisfy accepted decellularization thresholds may differ substantially in their composition, structural organization and mechanical properties. Increasing evidence indicates that aggressive decellularization can disrupt ECM structure and deplete biologically important matrix components, including basement membrane proteins and other functionally important ECM components, while maintaining efficient cellular removal.^25–28^ These alterations may compromise the biological microenvironment required for appropriate cell attachment, tissue organization, and long-term remodeling.^29–32^ Collectively, these observations suggest that successful decellularization cannot be defined solely by the extent of cellular removal but must also account for preservation of biologically relevant ECM characteristics.

A major obstacle to the rational design of decellularized biomaterials is the likelihood that susceptibility to decellularization differs between tissues. Native ECMs differ markedly in their molecular composition,^33^ ultrastructural organization^24^ and biological function,^34^ suggesting that tissues may respond differently to identical chemical treatments.^35^ Nevertheless, most decellularization protocols are typically optimized for individual tissues and subsequently evaluated using largely independent histological, biochemical and mechanical measurements. Consequently, it remains unknown whether intrinsic ECM composition governs tissue-specific susceptibility to decellularization-induced damage and thereby determines the biological quality of the resulting biomaterial.

Building on our previously established decellularized human vaginal matrix,^36,37^ which highlighted the importance of preserving ECM structures including the vasculature for regenerative biomaterials, the present study investigated whether intrinsic ECM composition determines tissue-specific susceptibility to decellularization-induced damage. Human urethral tissue was used as the primary model and structurally distinct glans tissue for comparison to systematically optimize detergent and enzymatic decellularization conditions and evaluate matrix preservation through integrated histological, biochemical, quantitative proteomic and mechanical analyses. This approach enabled investigation of whether tissue-specific ECM composition is associated with differential susceptibility to decellularization and subsequent structural, mechanical, and biological matrix preservation.

## 2. Results

### 2.1 Systematic optimization identifies tissue-specific narrow windows for successful decellularization

To identify decellularization conditions that simultaneously achieve efficient cellular removal and preservation of biologically relevant ECM components, a systematic optimization strategy was applied to human urogenital tissues. Donor-matched urethral samples were processed using up to three decellularization protocols per donor (Figure 1B). Seven iterative decellularization protocols (DC1-DC7) were evaluated by stepwise modulation of Triton X-100, sodium deoxycholate (SDC) and DNase I concentrations (Table 1A), with protocol refinement guided by histological assessment, DNA quantification, and DNA fragmentation analysis (Figure 1C).

**Figure 1.**
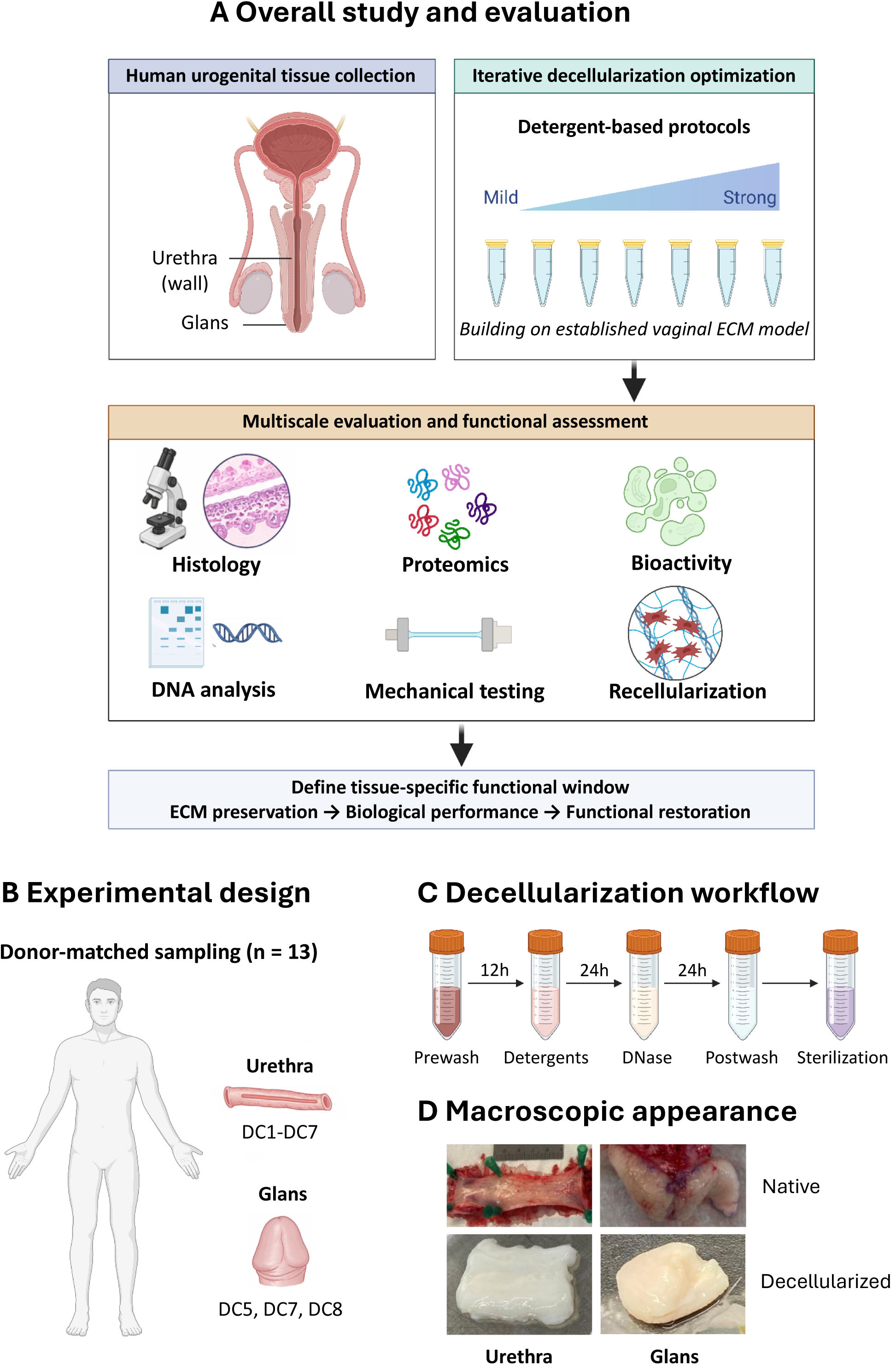
Decellularization strategy and workflow. **(A)** Overall study design. Human urethra and glans were subjected to detergent-based decellularization using iterative optimization and evaluation of histology, DNA analysis, proteomics, mechanical testing, bioactivity assay and recellularization. **(B)** Experimental design. Resected tissue yielded full-thickness urethra and matched glans from the same donors (n = 13). Urethra was exposed to protocols DC1-DC7 and glans to DC5, DC7 and DC8. **(C)** Decellularization workflow. Samples were extensively washed, exposed to detergents, treated with DNase I, extensively washed and exposed to sterilization and post-processing. **(D)** Macroscopic appearance: native (pink) versus decellularized tissue (milky-white, translucent). **(E)** Optimization of detergent conditions and their outcomes. ECM preservation and Cell removal in relation to the Triton X-100 and SDC concentrations.

**Table 1A:** Applied Triton X-100, SDC and DNase I concentrations for decellularization of urethra.

| Protocol urethra | Triton X-100 (%) | SDC (%) | DNase I |
| --- | --- | --- | --- |
| **DC1** | 0.1 | 0.1 | 200 IU/mL |
| **DC2** | 0.5 | 0.5 | 300 IU/mL |
| **DC3** | 1.0 | 1.0 | 400 IU/mL |
| **DC4** | 0.18 | 0.005 | 200 IU/mL |
| **DC5** | 0.18 | 0.015 | 300 IU/mL |
| **DC6** | 1.8 | 0.025 | 400 IU/mL |
| **DC7** | 0.20 | 0.005 | 200 IU/mL |

**Table 1B:** Applied Triton X-100, SDC and DNase I concentrations for decellularization of glans.

| Protocol glans | Triton X-100 (%) | SDC (%) | DNase I |
| --- | --- | --- | --- |
| **DC5** | 0.18 | 0.015 | 300 IU/mL |
| **DC7** | 0.20 | 0.005 | 200 IU/mL |
| **DC8** | 0.30 | 0.005 | 200 IU/mL |

Across all tested conditions, decellularization produced a consistent macroscopic transition from native pink tissue to a translucent, milky-white matrix (Figure 1D), indicating extensive cellular removal. Quantitative DNA analysis confirmed marked reductions in residual DNA across all protocols (Figure 2B), while DNA fragmentation analysis demonstrated efficient enzymatic degradation under every condition tested (Figure S1). Collectively, these findings indicate that effective cellular removal is robustly achieved across a broad range of chemical conditions.

**Figure 2.**
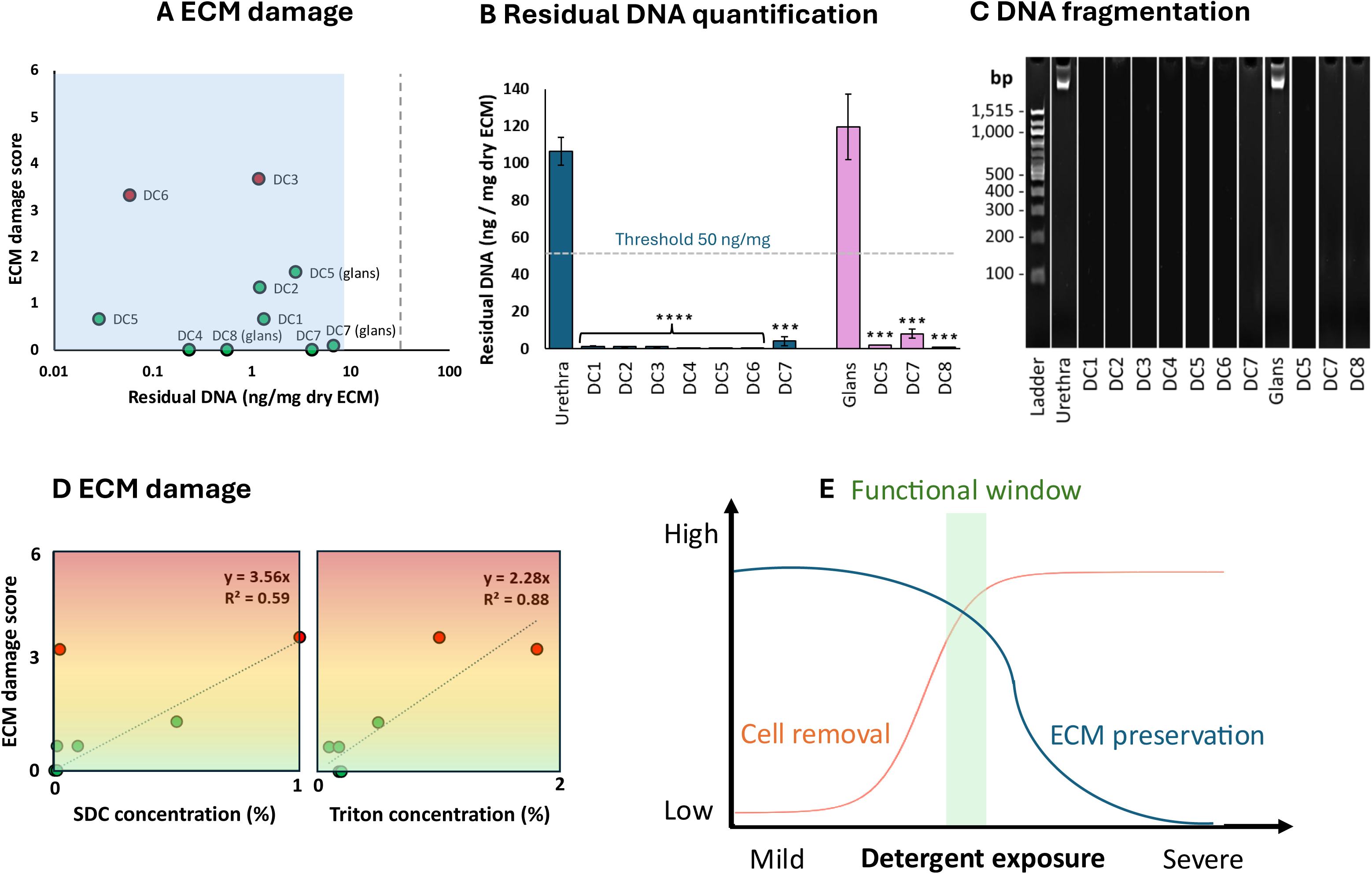
DNA-based decellularization measures fail to predict ECM preservation. (A) Representative overview of iterative decellularization outcomes in human urethral tissue. While all protocols induced effective removal of visible cellular material, ECM preservation varied substantially between conditions. (B) Quantification of residual DNA content across decellularization protocols. Dashed lines indicate accepted thresholds.^23^ (C) DNA fragmentation analysis demonstrating efficient digestion of residual DNA under all conditions.^23^ (D) Histological assessment and quantification of ECM preservation following decellularization. Increasing combined detergent exposure induced progressive ECM disruption despite uniformly effective DNA removal. (E) Concept illustration of the narrow functional window in which efficient cell removal and preservation of ECM integrity overlap. Data are presented as mean ± SD. Statistical analysis was performed using one-way ANOVA with appropriate multiple-comparison correction.

In contrast, preservation of ECM structure varied markedly between protocols. Whereas certain conditions maintained intact tissue organization, others induced pronounced structural deterioration despite similarly effective DNA removal, demonstrating that efficient cellular removal does not necessarily coincide with ECM preservation. Iterative refinement of detergent composition identified a restricted window in which effective cellular removal and preservation of biologically relevant ECM structure could be achieved simultaneously.

Within urethral tissue, intermediate detergent conditions preserved ECM structure while maintaining complete cellular clearance. Deviations towards either excessive or imbalanced detergent exposure resulted in progressive structural loss, indicating that ECM preservation is highly sensitive to coupled detergent effects rather than absolute detergent concentrations alone. Final optimization identified DC7 (0.20% Triton X-100, 0.005% SDC, 200 IU/mL DNase I) as the condition achieving the optimal balance between effective decellularization and structural preservation.

To determine whether this optimal window is transferable across tissues, urethral-optimized conditions were applied to human glans tissue. Despite their effectiveness in urethra, these conditions failed to reproducibly achieve both complete cellular removal and ECM preservation in glans tissue. A modified protocol (DC8) incorporating increased Triton X-100 concentration was required to restore this balance while maintaining structural integrity. Similarly, previously optimized vaginal decellularization conditions (0.18% Triton X-100, 0.015% SDC, 150 IU/mL DNase I) differed from both urethral and glans protocols, further demonstrating that optimal decellularization windows are inherently tissue-specific and likely determined by intrinsic ECM composition and organization.

Together, these findings demonstrate that efficient cellular depletion and preservation of biologically relevant ECM components were achieved within distinct, tissue-specific processing windows. The distinct optimization windows observed across tissues indicate that intrinsic matrix composition and organization are associated with tissue-specific susceptibility to decellularization.

### 2.2 Histological analysis reveals ECM vulnerability governed by coupled detergent interactions and tissue-specific structure

Histological evaluation using Masson’s Trichrome staining revealed substantial variability in ECM preservation across decellularization conditions despite uniformly effective removal of cellular material (Figure 3A–H). Native urethral tissue displayed its characteristic multilayered organization, including stratified epithelium, smooth muscle bundles, vascular structures, and collagen-rich lamina propria (Figure 3A).

**Figure 3.**
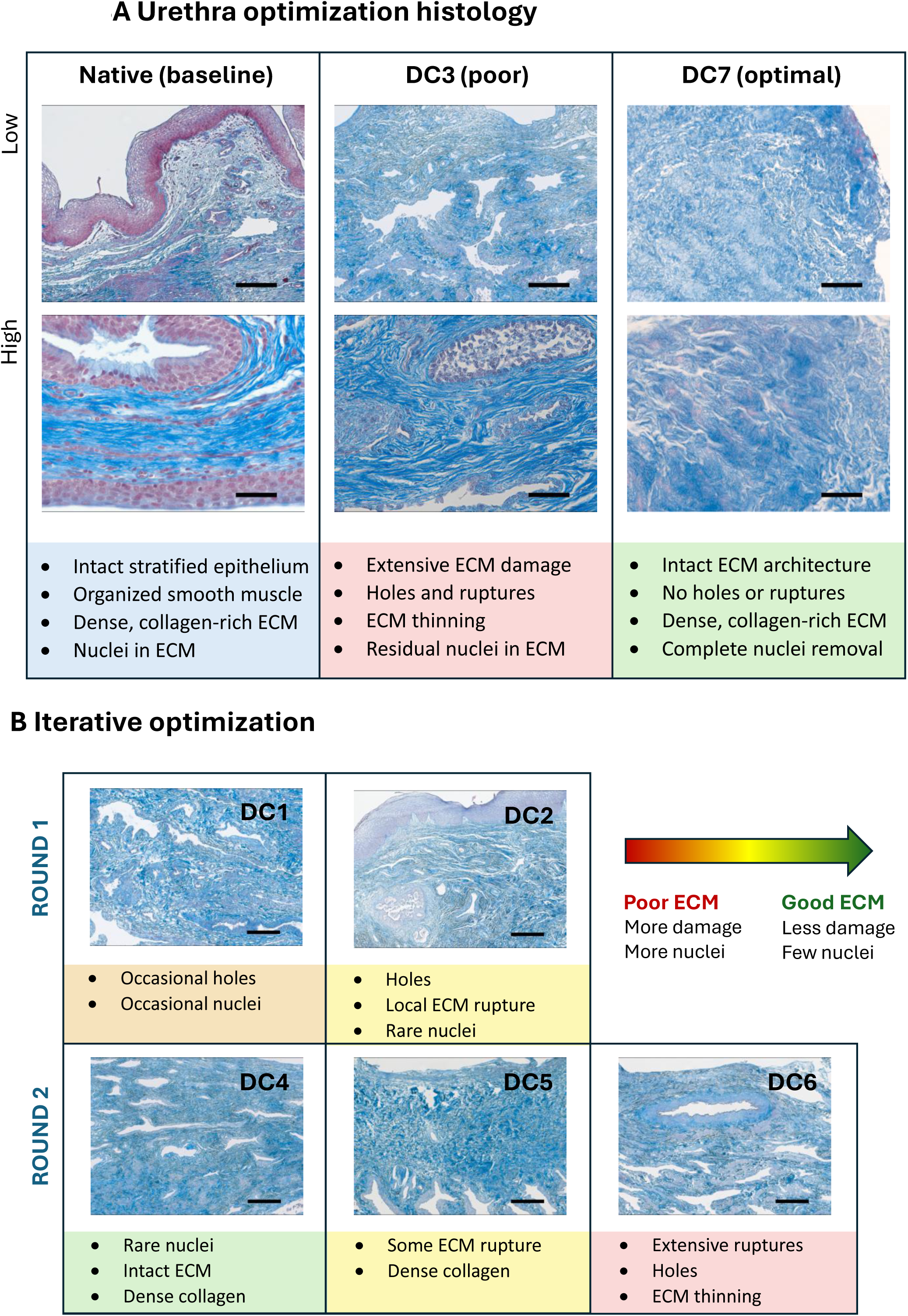
Histological analysis reveals progressive ECM deterioration and optimization-dependent preservation in decellularized urethral tissue. Masson’s trichrome staining of native and decellularized human urethral tissue following iterative optimization of Triton X-100 and sodium deoxycholate concentrations. (A) Representative low- and high-magnification images of native tissue (10x and 40x magnification, respectively), the most disruptive condition (DC3), and the optimized condition (DC7) are shown to illustrate the preserved to severely disrupted ECM architecture. (B) Iterative optimization conditions (DC1, DC2, DC4, DC5, and DC6) demonstrate the progressive effects of detergent composition on matrix preservation. Increasing combined detergent exposure induced ECM thinning, holes, and ruptures, whereas optimized conditions preserved collagen organization and tissue architecture while maintaining effective cellular removal. Scale bars, 200 μm (low magnification, 10x) and 50 μm (high magnification, 40x).

Across all protocols, nuclear material was largely absent, confirming effective decellularization. Residual nuclei were restricted primarily to early optimization conditions (DC1–DC3) and localized around vascular and peripheral tissue regions, frequently in association with residual blood clots, suggesting that incomplete detergent penetration contributes to localized inefficiency. Extended pre-washing steps improved removal of these remnants in subsequent optimization rounds.

Despite comparable cellular clearance, structural preservation was highly protocol-dependent. Importantly, structural damage was not attributable to a single detergent alone but related to the combined interaction between Triton X-100 and SDC. Concentrations of either detergent independently did not entirely predict structural outcome (Figure 2D). Instead, ECM disruption occurred once combined detergent exposure exceeded a tissue-specific tolerance threshold.

In the first optimization regime (DC1–DC3), increasing detergent exposure induced progressive matrix damage characterized by thinning, rupture, and perforation of ECM architecture. In contrast, lower detergent conditions preserved overall tissue organization. During the second optimization regime, reduction of SDC concentrations (0.005–0.025%) substantially improved ECM preservation. DC4 maintained intact structural organization, whereas DC5 induced only localized alterations. However, combining elevated Triton X-100 levels with even minimal SDC exposure (DC6) reintroduced marked ECM disruption, indicating that Triton X-100 impacts tissue susceptibility to SDC-mediated structural damage through coupled chemical interactions rather than independent detergent effects.

Notably, near-complete absence of visible nuclei was achieved across all second-round conditions, demonstrating that reduced detergent exposure does not compromise decellularization efficiency when chemical conditions are appropriately balanced. Final optimization identified DC7 (0.20% Triton X-100, 0.005% SDC, 200 IU/mL DNase I) as the optimal condition. Building on DC4, a modest increase in Triton X-100 concentration was introduced to eliminate the remaining nuclei while maintaining the favorable ECM preservation observed in DC4. This adjustment resulted in complete cellular removal while preserving matrix organization (Figure 3G–H).

Application of urethral-optimized protocols to glans tissue revealed pronounced tissue-specific susceptibility to decellularization (Figure 4B). Conditions that preserved urethral ECM (DC 5 and DC7) induced either incomplete cellular removal or substantial structural disruption in glans tissue. Increasing Triton X-100 concentration (DC8) resolved this trade-off, enabling effective decellularization while maintaining matrix architecture. Notably, previously optimized vaginal conditions resembled DC5, which induced mild-to-severe ECM disruption in both urethral and glans tissue, further supporting that decellularization susceptibility is associated with intrinsic tissue organization and cannot be predicted from detergent parameters alone.

**Figure 4.**
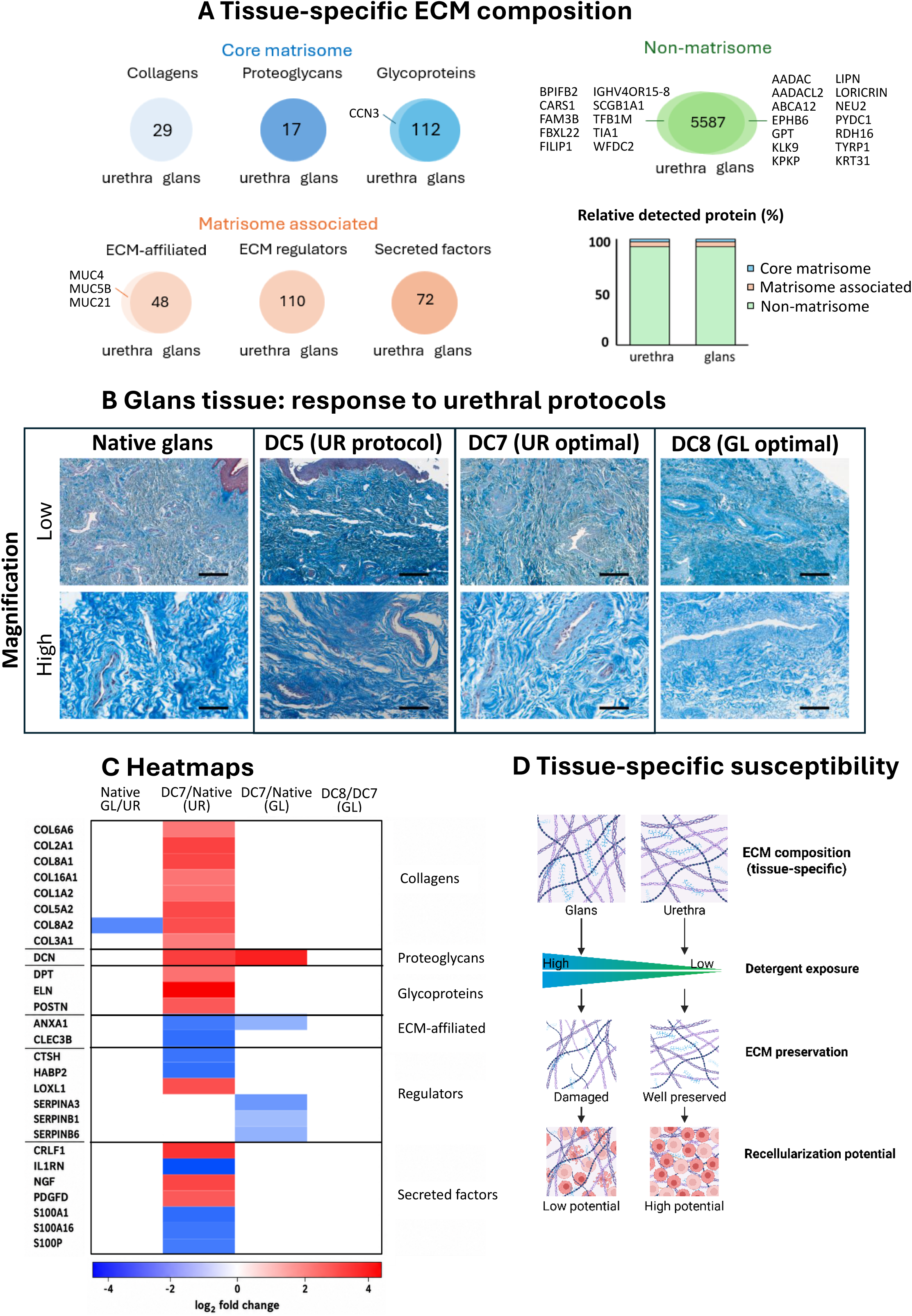
Proteomic analysis identifies tissue-specific ECM preservation patterns following decellularization. (A) Global matrisome profiling of native urethral and glans tissue reveals tissue-specific extracellular matrix composition, including distinct enrichment patterns among glycoproteins, ECM-affiliated proteins, ECM regulators, and non-matrisome proteins. (B) Histological evaluation of glans tissue following application of urethra-derived decellularization protocols demonstrates tissue-specific susceptibility to detergent exposure. Scale bars, 200 and 50 µm. (C) Heatmaps showing relative protein abundance of log₂-transformed quantitative proteomic data across native and decellularization conditions. Colors indicate relative abundance of each protein across experimental groups, with red representing relative abundance, white baseline, and blue relative depletion. (D) Schematic summary illustrating the proposed relationship between tissue-specific ECM composition, susceptibility to chemical processing, ECM preservation following decellularization, and subsequent recellularization characteristics. The schematic represents the conceptual model derived from the experimental observations and is not intended to imply direct causality.

Together, these findings demonstrate that ECM preservation is constrained by tightly coupled detergent interactions and that optimal decellularization conditions differ between tissues with distinct structural organization, indicating that decellularization is a tissue-dependent rather than universally transferable chemical process.

### 2.3 Biochemical assessment confirms decellularization efficiency but fails to predict ECM structural integrity

To quantify decellularization efficiency, residual DNA content and fragmentation were analyzed across all experimental conditions (Figure 2B, Figure S1). Native urethral tissue contained 106.35 ± 21.29 ng DNA per mg tissue.

Across all urethral protocols, residual DNA content was significantly reduced to levels well below accepted decellularization thresholds (<50 ng/mg and <10% of native content).^23^ During the first optimization regime (DC1-DC3), residual DNA levels ranged from 1.19 ± 0.19 ng/mg (1.12%, P < 0.0001) to 1.33 ± 0.28 ng/mg (1.25%, P < 0.0001), with minimal variation between conditions despite substantial differences in ECM preservation observed histologically (Figure 2C).

Optimization during the second regime (DC4-DC6) further reduced DNA content to levels as low as 0.03 ± 0.01 ng/mg (0.1%, P < 0.0001), confirming highly efficient cellular removal across all conditions. Final validation using DC7 yielded 4.07 ± 2.42 ng/mg (3.83%, P<0.001), remaining well within accepted decellularization criteria. Importantly, no relationship was observed between residual DNA levels and preservation of ECM architecture, indicating that biochemical measures of cellular removal do not predict matrix integrity.

DNA fragmentation analysis further demonstrated efficient enzymatic degradation across all protocols. Native tissue contained intact high-molecular-weight DNA fragments exceeding 1,500 bp, whereas all decellularized samples exhibited DNA fragments well below the recommended 200 bp threshold,^23^ with fragment sizes consistently below 100 bp regardless of protocol composition.

Comparable findings were observed in glans tissue. Residual DNA was significantly reduced under all conditions, including DC5 (2.10 ± 1.84 ng/mg, 2.78%, P < 0.001), DC7 (8.04 ± 2.42 ng/mg, 6.73%, P < 0.001), and DC8 (0.67 ± 0.11 ng/mg, 0.56%, P < 0.001). DNA fragments were similarly reduced to ≤75 bp across conditions, again demonstrating uniformly efficient degradation. Notably, DC7 exhibited low residual DNA despite incomplete cellular removal identified histologically, further illustrating the disconnect between biochemical decellularization metrics and true matrix preservation.

Together, these findings demonstrate that DNA-based measurements reliably quantify cellular degradation but provide limited information regarding preservation of ECM architecture and composition. Consequently, biochemical indicators of cellular removal alone are unable to distinguish matrices that retain biologically relevant ECM characteristics from those that have undergone substantial structural damage.

### 2.4 Proteomic analysis identifies tissue-specific ECM composition and preservation patterns following decellularization

Histological analyses demonstrated marked tissue-specific differences in ECM preservation despite comparable cellular removal. Quantitative proteomics was therefore performed to characterize differences in native ECM composition and to determine how protein abundance changed following decellularization of urethral and glans tissue (Figure 4).

Global matrisome analysis demonstrated distinct protein profiles between native urethral and glans tissues (Figure 4A). Although collagens, proteoglycans, ECM regulators and secretion factors were represented in both tissues, differences were observed across glycoproteins, ECM-affiliated proteins, and non-matrisome proteins, indicating distinct molecular composition of the two urogenital ECMs. Native urethral tissue contained proteins associated with mucosal and secretory epithelial functions, including MUC4, MUC5B, MUC21, BPIFB2, and WFDC2, whereas native glans tissue contained proteins associated with epithelial differentiation and barrier formation, including AADAC, AADACL2, ABCA12, EPHB6, GPT, KLK9, KPRP, LIPN, LORICRIN, NEU2, PYDC1, RDH16, TYRP1, and KRT31.

Consistent with these global differences, direct comparison of native tissues identified 15 proteins that differed significantly in abundance between glans and urethral tissue (Figure S2). Seven proteins were more abundant in glans tissue, including COL6A6, CFHR5, ENPP6, KLK14, KRT31, CD36, and RAC3, whereas eight proteins were more abundant in urethral tissue, including ANO1, AGR2, KCNMA1, OLFM4, POPDC2, SLC12A2, TNMD, and KIAA0408. Among these, KRT31, AGR2, TNMD, and KIAA0408 showed particularly pronounced differences between tissues.

The effect of decellularization on protein abundance was assessed within each tissue (Figure 4C). In urethral tissue, optimized decellularization resulted in differential changes in the abundance of ECM-associated proteins with increases in several collagens, proteoglycans, and glycoprotein-associated proteins alongside decreases in several ECM-affiliated proteins and ECM regulators. Several fibrillar collagen proteins, including COL1A1, COL1A2, COL2A1, COL3A1, COL5A1, COL5A2, COL5A3, COL6A1, COL6A2, and COL6A6, showed significantly higher measured abundance in DC7 compared with native urethra. In contrast, several major basement membrane components, including COL4A1, COL4A2, COL4A6, LAMA2, LAMA3, LAMA4, LAMA5, LAMB1, LAMB2, LAMC1, NID1, NID2, and HSPG2, did not differ significantly between native and DC7 urethra. Similarly, the elastic fiber-associated proteins FBLN5, FBN2, ELN and FBN1 showed no significant differences following decellularization. These changes indicate that decellularization did not affect all ECM-associated proteins uniformly but instead produced a selective response across different protein classes. Thus, although selected ECM proteins showed altered abundance following DC7, several major basement membrane and matrix-organizing components remained quantitatively comparable to native tissue, indicating selective rather than uniform changes in ECM protein abundance.

Application of the same urethral-optimized conditions to glans tissue resulted in a more restricted proteomic response. DCN was increased, whereas ANXA1, LOXL1, SERPINA3, SERPINB1, and SERPINB6 were decreased following decellularization. In contrast, major structural ECM components showed no significant changes in abundance following decellularization. These included fibrillar and basement membrane-associated collagens, including COL1A1, COL1A2, COL4A1, COL4A2, COL5A1, COL5A2, COL5A3, COL6A2, and COL6A3, as well as ELN, fibrillin- and fibulin-associated proteins (FBN1, FBN2, FBLN1, FBLN2, and FBLN5), laminin subunits (LAMA2, LAMA3, LAMA4, LAMA5, LAMB1, LAMB2, and LAMC1), and basement membrane-associated proteins including HSPG2, NID1, and NID2. Other ECM components, including TNC, TGFBI, and VCAN, were likewise not significantly altered. Importantly, no significant differences in protein abundance were detected between DC7 and DC8 glans tissue, indicating that the increased Triton X-100 concentration did not measurably alter the glans protein composition under the conditions tested. Thus, although the same chemical processing was applied, the magnitude and composition of the proteomic response differed between urethral and glans tissues.

The tissue-specific differences in decellularization-associated protein changes are summarized in Figure 4D, which illustrates the broader response observed in urethral tissue compared with the more restricted response in glans tissue. Rather than producing a uniform change in protein abundance, chemical processing resulted in tissue-specific alterations in selected ECM-associated and non-ECM proteins, consistent with the tissue-specific differences in matrix preservation observed histologically.

### 2.5 Post-processing sterilization modulates matrix bioactivity independently of ECM architecture

To determine whether post-decellularization processing alters matrix–cell interactions independently of ECM preservation, three sterilization strategies were evaluated: decellularization alone (DC), decellularization followed by antibiotic treatment (DC + AB), and decellularization followed by combined antibiotic and ethanol treatment (DC + AB + EtOH). Previous work demonstrated that DC + AB + EtOH effectively eliminates microbial contaminants in vaginal DMs without disrupting ECM architecture.^37^ Here, the analysis examined whether these post-processing strategies influence early cellular responses in optimized urethral DMs. Sterility assessment demonstrated that decellularization alone was insufficient to eliminate microbial contamination, whereas both DC + AB and DC + AB + EtOH achieved complete sterility over 14 days of culture (Figure 5A).

**Figure 5.**
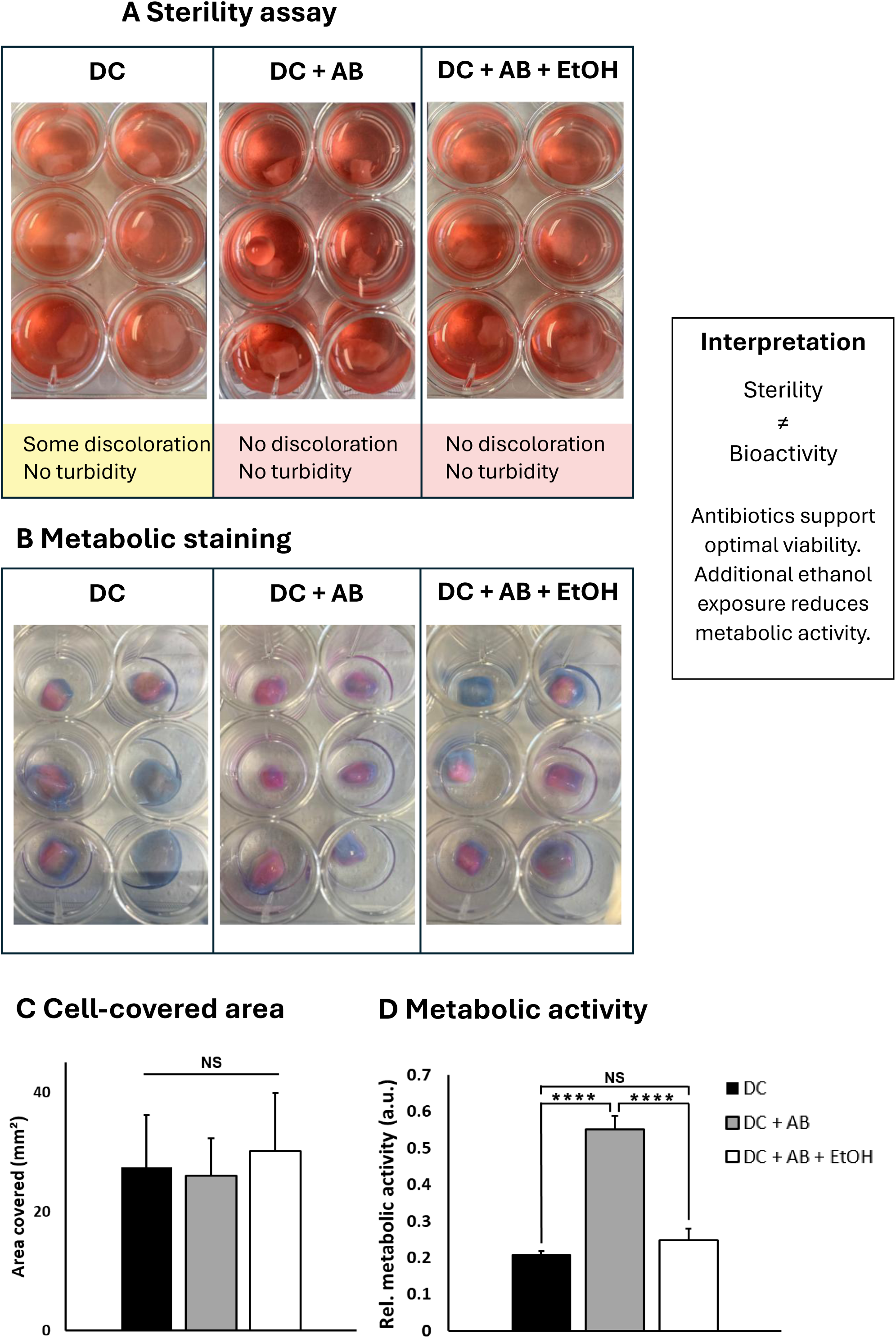
Sterilization modulates matrix bioactivity independently of ECM structural preservation. (A) Sterility assessment at day 14 reveals that post-processing with antibiotics (DC + AB) and antibiotics with ethanol (DC + AB + EtOH) provides sterile urethral matrices, whereas decellularization is insufficient to remove contaminants. (B) Image showing cellular attachment and distribution on sterilized urethral matrices followed by recellularization. (C) Quantification of matrix surface coverage by attached cells across sterilization conditions demonstrates no significant differences. (D) Metabolic activity of recellularized matrices following different sterilization strategies revealed significantly increased activity in DC + AB relative to DC, and DC + AB + EtOH.

Despite equivalent sterility outcomes, marked differences in cellular responses were observed. All DMs supported cellular attachment and surface colonization, with no significant differences in overall cell-covered area (Figure 5B-C). However, metabolic activity differed substantially between processing conditions. DC + AB matrices exhibited significantly increased metabolic activity compared with both untreated DC and ethanol-treated matrices (Figure 5D, *P* < 0.0001). These findings demonstrate that post-processing sterilization can modify matrix bioactivity independently of gross ECM architecture, resulting in distinct cellular responses despite equivalent sterility.

### 2.6 Mechanical and surgical assessment establishes functional preservation of optimized matrices

To determine whether preservation of ECM composition and architecture correlates with preserved mechanically relevant matrix properties, optimized DC7 matrices were subjected to surgical handling assessment, suture retention testing, and uniaxial tensile analysis. Together, these measurements evaluated the capacity of decellularized matrices to preserve mechanically relevant properties required for tissue reconstruction.

Surgical handling characteristics were first quantified using a standardized suturability scoring system. DC7 matrices demonstrated robust intraoperative handling properties, achieving a score of 5.67 ± 0.49, reflecting stable needle passage, knot stability, and resistance to tearing under suture placement. Native urethral tissue exhibited a maximal score of 6 out of 6 and served as the physiological reference standard. These findings indicate preservation of an ECM network capable of dissipating localized mechanical stress during surgical manipulation.

Resistance to localized mechanical loading was subsequently assessed through suture retention testing in both longitudinal and transversal orientations. DC7 matrices exhibited significantly greater retention forces longitudinally (1241 ± 109 gf, P < 0.05) compared with transversally (902 ± 95 gf), demonstrating preservation of anisotropic mechanical behavior consistent with the underlying organized collagen architecture.

Bulk mechanical properties were further characterized using uniaxial tensile testing. DC7 matrices displayed a peak load of 3.91 ± 0.43 N, strain at failure of 340 ± 33%, ultimate tensile strength of 567 ± 83 kPa, and elastic modulus of 5.07 ± 0.96 kPa. Together, these parameters define a matrix that is highly deformable while maintaining substantial load-bearing capacity. This mechanical behavior coincided with retention of fibrillar collagen, basement membrane and elastin-associated ECM proteins identified by quantitative proteomics.

Importantly, these measurements establish the intrinsic mechanical behavior of the optimized decellularized matrix rather than direct replication of native urethral mechanics, for which standardized human reference datasets remain limited. Nevertheless, preservation of anisotropic tensile behavior, load-bearing capacity, tensile stiffness, and suture resistance demonstrates maintenance of functionally relevant ECM architecture beyond histological and biochemical integrity alone.

Together, these findings establish that optimized decellularization preserves mechanically and surgically relevant matrix properties, demonstrating that preservation of ECM architecture is accompanied by retention of properties required for surgical handling and mechanical loading.

**Figure 6.**
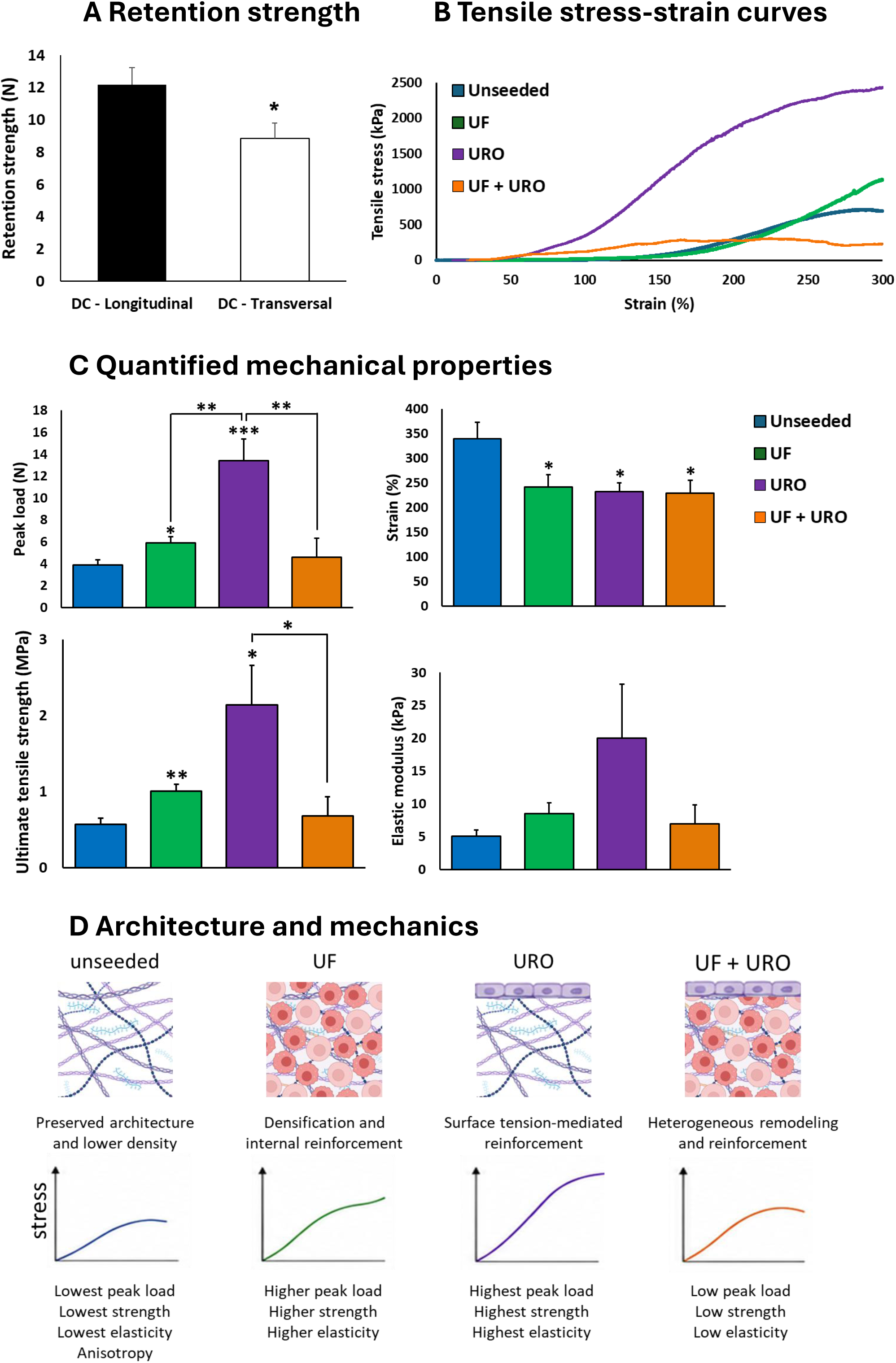
Optimized decellularization preserves mechanically functional ECM architecture and enables cell type-dependent matrix reinforcement. (A) Suture retention testing of optimally decellularized urethral matrices (DC7) in longitudinal and transverse orientations, demonstrating preserved anisotropic mechanical behavior. (B) Representative tensile stress-strain curves of unseeded urethral matrices and after recellularization with urethral fibroblasts (UF), urothelial cells (URO), or combined cultures (UF + URO). (C) Quantification of mechanical properties including peak load, strain at failure, ultimate tensile strength (UTS), and elastic modulus across experimental groups. (D) Preserved ECM architecture and cell type-specific mechanical remodeling. Decellularization preserves aligned collagen and elastin networks and the intrinsic anisotropy but reduces matrix density by cell removal. Fibroblasts reinforce mechanics through volumetric matrix repopulation and densification, whereas urothelial cells primarily strengthen constructs through formation of a tension-bearing surface layer. Combined cultures exhibit heterogeneous remodeling behavior.

### 2.7 Optimized matrices support robust cell attachment, viability and cell-type-dependent expansion

To determine whether optimized urethral DMs exhibiting preserved ECM composition and structure support recellularization, matrices were seeded with primary urothelial cells (URO), urethral fibroblasts (UF), or both cell populations in co-culture using clinically relevant matrix dimensions (Figure 7).

**Figure 7.**
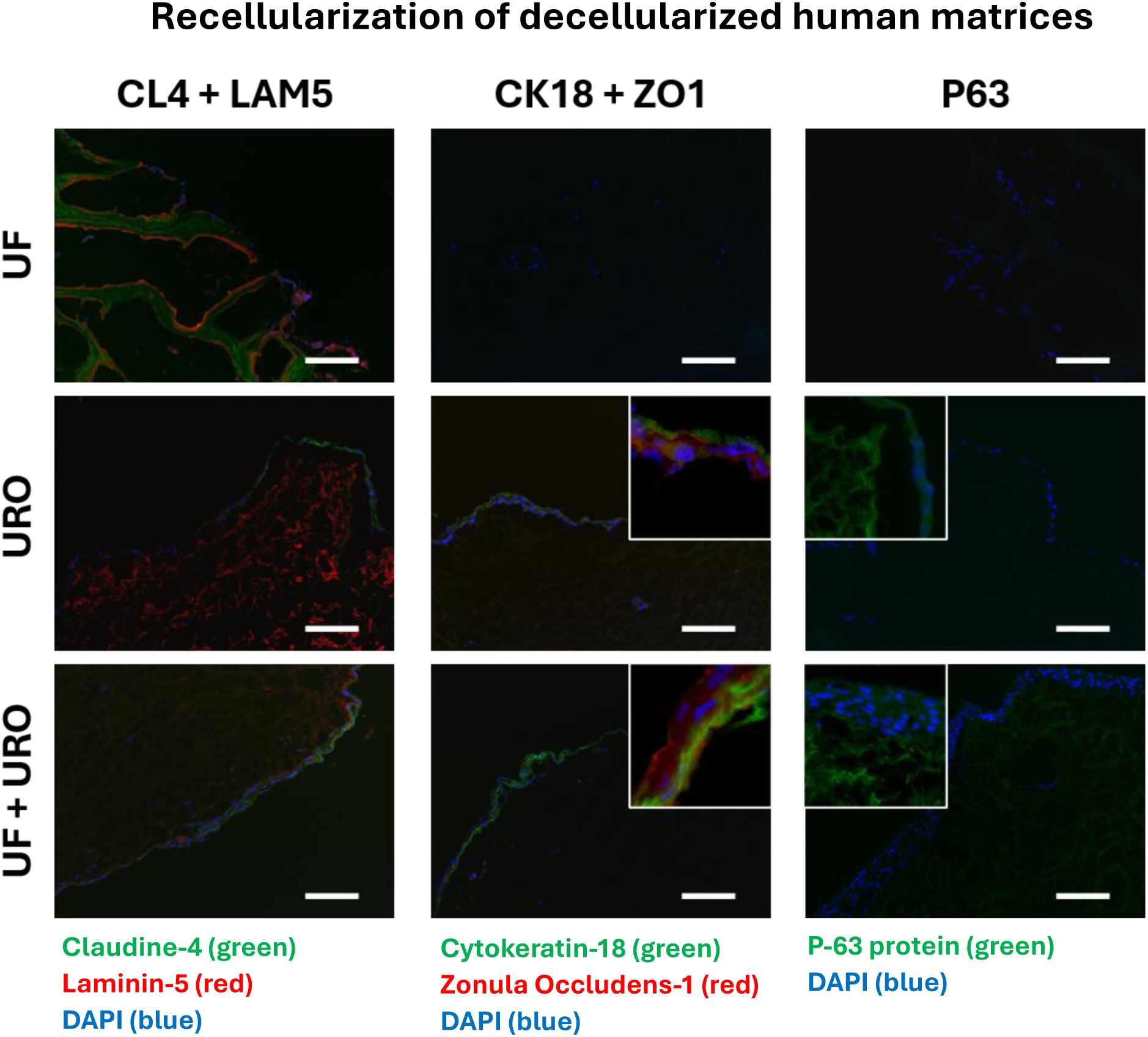
Recellularization of optimized decellularized matrices supports viable, organized, and lineage-appropriate tissue formation. Representative immunofluorescence images of decellularized human urethral matrices after 2.5 weeks of recellularization with urethral fibroblasts (UF), urethral epithelial cells (URO), or a co-culture of fibroblasts and urothelial cells (UF + URO). Claudin-4 (green) and Laminin-5 (red) were used to assess epithelial barrier formation and basement membrane organization (left column), Cytokeratin-18 (green) and Zonula Occludens-1 (red) to evaluate urothelial differentiation and tight junction formation (middle column), and p63 protein (green) to identify basal epithelial cells (right column). Nuclei were counterstained with DAPI (blue). The frames show higher magnification of the epithelial layer. Scale bars = 100 μm.

All recellularized matrices supported rapid cellular attachment and surface colonization throughout the 16-day culture period. Both urothelial cells and fibroblasts exhibited homogeneous surface distribution and spreading across the preserved ECM, indicating that optimized DMs provide a biologically permissive microenvironment for tissue-specific cell populations.

Quantitative analysis revealed distinct cell-type-dependent expansion behavior. Fibroblast-seeded matrices demonstrated extensive cellular expansion, reaching approximately 400% of the initially seeded population by day 16. In contrast, urothelial cells expanded to approximately 60% of the initial seeding density, whereas co-cultured matrices reached approximately 55% of the initial seeding density, indicating distinct cell-type-dependent growth behavior within the preserved ECM.

Metabolic assays confirmed sustained cellular viability throughout the culture period across all experimental groups. These observations are consistent with preservation of ECM composition and structural integrity identified in histological and proteomic analyses.

Together, these findings demonstrate that optimized decellularized matrices support cell attachment, survival, and expansion of primary urethral cell populations, indicating preservation of ECM characteristics compatible with subsequent recellularization.

### 2.8 Optimized matrices support tissue-specific organization and cell-mediated remodeling

To determine whether the preserved extracellular matrix (ECM) architecture retained biologically relevant cues to support tissue-specific cellular organization following recellularization, urethral DMs were cultured with primary urethral fibroblasts, urothelial cells, or a combination of both cell populations for 2.5 weeks and evaluated by immunofluorescence (Figure 7).

Recellularized matrices supported distinct cellular organization depending on the seeding condition. Matrices seeded with urothelial cells displayed epithelial-associated marker expression, including CK18 and Claudin-4, with ZO-1 staining at organized cell-cell junctions, consistent with epithelial organization. Furthermore, p63-positive cells were detected within the urothelial cell-seeded matrices, consistent with the maintenance of a basal epithelial population. Laminin-5 was retained within the ECM in the URO condition, while a pronounced Laminin-5-positive ECM boundary was observed in matrices containing fibroblasts. In the co-culture condition, Laminin-5-associated matrix organization occurred together with epithelial-associated CK18 and Claudin-4 expression, while ZO-1 and p63 staining were observed within the surface-associated cellular layer. These findings indicate that the preserved ECM supported distinct matrix- and cell-associated organization following recellularization.

Notably, this tissue organization occurred within matrices processed under the optimized decellularization conditions that retained basement membrane-associated proteins, mechanically relevant ECM components, and matrix-remodeling-associated factors identified by quantitative proteomics. Together, the immunofluorescence findings demonstrate that the optimized matrices retained matrix-associated features compatible with tissue-specific cellular organization and epithelial-associated characteristics following recellularization.

### 2.9 Recellularization induces cell-type-dependent remodeling of optimized urethral matrices

To determine whether optimized decellularized matrices remain permissive to cell-mediated remodeling following recellularization, mechanically characterized DC7 matrices were repopulated with urethral fibroblasts (UF), urothelial cells (URO), or combined co-cultures (UF + URO), after which the mechanical properties were re-evaluated (Figure 7E-H).

Recellularization altered the mechanical behavior of the matrices in a cell-type-dependent manner relative to unseeded controls (peak load: 3.91 ± 0.43 N, UTS: 567 ± 83 kPa). Urothelial cell-seeded matrices exhibited the largest increases in both peak load (13.41 ± 1.97 N) and UTS (2141 ± 519 kPa), whereas fibroblast-seeded matrices demonstrated more moderate increases (peak load: 5.90 ± 0.57 N, UTS: 1009 ± 86 kPa). In contrast, co-culture conditions did not produce additive improvements in mechanical properties (peak load: 4.58 ± 1.74 N, UTS: 679 ± 257 kPa), indicating that the two cell populations influenced matrix remodeling differently under the present culture conditions.

Across all recellularization groups, strain at failure decreased (UF: 241.81 ± 24.68%; URO: 232.28 ± 17.94%; UF + URO: 228.63 ± 26.15%), while the elastic modulus increased (UF: 8.51 ± 1.68 kPa; URO: 19.98 ± 8.26 kPa; UF + URO: 6.98 ± 2.90 kPa) compared with unseeded matrices. These changes indicate a shift from the highly compliant decellularized matrix toward a mechanically reinforced construct following cellular repopulation.

The observed changes in mechanical properties occurred alongside the cell-type-specific organization described in Section 2.8. Proteomic analysis of DC7 urethral matrices further showed that several major ECM components remained detectable without significant changes in abundance following decellularization, including fibrillar collagens (such as COL1A1 and COL3A1), elastin, fibrillin- and fibulin-associated proteins, and basement membrane-associated components (including COL4A1, COL4A2, HSPG2, NID1, and NID2). In contrast, decellularization was associated with significant changes in the abundance of other ECM-associated proteins, including multiple collagens, proteoglycans, and ECM regulators.

The relationship between ECM composition and the subsequent mechanical response should, however, be interpreted as an association rather than evidence of a direct causal contribution of individual proteins. Quantitative proteomics measures bulk protein abundance and therefore does not directly assess the spatial organization, supramolecular structure, or functional integrity of individual ECM components. Nevertheless, the preservation of multiple major structural ECM components in the optimized DC7 matrices is consistent with their ability to undergo substantial cell-mediated mechanical remodeling following recellularization.

Together, these findings demonstrate that optimized DMs support cell-dependent changes in mechanical properties following recellularization. The magnitude of this remodeling differed between cell populations, with urothelial recellularization producing the greatest increase in peak load and UTS under the conditions tested. Thus, the mechanical response of the optimized matrix depended on the cell population introduced.

## 3. Discussion

Decellularized matrices are intended to retain native ECM architecture and tissue-specific biochemical composition, but their biological performance remains variable across tissues and applications. Current decellularization criteria primarily assess cellular removal through residual DNA content and histological absence of nuclei. While these criteria reduce risk of immunorejection, they provide limited information on preservation of biologically relevant ECM characteristics.^26,39,40,41^ In the present study, systematic optimization combined with quantitative proteomics, structural analyses, mechanical characterization, and recellularization demonstrated that efficient cellular removal does not necessarily coincide with ECM preservation or subsequent biological and mechanical performance.

A central finding of this study is that effective decellularization occurs within a narrow and tissue-dependent operational window. Across a broad range of detergent and enzymatic conditions, residual DNA content and fragment lengths were consistently reduced below accepted thresholds, indicating that cellular removal is comparatively robust. In contrast, ECM preservation was highly sensitive to relatively small changes in chemical composition. Structural damage did not correlate linearly with detergent concentration but instead emerged from coupled interactions between Triton X-100 and sodium deoxycholate, suggesting that ECM disruption is governed by threshold-dependent chemical interactions rather than the isolated effects of an individual chemical. Moreover, identical chemical conditions produced different structural outcomes in urethral and glans tissues, demonstrating that optimal decellularization conditions are tissue dependent rather than universally transferable. Numerous studies have similarly reported the need for tissue-specific optimization across cardiovascular, pulmonary, hepatic and urogenital tissues.^28,42–56^ These findings emphasize that decellularization should be optimized according to the susceptibility of the target tissue to matrix disruption rather than cellular removal alone.

Quantitative proteomics demonstrated that native urethral and glans tissues have distinct molecular compositions, with differences among glycoproteins, ECM-affiliated proteins, ECM regulators, and non-matrisome-associated proteins. Direct comparison of the native tissues further identified 15 proteins with significantly different abundance between glans and urethra. Seven proteins, including COL6A6, CFHR5, ENPP6, KLK14, KRT31, CD36, and RAC3, were more abundant in glans tissue, whereas eight proteins, including ANO1, AGR2, KCNMA1, OLFM4, POPDC2, SLC12A2, TNMD, and KIAA0408, were more abundant in urethral tissue. These differences included both ECM-associated proteins, such as COL6A6, and proteins associated with epithelial, membrane, enzymatic, and other cellular functions, indicating that the molecular distinction between the tissues extends beyond the ECM alone.

Following decellularization, the proteomic response also differed between tissues. In optimized urethral matrices, quantitative proteomics revealed differential changes in the abundance of collagen-, proteoglycan-, and glycoprotein-associated proteins, together with changes in ECM-affiliated proteins and ECM regulators. In glans tissue, the observed changes were more limited, including increased DCN and decreased ANXA1, LOXL1, SERPINA3, SERPINB1, and SERPINB6. Thus, decellularization altered selected ECM-associated proteins rather than producing a uniform change across the matrix. The distinct proteomic responses observed under identical processing conditions further support the need to consider tissue-specific molecular composition when optimizing decellularization.

These findings are consistent with previous tissue-engineering studies demonstrating that tissue-specific ECM microenvironments contribute to tissue-specific cellular behavior.^9,57^ For example, Orabi et al. showed that replacement of dermal fibroblasts with bladder-derived stromal cells improved urothelial differentiation and biomechanical maturation in self-assembled urinary bladder substitutes, emphasizing that organ-specific extracellular environments influence tissue development beyond simple structural support.^57^ Together with the present findings, these observations support intrinsic tissue composition as an important determinant of biomaterial performance. Previous proteomic studies have primarily provided inventories of ECM composition following decellularization, whereas relatively few have directly compared intrinsic tissue composition with preservation following identical chemical processing.^58^ The present comparative approach therefore extends proteomic analysis beyond descriptive characterization by relating intrinsic tissue composition to tissue-specific responses to identical decellularization conditions.

Basement membrane proteins, fibrillar collagens, elastin-associated proteins, and matrix remodeling-associated ECM components have been implicated in epithelial attachment, tissue compartmentalization, mechanical integrity and matrix homeostasis.^59,60^ These functions are consistent with the established role of the ECM in regulating tissue organization and regeneration rather than serving solely as a structural scaffold.^10,24,61^ Preservation of basement membrane components has furthermore been associated with improved epithelial regeneration in decellularized pulmonary and renal scaffolds.^62–65^ In the current study, several major structural ECM components, including fibrillar collagens, elastin, fibrillin- and fibulin-associated proteins, and basement membrane-associated proteins, remained detectable in optimized DC7 urethral matrices without significant changes in abundance. Their retention, together with preservation of tissue architecture, was accompanied by tissue-specific cellular organization and cell-dependent mechanical remodeling following recellularization.

The retained matrix also supported tissue-specific organization following recellularization, including epithelial-associated marker expression and maintenance of matrix-associated laminin organization. Recellularization induced cell-dependent mechanical remodeling, with greater mechanical reinforcement following urothelial than fibroblast recellularization. Although fibroblasts are generally considered important regulators of ECM synthesis and remodeling,^24^ the greater mechanical reinforcement following urothelial recellularization could reflect differences in cellular organization, matrix contraction, ECM turnover, or other lineage-specific interactions with the matrix. Cell-mediated remodeling of biological scaffolds depends on coordinated interactions among cellular contractility, ECM turnover and matrix organization, which can differ between cell populations and scaffold composition.^66,67^ These findings show that optimized DMs remain responsive to cellular repopulation and that the mechanical response is dependent on the cell population introduced.

Post-processing sterilization also influenced cellular metabolic activity despite similar preservation of ECM architecture. Although both sterilization strategies achieved complete sterility, antibiotic treatment alone resulted in higher metabolic activity than matrices subsequently exposed to ethanol. These findings suggest that processing steps after decellularization can modulate matrix bioactivity independently of gross structural preservation, consistent with previous reports that sterilization can alter cellular responses despite minimal detectable changes in matrix architecture.^40,68,69^ Thus downstream processing represents an additional factor that can influence the biological performance of decellularized matrices.

By integrating quantitative proteomics with structural, mechanical, and biological analyses, the present study demonstrates that DNA removal and ECM preservation represent complementary aspects of decellularization quality. Quantitative proteomics provides molecular information that complements conventional biochemical and histological assessment by identifying both preserved structural ECM components and selective changes in protein abundance. Together with the recellularization findings, these results support tissue-specific decellularization strategies that evaluate ECM composition, structure, and function alongside cellular removal.

## Conclusions

Successful decellularization is not defined solely by efficient cellular removal but also by preservation of biologically relevant ECM composition and architecture. Across human urogenital tissues, effective decellularization occurs within a narrow, tissue-specific operational window in which extensive cellular depletion and preservation of key ECM characteristics can be achieved simultaneously. Quantitative proteomic analysis identified tissue-specific differences in intrinsic protein composition, including differences in ECM-associated and non-ECM proteins, that provided a molecular context for the distinct responses to chemical processing observed between tissues. Importantly, preserved ECM composition was associated with tissue-specific cellular organization and cell-dependent mechanical remodeling following recellularization. Together, these findings establish a composition-informed approach for evaluating decellularized matrices and provide a foundation for rational, tissue-specific biomaterial design.

## Supplements

Supporting Information is available on the journal’s webpage.

## Acknowledgements

The authors would like to thank Manon Labrecque, Todd Galbraith, and Sébastien Larochelle for their support with experimental procedures, and Robert de Leeuw for software support.

## CRediT author statement

S.B.: conceptualization, investigation, resources, supervision, project administration, funding acquisition, writing – review & editing. Y.S.: investigation, writing – review & editing. V.F.: Methodology, formal analysis, resources, data curation, writing – review & editing. F.R.-D.: software, validation, resources, supervision, writing – review & editing. A.D.: software, resources, writing – review & editing, supervision. S.C.: writing – review & editing. J.J.S.: Conceptualization, methodology, validation, formal analysis, investigation, data curation, writing – original draft, visualization, supervision, project administration, funding acquisition.

All authors have read and agreed to the published version of the manuscript.

## 4. STAR Methods

### 4.1 Key resource table

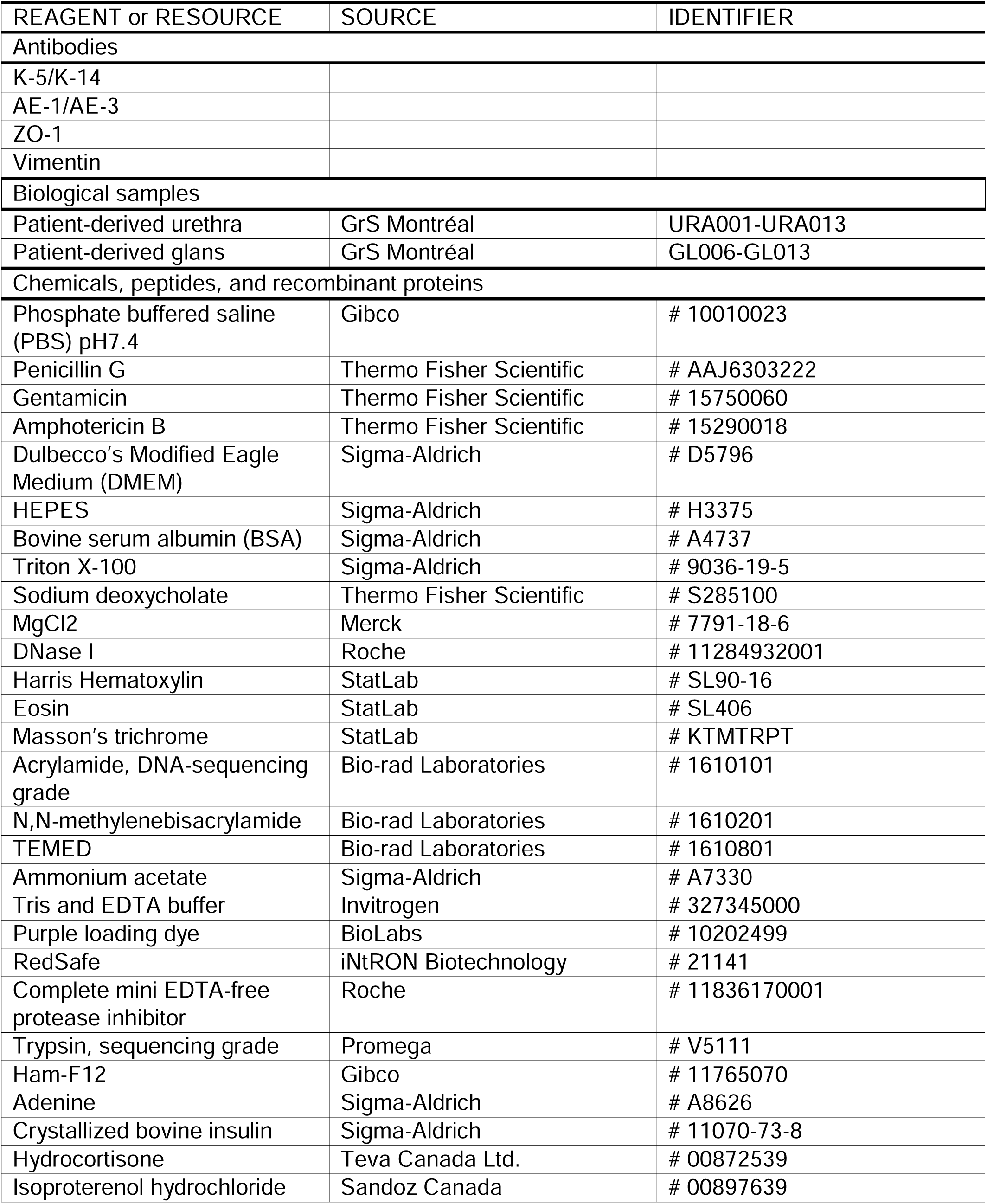
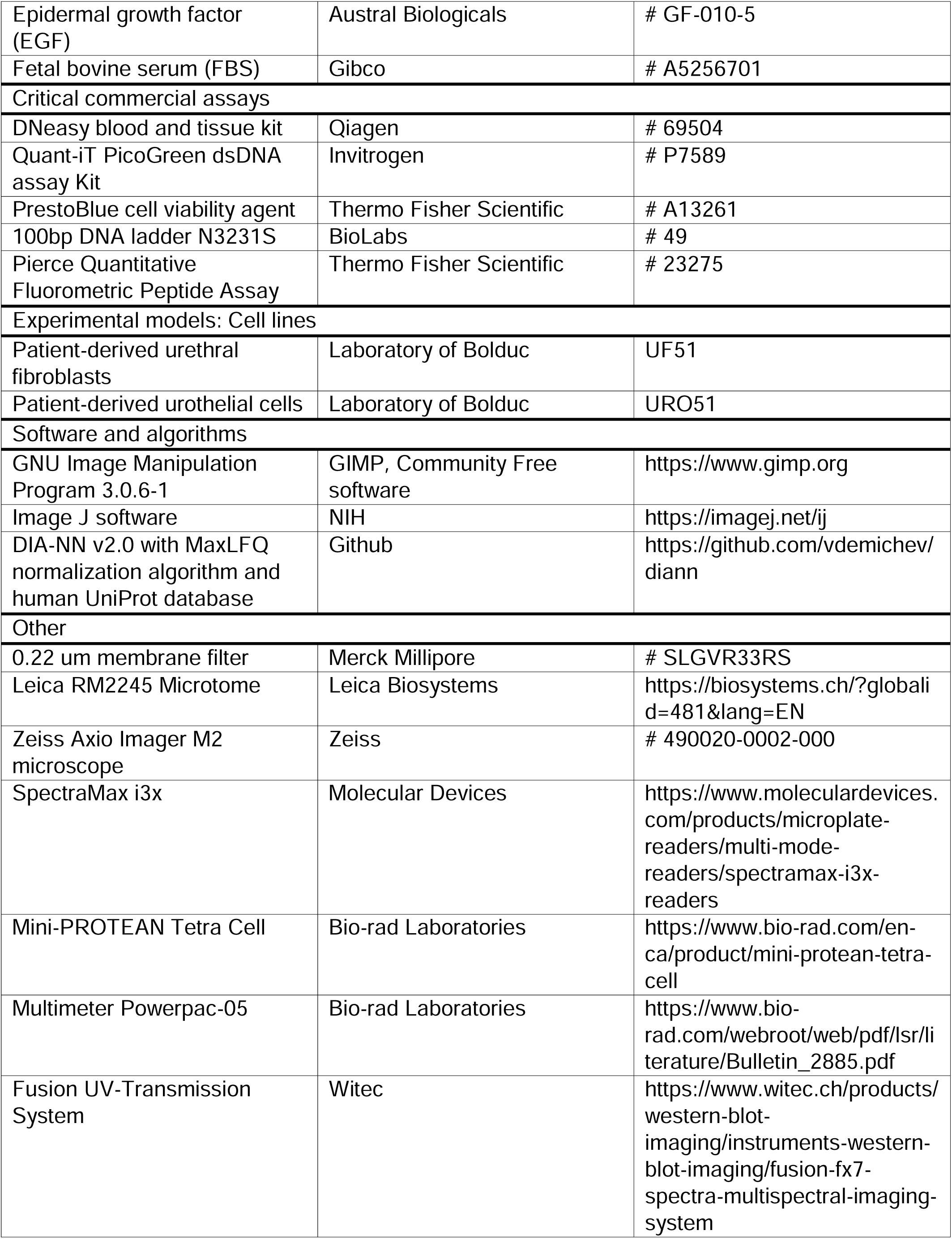
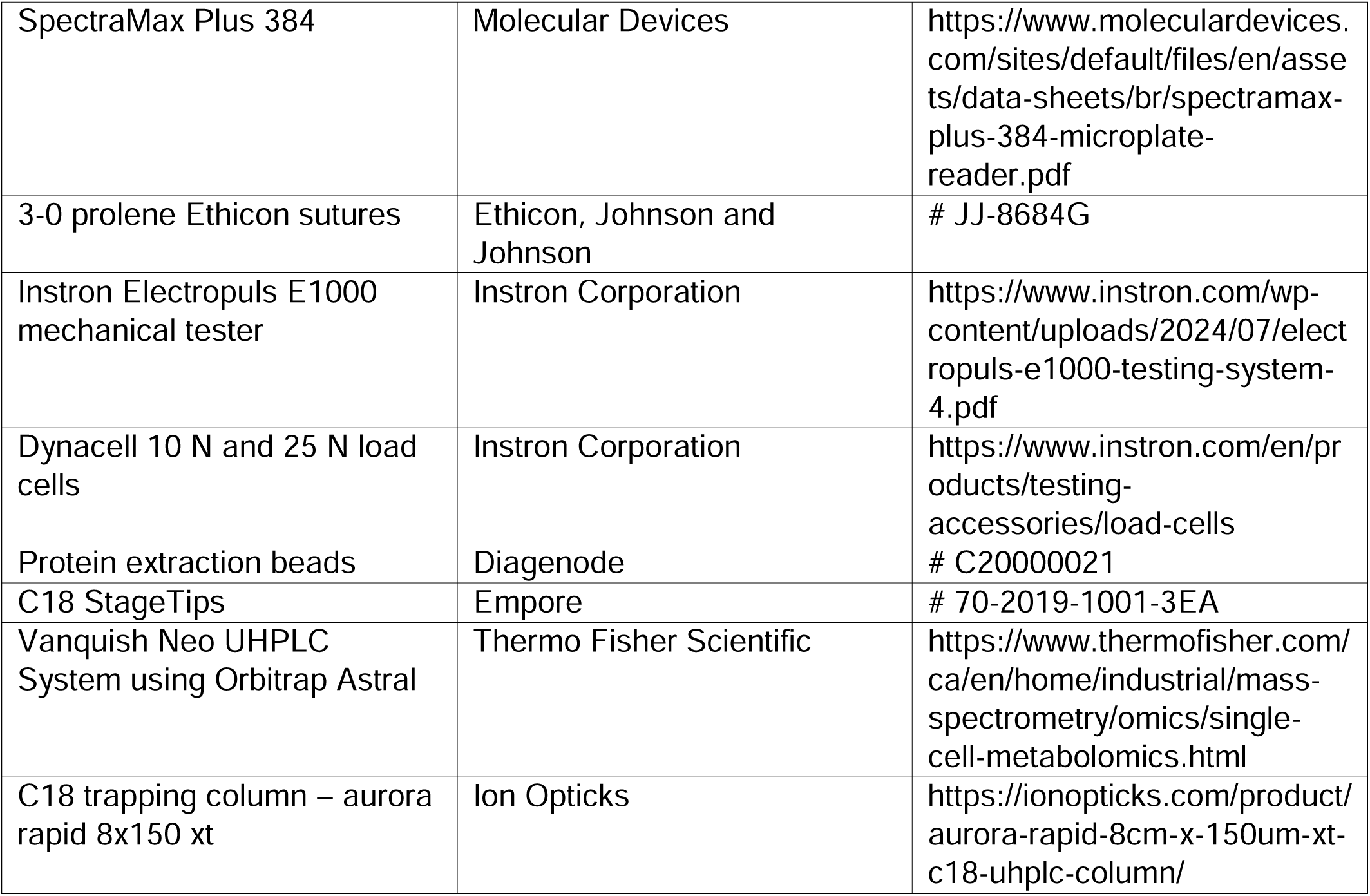

### 4.2 Resource availability

#### LEAD CONTACT AND MATERIALS AVAILABILITY

This study did not generate new unique reagents. Further information for resources should be directed to and will be fulfilled by the lead contact Stéphane Bolduc.

#### EXPERIMENTAL MODEL AND STUDY PARTICIPANT DETAILS

##### Human subjects

Human urethral and glans tissue was obtained from 13 healthy donors during gender-affirming vaginoplasty at GrS Montréal (Montreal, Canada) between 2025 and 2026. All donors were assigned male at birth and identified as transfeminine. Donor age ranged from 18–50 years and all donors provided written informed consent. The study was approved by the Comité d’éthique de la recherche du CHU de Québec–Université Laval (protocol DR-002-1190) and conducted in accordance with the Declaration of Helsinki.

#### METHOD DETAILS

##### Tissue retrieval and processing

Resected penile tissue containing 8 cm of urethra, corpus spongiosum, partial glans, and segments of corpus cavernosum was immediately rinsed three times in PBS (Gibco) supplemented with 100 U/mL penicillin (Thermo Fisher Scientific), 25 μg/mL gentamicin (Thermo Fisher Scientific), and 0.5 μg/mL amphotericin B (Thermo Fisher Scientific) for 30-60 s each by manual shaking. Tissues were stored on ice in DMEM (Sigma-Aldrich) supplemented with 25 mM HEPES (Sigma-Aldrich), 0.15% BSA (Sigma-Aldrich) and 2.5 μg/mL amphotericin B and processed within 4–7 h post-collection. In a biosecurity hood, tissues were longitudinally opened along the middle of the ventral surface and cut into full-thickness, full-width urethral strips (3.2 ± 0.6 cm).

##### Decellularization of urethral tissue

Urethral tissues were decellularized using seven detergent-based protocols (Table 1) derived from a previously established vaginal tissue protocol.^36^ Samples were treated with Triton X-100 (Sigma-Aldrich), sodium deoxycholate (Thermo Fisher Scientific), and DNase I (Roche) under constant agitation (70 RPM, orbital shaker). The protocol started with a PBS wash containing 100 U/mL penicillin, 25 μg/mL gentamicin, and 0.5 μg/mL amphotericin B (24 h, RT), followed by incubation in PBS with Triton X-100 and sodium deoxycholate (24 h, 37 °C). After new PBS washes (2 × 20 min, RT) incubation in PBS with DNase I and 23.4 mmol MgCl_2_ (Merck; 24 h, 37 °C) was performed. Lastly, final PBS washes (2 x 20 min, RT) were performed. The total protocol was 4 days.

##### Processing of glans tissue

Glans tissue was dissected from the same donor specimens, cut into halves and subjected to identical decellularization protocols. Glans samples were used for histological ECM integrity scoring, DNA quantification, DNA fragment length assays and quantitative proteomic analysis. Structural ECM integrity was assessed histologically, and glans samples were included in comparative proteomic analysis. No mechanical or suturability testing was performed on glans samples.

##### Sterilization treatments

Decellularized matrices were sterilized using three testing conditions: 1) decellularization using the optimal procedure for urethra, 2) decellularization followed by a high-concentration antibiotics treatment (3 h, 4 °C) of 1,000 U/mL penicillin, 250 μg/mL gentamicin, and 5 μg/mL amphotericin B in PBS, followed by PBS washes (2 x 20 min, 4 °C), and 3) decellularization followed by high-concentration antibiotic and 70% ethanol treatment (5 min, 4 °C), and PBS washes (2 x 20 min, 4 °C). All solutions were sterilized before use with a 0.22 μm membrane filter (Merck Millipore).

##### Histology and ECM structural assessment

Native and decellularized samples from each donor were fixed in 3.7% formaldehyde (overnight, RT) and washed in PBS (2 x 20 min, RT) and 70% ethanol (3 x 30 min, RT) under constant agitation (70 RPM, orbital shaker). Samples were paraffin-embedded and sectioned to 6 μm on a Leica RM2245 microtome (Leica) with a 6° knife angle. Slides were stained with Masson’s trichrome (MT; StatLab) using deparaffinization, rehydration, staining and mounting. Images captured on a Zeiss Axio Imager M2 microscope (Zeiss) with AxioCam ICc1 camera (Leica) were processed in GNU Image Manipulation software (GIMP).

##### ECM integrity scoring (blinded analysis)

Structural ECM integrity was evaluated using a semi-quantitative blinded scoring system (0–6 scale per parameter) of holes without lumen, structural disruption and thinning of the matrix. Scores were averaged across 20 ROIs per section for two sections.

##### DNA quantification and fragment analysis

DNA was extracted from native and decellularized samples below 25 mg in weight using DNeasy Blood & Tissue Kit (Qiagen). Extracted DNA was diluted in 200 µl of the kit-supplied buffer AE. DNA quantification was performed using Quant-iT PicoGreen dsDNA Assay Kit (Invitrogen) with 485 nm excitation and 520 nm fluorescence detection on a SpectraMax i3x (Molecular Devices). Quantities were normalized to dry ECM weight.

The residual fragment length was assessed by 6% polyacrylamide gel electrophoresis following precipitation with 2M ammonium acetate (Sigma-Aldrich) and 70% ethanol. For this, samples were incubated 30-60 min on ice and centrifuged (10 min at 15,000 RPM, 0 °C). Supernatant was removed, 750 µl of 70% ice-cold ethanol was added and centrifugation was repeated (2 min at 15,000 RPM, 0°C). Supernatant was removed, ethanol evaporated and residual DNA was left to dissolve overnight in 50 µl of 10 mM tris and 1 mM EDTA buffer (Invitrogen). Electrophoresis was performed (30 min, constant 120 V) using a Multimeter Powerpac-05 (Bio-rad). After 30 min RedSafe staining (iNtRON Biotechnology), images were acquired and processed using a Fusion UV-Transmission System (Witec).

##### Sterility testing

Scaffolds were incubated in antibiotic-free medium with phenol-red for 14 days at 37 °C. Media turbidity and color change were monitored daily as indicators of infection, in accordance to previously established methods.^70^

##### Cytotoxicity and growth inhibition assay

To determine potential matrix-induced inhibiting effects, a growth inhibition test was performed. A 24-well plate was preheated (10 min, 37 °C) with 1 mL DMEM/well. Decellularized urethra blocks of 0.3-0.5 cm^2^ in size were sterilized according to the three testing conditions, seeded with 4 x 10^4^ human urethral fibroblasts (uFBs, passage 3) and incubated in 750 µl DMEM with 10% v/v fetal bovine serum (FBS, Gibco), 100 U/mL penicillin, 25 μg/mL gentamicin, and 0.5 μg/mL amphotericin B for 14 days at 37 °C in 8% CO_2_.

A 40-min staining with 750 µl 10% PrestoBlue Cell Viability Agent (Thermo Fischer Scientific) in PBS was used to assess metabolic activity at days 7 and 14, and to confirm live cell coverage. Medium without cells served as background control. Digital photographs were obtained and the cell growth area was quantified using Image J software (NIH). The PrestoBlue absorbance was measured at 570 nm (sample measurement) and 600 nm (background reference) using a 96-well plate on a SpectraMax Plus 384 (Molecular Devices). The blank subtraction method was performed.

##### Suturability assessment

Surgical handling was evaluated using 6-0 Prolene sutures (Ethicon, Johnson and Johnson) that were placed by an experienced urologist. A 7-point Likert scale (0-very poor, 1-poor, 2-fair, 3-average, 4-good, 5-very good and 6-excellent) was used to score suture placement quality for three sutures/sample in both lateral and transversal orientation. Native urethra served as reference.

##### Mechanical testing Tensile testing

Dog-bone-shaped specimens were cut using a homemade stainless-steel punch (Figure Sx) and tensile tested using an Instron E1000 system (Instron Corporation) with a Dynacell 10 N load cell (accuracy of ±0.0005 N). Starting with an initial gauge length of 3 mm between grips, specimens were stretched at 1 mm/s until failure. Cross-sections were defined by the width of the punch (2 mm) and the full thickness, which was measured using a laser millimeter. Data were exported to Excel 365 (Microsoft Office). The Peak load F_max_ was determined as the maximum strength (N) before failure. The ultimate tensile strength σ_ult_ (UTS) was calculated using the peak load and tissue cross section: σ_ult_ = F_max_ / (tissue width x thickness). The strain was expressed as the length gain (%) relative to the initial gauge length and the elastic modulus (E) was calculated using the slope of the strain curve at 20% and 60% of the UTS: *E* = 0.4 x σult/(ε60%σ - ε20%σ).^34,35^

##### Suture retention testing

Suture retention tests^36,37^ were performed on urethral samples of 10-15 mm in length. Sutures of 3-0 Prolene (Ethicon, Johnson and Johnson) were placed 2 mm away from the edge and 5 mm apart in both lateral and transversal orientation. Samples were vertically fixed in an Instron E1000 system with Dynacell 25 N load cell (accuracy of ±0.0025 N) and sutures were pulled at 50 mm/min. The suture retention strength was defined as the maximum force F_max_ (gf) before failure.

##### Proteomic analysis

###### Protein extraction and digestion

Native and decellularized samples of 50-80 mg in weight were placed in 1.5mL sonication compatible TPX tubes (Bioruptor) with 60-80 mg sonication beads (Diagenode) and a 200 µL solution of 8M Urea. Tissue was solubilized by sonication with a Bioruptor (15×30 sec at high intensity and at 4°C, Diagenode). Proteins were precipitated by mixing with five volumes of cold acetone and incubating overnight at −20°C. Protein denaturation was done by heating at 95 °C for 5 min, reduction of cysteine disulfide bonds with 4 µL dithiothreitol (500 mM at 37 °C for 2h at 1400 RPM) and cysteine alkylation with 4 µL iodoacetamide (500 mM at RT for 30 min at 1400 RPM). Urea was diluted by addition of 600 µL 100 mM ammonium bicarbonate. For protein digestion, samples were incubated 2h at 37 °C with 1000 units PNGaseF at 1400 RPM, 2h at 37 °C with 1 µg LysC at 1400 RPM, and overnight at 37 °C with 1.5 µg trypsin (sequencing grade, Promega). Samples were incubated 2h with 1.5 µg trypsin at 37 °C. Samples were acidified to a pH of 2.5 with 12 µL 50% trifluoroacetic acid and evaporated under vacuum. Tryptic peptides were desalted on C18 StageTips (Empore) and vacuum-dried. Samples were solubilized with 0.1% FA and adjusted at a peptide concentration of 0.050 µg/µL using fluorescence measurements (Pierce Quantitative Fluorometric Peptide Assay method, Thermo Fisher Scientific) and 0.25 µg was transferred into vials.

##### Mass spectrometry

Peptides were analyzed on a Vanquish Neo UHPLC System using an Orbitrap Astral (Thermo Fisher Scientific). Chromatography was operated in trap and elute mode using a C18 trapping column (PepMap Neo Trap Cartridge, Thermo Fisher Scientific) and a C18 separation column (Aurora Rapid 8×150 XT C18, Ion Opticks). Peptides were eluted 15 min using 5-34% of linear gradient solvent B (B: 100% acetonitrile in 0.1% formic acid) with flow rate of 1µL/min followed by a column wash at 85% B at 2uL/min for 2min for a total runtime of 17 minutes. Spray voltage was set to 1800V and heated capillary temperature at 280°C. All data were acquired in profile mode using positive polarity and lock mass internal calibration with the EASY-IC setting. Using a data independent acquisition (DIA) method, precursors were acquired in the Orbitrap analyzer at 180,000 resolutions on a 380-980 m/z mass range, with 500% AGC target and 5 ms maximum injection time. For MS2, precursors were isolated using a 380-980 m/z mass range with 150 windows of 4 m/z and fragmented using 25 % HCD collision energy with Astral analyzer acquisition on a 150 to 1,500 m/z mass range. For fragment ions, a 500% AGC target with 5 ms maximal injection time were selected.

##### Data analysis

Spectra were analyzed with Spectronaut (version 20, Biognosys) using an in-silico digested Homo sapiens protein sequence database (UniProt Reference Proteome – Proteome ID UP000005640 – 83,339 entries – 2025.02) to perform a directDIA search. The following parameters were set at: Trypsin/P as enzyme parameter, a maximum of 2 missed cleavage, Carbamido-methylation as fixed modificationand methionine N-terminal excision and oxidation, Proline and Lysine oxidation and Glutamine conversion in pyroglutamate were variable modifications with a maximum of 5 allowed per peptide. Imputation strategy was set to Run Wise Imputing and LFQ method to MaxLFQ.Only 2+ to 4+ precursors were considered for a 350-980 m/z mass range, and fragments on a 100-2,000 m/z range. MBR (match between runs) option was enabled. A protein was considered for quantification if it presented at least 2 identified peptides and if there was an intensity value in at least 75% of the replicates in one of the two groups. Missing values were imputed from the 1^st^ percentile intensity value for each sample independently. Significance of the abundance variation was determined from the proteins that presented a limma (FDR < 0.05, |z| > 1.96). Candidate proteins were selected based on a paired Welch t-test and z-score thresholds.

##### Cell seeding of urethral matrices

To test functionalization, the DMs were placed in a 48-well plate (Nunc, Thermo Fisher Scientific) and seeded with human urethral fibroblasts (UFs) and/or urothelial cells (UCs). The UFs and UCs were isolated from human urethral biopsies, as previously described.^38^ Biopsies were obtained from healthy patients during reconstructive surgeries for benign conditions. All biopsies came from male donors (one donor for each cell type). Cells were used between passage 1 and 3. UFs were seeded at 2 x 10^5^ cells/mL and UCs at 1 x 10^6^ cells/mL. Different seeding densities were applied to prevent overgrowth by UFs. First, either 20 µl of cell suspension was added (UFs on the bottom and UCs on the epithelial side to ensure epithelial–stromal organization) and incubated for 30 min to facilitate adherence. DMs were immersed by adding another 0.73 mL cell suspension and incubated at 37 °C in 8% CO_2_. UFs and UCs were cultured in UC medium containing a 3:1 mix of DMEM and Ham-F12 (GE Healthcare), 24.3 µg/mL adenine (Sigma-Aldrich), 5 µg/mL crystallized bovine insulin (Sigma-Aldrich), 1.1 µM hydrocortisone (Teva Canada Ltd.), 0.212 µg/mL isoproterenol hydrochloride (Sandoz Canada), 10 ng/mL epidermal growth factor (Austral Biologicals), 100 U/mL penicillin and 25 µg/mL gentamicin. Cells were seeded x days on urethral matrices with media changes performed three times a week.

##### Quantification and statistical analysis

Decellularized urethral matrices were developed from 13 patients to perform each experiment with biological (n) and/or technical replicates (N). Microsoft Office Excel 2019 (Microsoft Corporation) was used for statistical analyses. Pre-testing of decellularized matrices before colonization involved sterility testing, and cytotoxicity tests on 6 technical replicates (n = 1, N = 6). The growth inhibition test involved triplicates of these 6 samples (N = 3 x 6). Extensive evaluation of the DMs using DNA quantification and fragment length analysis, were based on 8 readings (n = 4, N = 2). Histological analysis involved 8 sections (n=4, N = 2) that were each investigated at 20 regions of interest (ROIs). Suture placement and the suture retention test were performed with 9 sutures per condition (n = 3, N = 3), and the tensile stretch test was performed on 7 technical replicates (n = 1, N = 7). Proteomics was performed on 4 specimens (n = 4, N = 1). Any significant difference between conditions was tested using a two-way ANOVA test (assuming equal variance) after confirmation of normal distributions by a Shapiro-Wilk test. Sample sizes were based on a priori power analysis for a one-way ANOVA with eight independent groups (α = 0.05), providing a 80-90% power to detect an effect size of Cohen’s f = 0.6 (large, ISO 10993-5:2009 predicts a 30-50% reduction) for a recommended 6-9 observations per group. Post-hoc Dunn’s tests with Bonferroni α corrections were performed. Data analysis was performed in Graph Pad Prism 10.2.3 (Graph Pad software). Data was plotted as mean with standard error (SER). Statistical significance was illustrated with: * for P ≤ 0.05, ** for P ≤ 0.01, *** for P ≤ 0.001 and **** for P ≤ 0.0001.

##### Immunofluorescence staining

Immunofluorescence on OCT sections was performed on cryo-sectioned slides stored at −20 °C. Prior to staining, an appropriate volume of PBS 1× supplemented with CaCl□ (0.131 g/L) and MgCl□ (0.17 mL/L of 2.8 M stock solution) was prepared in DI water under stirring. PBS 10X stock was added to reach 1X final concentration, and the solution was adjusted to a working volume of approximately 200 mL per histology bath for approximately 24 slides.

Cryo-sections were removed from −20 °C storage and, when required, additional slides were included for positive and negative controls. The slides were fixed in pre-cooled acetone (−20 °C) in a histology bath for 10 min at −20 °C. Following fixation, slides were washed three times for 2 min in PBS 1X supplemented with CaCl□ and MgCl□, using alternating wash baths to prevent drying of the tissue. From this step onward, sections were kept continuously hydrated to minimize background staining.

For immunostaining, all samples were fixed in a 3.7% neutral buffered formaldehyde solution, washed 1x in PBS and 3x in 70% ethanol for each 30 min and embedded in paraffin. Histological sections, 5 mm thick, were stained. Sections were permeabilized with 0.2% Triton X-100 in PBS for 30 min, followed by blocking with 1% bovine serum albumin (BSA) in PBS for 30 min at room temperature to reduce nonspecific antibody binding. Samples were subsequently incubated for 45 min at 25°C with primary antibodies diluted in blocking buffer. The following primary antibodies were used: anti-Laminin-5 (rabbit, Abcam, Ab14509, 1:100), anti-Claudin-4 (mouse, Invitrogen, #32-9400; 1:100), anti-Cytokeratin-18 (CK18) (mouse, ARP, #03-61009; 1:100), anti-ZO-1 (rabbit, Invitrogen, #40.2200; 1:50), and anti-p63 (mouse, Abcam, Ab124762; 1:100).

Following three washes with PBS, sections were incubated for 30 min at 25°C in the dark with AF488 donkey anti-mouse (Invitrogen, A-21203; 1:100) and AF594 donkey anti-rabbit (Invitrogen, A-21206; 1:100), using Hoechst as counterstain. After a final series of PBS washes, sections were mounted and left to dry at RT for at least 48 h before microscopic analysis.

Fluorescence images were acquired using a Nikon Eclipse E600 epifluorescence microscope (Nikon, Mississauga, ON, Canada) with 10/20x objective under identical acquisition settings for all samples within each staining. Scale bars were added with GIMP 3.2.4 (GNU Image Manipulation Program, version 3.2.4). At least two tissue sections of each experimental group were analyzed.

