## Supplements for "Extracellular matrix composition is associated with tissue-specific decellularization susceptibility and mechanical remodeling across human urogenital tissues"

***Table S1: Histology ratings of ECM damage.*** *Assessment of visible DNA, and holes, rupture and thinning of the ECM for decellularization protocol 1-7 using a 7-point Likert scale (0-never, 1-rarely, 2-sometimes, 3-in general, 4-often, 5-usually, 6-always).*

| Protocol urethra | Visible DNA | ECM holes | ECM rupture | ECM thinning |
|---|---|---|---|---|
| DC1 | 2 | 2 | 0 | 0 |
| DC2 | 1 | 3 | 1 | 0 |
| DC3 | 2 | 4 | 3 | 4 |
| DC4 | 1 | 0 | 0 | 0 |
| DC5 | 0 | 0 | 2 | 0 |
| DC6 | 0 | 2 | 5 | 3 |
| DC7 | 0 | 0 | 0 | 0 |
| Protocol glans | Visible DNA | ECM holes | ECM rupture | ECM thinning |
| DC5 | 2 | 2 | 3 | 0 |
| DC7 | 3 | 0 | 0 | 0 |
| DC8 | 0 | 0 | 0 | 0 |

***Figure S2: Differential protein abundance between native urethral and glans tissues.*** *Heatmap showing proteins that were differentially abundant between native urethral and glans tissues based on quantitative proteomic analysis. Values represent $\log_2$ fold changes for glans relative to urethra. Red indicates higher protein abundance in glans tissue, whereas blue indicates higher protein abundance in urethral tissue. Color intensity reflects the magnitude of the $\log_2$ fold change. All proteins shown met the significance threshold of adjusted $p < 0.05$.*

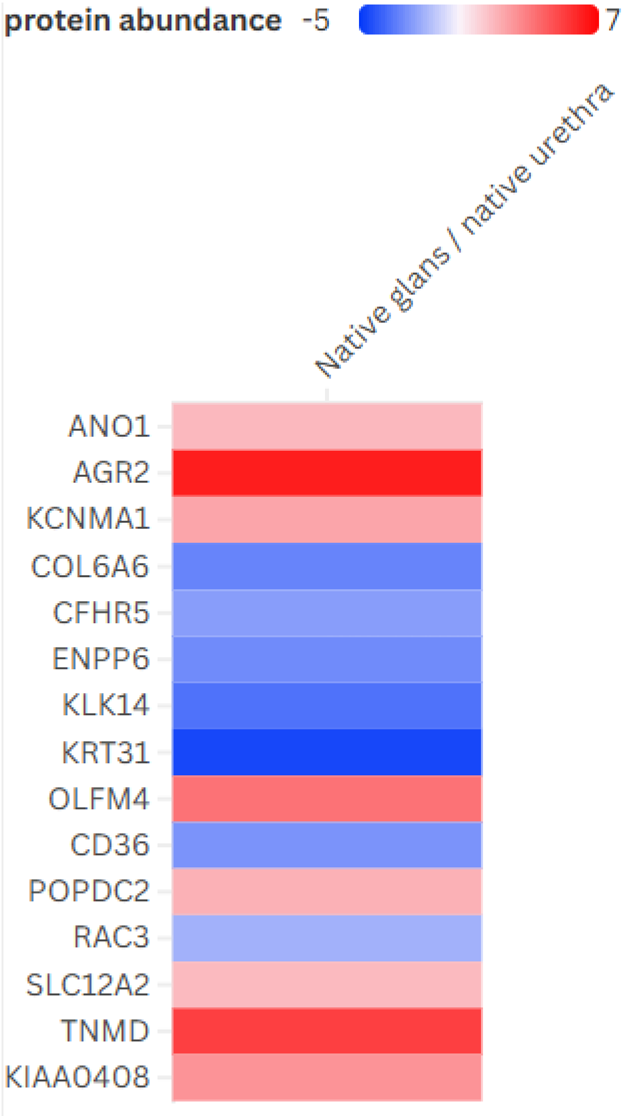

***Figure S3: Metal stamp for preparation of bone-shaped matrix specimens for mechanical testing.*** *Photograph of the custom metal stamp used to prepare bone-shaped specimens for mechanical characterization. The stamp was used to cut standardized bone-shaped specimens from the matrix for subsequent uniaxial tensile testing. The resulting geometry enabled consistent specimen dimensions and mechanical loading across samples.*

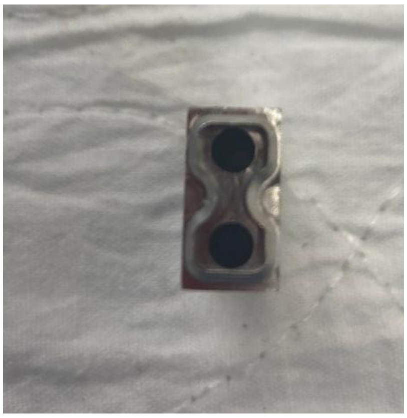

***Table S4: List of identified genes in native and decellularized tissue.*** *List of all identified proteins in native urethra (6982 proteins).*

36951
;;HBG1
;;SRCAP
;AHR
;ALG9
;BBS5
;BCKDHA
;COX15
;EXOSC4
;GDPGP1;GD
PGP1
;HIGD1A;HIG
D1A
;IRF6
;KLC1;KLC1;
KLC1;KLC1;K
LC1
;KLKB1
;LRRC57;LRR
C57
;MRPL2
;MRPL23;MR
PL23;MRPL2
3;MRPL23
;MYCBP
;OC90
;PARL
;POLA2;POL
A2;POLA2;P
OLA2
;QRICH1
;RPS17
;SCGN
;SHPK
;TGM3
;VKORC1;;VK
ORC1
;ZNF653
A1BG
A2M
A2ML1
A4GALT
AAAS
AACS
AAGAB
AAK1
AAMDC
AAMP
AAR2
AARS1
AARS2
AASDHPPT
AASS
ABAT
ABCA1
ABCA2
ABCA6
ABCA8
ABCB1
ABCB10
ABCB7
ABCC1
ABCC4
ABCC9
ABCD1
ABCD3
ABCD4
ABCE1
ABCE1
ABCF1
ABCF2
ABCF3
ABCG2
ABHD10
ABHD11
ABHD12
ABHD14B
ABHD16A
ABHD17A;AB
HD17A;ABH
D17A;ABHD1
7C;ABHD17A
ABHD17B
ABHD4
ABHD5
ABHD6
ABI1
ABI2
ABI3
ABI3BP
ABLIM1
ABLIM2
ABLIM3
ABR
ABRACL
ABRAXAS2
ACAA1
ACAA2
ACACA
ACAD8
ACAD9
ACADM
ACADS
ACADSB
ACADVL
ACAN
ACAP2
ACAT1
ACAT1
ACAT2
ACBD3
ACBD5
ACBD6
ACE
ACER1
ACHE
ACIN1
ACKR1
ACLY
ACO1
ACO2
ACO2
ACOT1
ACOT11
ACOT13
ACOT2
ACOT7
ACOT9
ACOX1
ACOX2
ACOX3
ACP1
ACP2
ACP3
ACP5
ACSF2
ACSF3
ACSL1
ACSL3
ACSL4
ACSL5
ACSS1
ACSS2
ACSS3
ACTA1
ACTA2
ACTB
ACTB
ACTB
ACTC1
ACTG1
ACTG2
ACTL6A
ACTN1
ACTN1
ACTN1
ACTN2
ACTN4
ACTR10
ACTR1A
ACTR1B
ACTR2
ACTR3
ACTR3B
ACVR1
ACVRL1
ACY1;ABHD1
4A-
ACY1;ACY1
ACYP1
ACYP2
ADA
ADA2
ADAM10
ADAM15
ADAM9
ADAMTS17
ADAMTS4
ADAMTS5
ADAMTSL1
ADAMTSL3
ADAMTSL4
ADAMTSL5
ADAR
ADAR
ADARB1
ADCY2
ADCY5
ADCY7
ADD1
ADD3
ADGRA2
ADGRE2
ADGRE5
ADGRL2
ADGRL3
ADGRV1
ADH1B
ADH1C
ADH5
ADH7
ADI1
ADIPOQ
ADIRF
ADISSP
ADK
ADNP
ADO
ADPGK
ADPGK
ADPRH
ADPRS
ADRM1
ADSL
ADSS1
ADSS2
AEBP1
AFAP1
AFAP1L2
AFDN
AFG2A
AFG2B
AFG3L2
AFM
AFTPH

AGAP1
AGAP3
AGFG1
AGFG2
AGK
AGL
AGO1
AGO2
AGO3
AGPAT1
AGPAT3
AGPAT4
AGPAT5
AGPS
AGR2
AGR3
AGRN
AGRN
AGT
AGTPBP1
AGTRAP
AHCTF1
AHCY
AHCYL1
AHCYL2
AHNAK
AHNAK2
AHSA1
AHSG
AHSP
AIDA
AIF1
AIF1L
AIFM1
AIG1
AIMP1
AIMP2
AIP
AIRIM
AJUBA
AK1
AK2
AK3
AK4
AK6
AKAP1
AKAP10
AKAP12
AKAP13
AKAP8
AKAP8L
AKR1A1
AKR1B1
AKR1B10
AKR1C1
AKR1C2
AKR1C3
AKR7A2
AKT1
AKT1S1
AKT2
AKT3
AKTIP
ALAD
ALB
ALB
ALCAM
ALDH16A1
ALDH18A1
ALDH1A1
ALDH1A2
ALDH1A3
ALDH1B1
ALDH1L1
ALDH1L2
ALDH2
ALDH3A1
ALDH3A2
ALDH3B1
ALDH4A1
ALDH5A1
ALDH6A1
ALDH7A1
ALDH9A1
ALDOA
ALDOC
ALG1
ALG11
ALG12
ALG13
ALG2
ALG3
ALG5
ALG6
ALG8
ALKAL2
ALKBH2
ALKBH5
ALOX12
ALOX12B
ALOX15B
ALOX5
ALOX5AP
ALOXE3
ALYREF
AMBP
AMDHD2
AMN1
AMOTL1
AMPD2
AMPD3
AMPH
AMT;AMT;AMT;;AMT;AMT;AMT;AMT
AMZ2
ANAPC1
ANAPC13
ANAPC4
ANAPC5
ANAPC7
ANG
ANGEL1
ANGPTL2
ANGPTL4
ANGPTL5
ANGPTL6
ANGPTL7
ANK1
ANK2
ANK2
ANK3
ANKFY1
ANKHD1
ANKMY2
ANKRD13A
ANKRD22
ANKRD26
ANKRD28
ANKRD35
ANKRD44
ANKS1A
ANO1
ANO10
ANO6
ANP32A
ANP32B
ANP32E
ANPEP
ANTXR1
ANTXR2
ANTXR2
ANXA1
ANXA11
ANXA2
ANXA3
ANXA4
ANXA5
ANXA6
ANXA6
ANXA7
ANXA8
ANXA8L1
ANXA9
AOC3
AOX1
AP1B1
AP1G1
AP1G2
AP1M1
AP1M2
AP1S1
AP1S2
AP1S3
AP2A1
AP2A2
AP2B1
AP2M1
AP2S1
AP3B1
AP3D1
AP3M1
AP3M2
AP3S1
AP4B1
AP4S1
AP5B1
AP5M1
AP5Z1
APAF1
APBB1
APCS
APEH
APEX1
API5
APIP
APMAP
APOA1
APOA2
APOA4
APOB
APOBEC3A
APOBEC3B
APOBEC3C
APOBEC3G
APOC1
APOC3
APOC4-
APOC2
APOD
APOE
APOF
APOH
APOH
APOL1
APOL2
APOL3
APOL5
APOM
APOO
APOOL
APP
APPL1
APPL2
APRT

APTX
AQP1
AQP3
AQR
AR
ARAF
ARAP1
ARAP3
ARCN1
AREL1
ARF1
ARF4
ARF5
ARF6
ARFGAP1
ARFGAP2
ARFGAP3
ARFGEF1
ARFGEF2
ARFIP1
ARFRP1
ARG1
ARHGAP1
ARHGAP10
ARHGAP12
ARHGAP17
ARHGAP18
ARHGAP21
ARHGAP23
ARHGAP25
ARHGAP26
ARHGAP27
ARHGAP31
ARHGAP32
ARHGAP35
ARHGAP4
ARHGAP45
ARHGAP5
ARHGAP6
ARHGDIA
ARHGDIB
ARHGEF1
ARHGEF10
ARHGEF10L
ARHGEF11
ARHGEF12
ARHGEF16
ARHGEF17
ARHGEF18
ARHGEF2
ARHGEF25
ARHGEF37
ARHGEF5
ARHGEF6
ARHGEF7
ARHGEF7
ARID1A
ARID1B
ARID2
ARIH1
ARL1
ARL15
ARL2
ARL2BP
ARL3
ARL5A
ARL6IP4
ARL6IP5
ARL6IP6
ARL8A
ARL8B
ARLN
ARMC1
ARMC10
ARMC5
ARMC6
ARMC8
ARMCX1
ARMCX2
ARMCX3
ARMH3
ARMT1
ARNT
ARPC1A
ARPC1B
ARPC2
ARPC3
ARPC4-
TTLL3;ARPC4
ARPC5
ARPC5L
ARPP19
ARRB1
ARRB2
ARRDC1
ARSA
ARSB
ARSF
ARVCF
ASAH1
ASAP1
ASAP2
ASAP3
ASB3
ASCC1
ASCC2
ASCC3
ASGR2
ASH2L
ASL
ASMTL
ASNS
ASPA
ASPH
ASPM
ASPN
ASPRV1
ASPSCR1
ASS1
ASXL2
ATAD1
ATAD3A
ATAD3A
ATE1
ATG101
ATG16L1
ATG16L2
ATG2A
ATG2B
ATG3
ATG4B
ATG5
ATG7
ATG9A
ATG9B
ATIC
ATL1
ATL2
ATL2
ATL3
ATM
ATOX1
ATP11B
ATP11C
ATP13A1
ATP13A4
ATP1A1
ATP1A2
ATP1A3;;ATP
1A3;ATP1A3
ATP1B1
ATP1B1
ATP1B2
ATP1B3
ATP2A2
ATP2A3
ATP2B1
ATP2B4
ATP2B4
ATP2C1
ATP5F1A
ATP5F1A
ATP5F1B
ATP5F1C
ATP5F1D
ATP5F1E;ATP
5F1EP2
ATP5IF1
ATP5ME
ATP5MF
ATP5MF-
PTCD1
ATP5MG
ATP5MJ
ATP5MK
ATP5PB
ATP5PD
ATP5PF
ATP5PO
ATP6AP1
ATP6AP2
ATP6V0A1
ATP6V0A2
ATP6V0D1
ATP6V0D2
ATP6V0E1
ATP6V1A
ATP6V1B2
ATP6V1C1
ATP6V1D
ATP6V1E1
ATP6V1F
ATP6V1G1
ATP6V1H
ATP7A
ATP8A1
ATP8B1
ATP9A
ATPAF1
ATPAF2
ATRN
ATXN10
ATXN1L
ATXN2
ATXN2L
ATXN3
ATXN7L3B
AUH
AUP1
AVEN
AVIL
AVPR1A
AXL
AZGP1
AZU1
B2M
B3GALT5
B3GALT6
B3GAT3
B3GLCT
B4GALT1
BABAM1
BABAM2
BACC1
BAD

BAG1
BAG2
BAG3
BAG5
BAG6
BAIAP2
BAIAP2
BAIAP2L1
BAK1
BANF1
BAP1
BASP1
BAX
BAZ1B
BBOX1
BBS1
BBS2
BBS7
BBS9
BCAM
BCAP31
BCAS1
BCAS2
BCAS3
BCAT1
BCAT2
BCCIP
BCHE
BCKDHB
BCKDK
BCL10
BCL2
BCL2L1
BCL2L13
BCL2L2-
PABPN1
BCL9L
BCLAF1
BCR
BCS1L
BDH1
BDH2
BDNF
BECN1
BET1
BGN
BICD2
BICDL2
BIN1
BIN3
BIRC2
BIRC6
BLMH
BLOC1S1
BLOC1S2
BLOC1S3
BLOC1S5-
TXNDC5;BLO
C1S5
BLOC1S6
BLTP2
BLVRA
BLVRB
BMP4
BMPER
BMS1
BNIP1
BNIP2
BNIPL
BOD1L1
BOK
BOLA2
BOP1
BORCS5
BORCS6
BORCS7-
ASMT;BORCS
7
BPGM
BPHL
BPIFB1
BPIFB2
BPNT1
BPNT2
BRAF
BRAT1
BRCC3
BRD1
BRD3
BRD4
BRD8
BRI3
BRI3BP
BRIX1
BRK1
BROX
BSG
BST1
BST2
BTAF1
BTBD8
BTBD9
BTD
BTF3
BTF3L4
BTK
BTN2A1
BTN3A2
BTN3A3
BTRC;FBXW1
1;FBXW11;BT
RC
BUB3
BUD31
BYSL
BZW1
BZW2
C11orf54
C11orf58;C1
1orf58;SMAP
C11orf68
C11orf96
C11orf98
C12orf57
C12orf75;OC
C1
C15orf38-
AP3S2;ARPIN
C17orf58
C19orf12
C19orf25
C1orf198
C1QA
C1QB
C1QBP
C1QC
C1QTNF12
C1QTNF2
C1QTNF3-
AMACR;C1Q
TNF3
C1QTNF5
C1QTNF7
C1R
C1R
C1RL
C1S
C2
C2CD2L
C2CD5
C3
C3orf38
C4A
C4B
C4BPA
C4BPB
C5
C5orf46
C6
C6orf132
C6orf136
C6orf47
C6orf89
C7
C7orf50
C8A
C8B
C8G
C8orf33
C8orf58
C8orf82
C9
CA1
CA12
CA2
CA3
CA4
CA5B
CAB39
CAB39L
CACHD1
CACNA1C
CACNA2D1
CACNA2D2
CACNB2
CACYBP
CAD
CADM1
CADM3
CADM4
CADPS
CALB1
CALB2
CALCA
CALCOCO1
CALCOCO2
CALCRL
CALD1
CALD1
CALHM2
CALHM5
CALM1;CAL
M2;CALM3
CALML3
CALML5
CALR
CALU
CAMK1
CAMK1D
CAMK2D
CAMK2D
CAMK2G
CAMKK1
CAMKK2
CAND1
CAND2
CANX
CAP1
CAP2
CAPG
CAPN1
CAPN2
CAPN5
CAPN7
CAPNS1

| | | | | |
|---|---|---|---|---|
| CAPNS2 | CBX1 | CCT8 | CDC42BPA | CELF2 |
| CAPRIN1 | CBX3 | CCZ1B;CCZ1 | CDC42BPB | CELSR1 |
| CAPS | CBX5 | CD109 | CDC42BPG | CELSR2 |
| CAPZA1 | CBX7 | CD14 | CDC42EP1 | CENPE |
| CAPZA2 | CC2D1A | CD151 | CDC42EP4 | CENPV |
| CAPZB | CC2D1B | CD163 | CDC5L | CEP162 |
| CARD16 | CCAR1 | CD200 | CDC73 | CEP192 |
| CARD18 | CCAR2 | CD207 | CDCP1 | CEP250 |
| CARD19 | CCDC102B | CD209 | CDH1 | CEP43;CEP4 |
| CARHSP1 | CCDC115 | CD248 | CDH11 | 3;CEP43;;CE |
| CARM1 | CCDC124 | CD276 | CDH13 | P43;CEP43;; |
| CARMIL1 | CCDC127 | CD2AP | CDH19 | CEP43;CEP4 |
| CARNMT1 | CCDC175 | CD2BP2 | CDH2 | 3;CEP43;CEP |
| CARS1 | CCDC22 | CD302 | CDH23 | 43;;CEP43;C |
| CARS1 | CCDC25 | CD34 | CDH5 | EP43;CEP43; |
| CARS2 | CCDC43 | CD36 | CDIPT | CEP43;CEP4 |
| CASK | CCDC47 | CD38 | CDK1 | 3 |
| CASKIN2 | CCDC50 | CD4 | CDK10 | CEP44 |
| CASP1 | CCDC6 | CD40 | CDK11B | CEP76 |
| CASP10 | CCDC61 | CD44 | CDK12 | CEPT1 |
| CASP14 | CCDC65 | CD46 | CDK16 | CERCAM |
| CASP3 | CCDC80 | CD47 | CDK17 | CERS2 |
| CASP4 | CCDC88A | CD48 | CDK2 | CERS3 |
| CASP6 | CCDC9 | CD55 | CDK2AP1 | CERT1 |
| CASP7 | CCDC90B | CD58 | CDK4 | CES1 |
| CASP8 | CCDC91 | CD59;;CD59 | CDK5 | CES2 |
| CASQ2 | CCDC93 | CD5L | CDK5RAP3 | CETN2 |
| CAST | CCL14 | CD63 | CDK6 | CFAP20 |
| CAST | CCL21 | CD74 | CDK9 | CFAP298- |
| CASTOR1 | CCL28 | CD81 | CDKN1B | TCP10L;CFA |
| CASZ1 | CCM2 | CD82 | CDKN2A;CD | P298- |
| CAT | CCN2 | CD83 | KN2A;CDKN2 | TCP10L;CFA |
| CAV1 | CCN3 | CD8A | A;CDKN2A;C | P298- |
| CAV1 | CCN5 | CD9 | DKN2B | TCP10L;CFA |
| CAV2 | CCNK | CD93 | CDKN2AIP | P298- |
| CAVIN1 | CCNL1 | CD99 | CDKN2C | TCP10L;CFA |
| CAVIN2 | CCNY | CD99L2 | CDS1 | P298- |
| CAVIN3 | CCNYL1 | CDA | CDS2 | TCP10L;CFA |
| CBFB | CCS | CDC16 | CDSN | P298- |
| CBFB | CCT2 | CDC23 | CDV3 | TCP10L;CFA |
| CBL | CCT3 | CDC34 | CEACAM5 | P298;CFAP29 |
| CBLC | CCT4 | CDC37 | CEACAM7 | 8;CFAP298;C |
| CBR1 | CCT5 | CDC37L1 | CEBPZ | FAP298;CFA |
| CBR3 | CCT6A | CDC40 | CELA3B | P298 |
| CBR4 | CCT7 | CDC42 | CELF1 | CFAP57 |

CFAP69
CFD
CFH
CFHR1
CFHR2
CFHR4
CFHR5
CFI
CFL1
CFL2
CFLAR
CFP
CGGBP1
CGNL1
CHAD
CHAMP1
CHCHD10
CHCHD2
CHCHD3
CHCHD4
CHCHD5
CHCHD6
CHD1L
CHD3
CHD4
CHD8;CHD7;
CHD8;CHD8;
CHD7
CHERP
CHGB
CHID1
CHKB
CHL1
CHMP1A
CHMP1B
CHMP2A
CHMP2B
CHMP3
CHMP4A
CHMP4B
CHMP6
CHN1
CHORDC1
CHP1
CHRAC1
CHRD
CHRDL1
CHST13
CHST14
CHST7
CHTOP
CHUK
CIAO1
CIAO2A
CIAO2B
CIAO3
CIAPIN1
CIC
CILP
CILP2
CIMAP2
CIRBP
CISD1
CISD2
CIZ1
CKAP4
CKAP5
CKB
CKM
CKMT1A
CKMT2
CLASP1
CLASP2
CLC
CLCA2
CLCA4
CLCC1
CLCN7
CLCNKB
CLDN1
CLDN5
CLDND1
CLEC11A
CLEC14A
CLEC16A
CLEC2B
CLEC3B
CLIC1
CLIC2
CLIC3
CLIC4
CLIC5
CLIC6
CLINT1
CLIP1
CLIP1
CLIP2
CLMN
CLN3
CLN5
CLN6;CLN6;
CLN6;CLN6;;
CLN6;CLN6;
CLN6;CLN6
CLNS1A
CLP1
CLPB
CLPP
CLPTM1
CLPX
CLSTN1
CLTA
CLTB
CLTC
CLU
CLUAP1
CLUH
CLYBL
CMA1
CMAS
CMBL
CMC1
CMC2
CMC4
CMIP
CMPK1
CMPK2
CMSS1
CMTM3
CMTM4
CMTR1
CNBP
CNDP1
CNDP2
CNEP1R1
CNFN
CNIH4
CNKSR1
CNN1
CNN1
CNN2
CNN3
CNN3
CNNM2
CNNM3
CNOT1
CNOT10
CNOT11
CNOT2
CNOT3
CNOT4
CNOT6L
CNOT7
CNOT9
CNP
CNPY2
CNPY3
CNPY4
CNRIP1
CNTFR
CNTN1
CNTNAP1
CNTNAP3B;C
NTNAP3B;CN
TNAP3B;CNT
NAP3B;CNTN
AP3C;CNTNA
P3B;CNTNAP
3C;CNTNAP3
;CNTNAP3;C
NTNAP3;CNT
NAP3B;CNTN
AP3;CNTNAP
3B;CNTNAP3
CNTNAP4;C
NTNAP4;;CN
TNAP4;CNTN
AP4;CNTNAP
4
CNTNAP5
COA3
COA5
COA6
COA7
COASY
COBL
COBLL1
COCH
COG2
COG3
COG4
COG5
COG6
COG7
COG8
COIL
COL11A1
COL12A1
COL14A1
COL15A1
COL16A1
COL17A1
COL18A1
COL1A1
COL1A2
COL21A1
COL26A1
COL28A1
COL2A1
COL3A1
COL4A1
COL4A2
COL4A5
COL4A6
COL5A1
COL5A2
COL5A3
COL6A1
COL6A2
COL6A3
COL6A3
COL6A6
COL7A1
COL8A1
COL8A2

COLEC12
COLGALT1
COMMD1
COMMD10
COMMD2
COMMD3
COMMD3-BMI1
COMMD4
COMMD5
COMMD6
COMMD8
COMMD9
COMP
COMT
COMTD1
COPA
COPB1
COPB2
COPE
COPG1
COPG2
COPS2
COPS3
COPS4
COPS5
COPS6
COPS7A
COPS7B
COPS8
COPZ1
COPZ2
COQ10B
COQ3
COQ5
COQ6
COQ7
COQ8A
COQ9
CORO1A
CORO1B
CORO1C
CORO2A
CORO2B
COTL1
COX17
COX20
COX4I1
COX5A
COX5B
COX6A1
COX6B1
COX6C
COX7A1
COX7A2
COX7C
CP
CPA3
CPA4
CPA6
CPB2
CPD
CPE
CPEB2;CPEB2;CPEB2;CPEB2;CPEB2;CPEB3
CPM
CPN1
CPN2
CPNE1
CPNE2
CPNE3
CPOX
CPPED1
CPQ
CPSF1
CPSF2
CPSF3
CPSF4
CPSF6
CPSF7
CPT1A
CPT2
CPVL
CPXM2
CPZ
CR1
CRABP1
CRABP2
CRADD
CRAT
CRBN
CRCT1
CREB1
CREG1
CRELD1
CRELD2
CRIP1
CRIP2
CRISP3
CRISPLD2
CRK
CRKL
CRLF1
CRMP1
CRNKL1
CRNN
CROCC
CROT
CRPPA
CRTAC1
CRTAP
CRTC1
CRTC3
CRY2
CRYAB
CRYBG1
CRYBG2
CRYL1
CRYM
CRYZ
CRYZL1
CS
CS
CSAD
CSDE1
CSE1L
CSK
CSNK1A1
CSNK1D
CSNK1E
CSNK1G2
CSNK2A1
CSNK2A2
CSNK2B
CSPG4
CSRP1
CSRP2
CST3
CST6
CSTA
CSTB
CSTF1
CSTF2
CSTF3
CTBP1
CTBP2
CTBS
CTCF
CTDNEP1
CTDP1
CTDSP1
CTHRC1
CTIF
CTNNA1
CTNNAL1
CTNNB1
CTNNBIP1
CTNNBL1
CTNND1
CTNND2
CTPS1
CTPS2
CTR9
CTRB2;CTRB1;CTRB1;CTRB2
CTSA
CTSB
CTSC
CTSD
CTSF
CTSG
CTSH
CTSK
CTSL
CTSS
CTSV
CTSZ
CTTN
CTU2
CUL1
CUL2
CUL3
CUL4A
CUL4B
CUL5
CUL7
CUTA
CUTC
CWC15
CWC22
CWF19L1
CWH43
CXADR
CXCL12
CXCL13
CXCL14
CXCL17
CXXC1
CYB5A
CYB5B
CYB5R1
CYB5R2
CYB5R3
CYBA
CYBB
CYBC1
CYBRD1
CYC1
CYCS
CYFIP1
CYFIP2
CYGB
CYLD
CYP1B1
CYP20A1
CYP27A1
CYP2S1
CYP2U1
CYP4F11
CYP4F12
CYP4F22

CYP4X1
CYP51A1
CYP7B1
CYRIA
CYRIB
CYSRT1
CYSTM1
CYTH1
CZIB
D2HGDH
DAAM1
DAAM2
DAB2
DAB2IP
DACT3
DAD1
DAG1
DAGLB
DAP
DAP3
DAPK2
DAPK3
DAPL1
DARS1
DARS2
DAXX
DAZAP1
DBH
DBI
DBN1
DBNL
DBNL
DBR1
DBT
DCAF1
DCAF11
DCAF13
DCAF16
DCAF5
DCAF7
DCAF8
DCAKD
DCK
DCLK1
DCLK2
DCN
DCP1A
DCPS
DCTD
DCTN1
DCTN2
DCTN3
DCTN4
DCTN5
DCTN6
DCTPP1
DCUN1D1
DCUN1D4
DCUN1D5
DCXR
DDA1
DDAH1
DDAH2
DDB1
DDB2
DDHD2
DDI2
DDOST
DDR1
DDR2
DDRGK1
DDT
DDX1
DDX17
DDX18
DDX19A
DDX21
DDX23
DDX24
DDX27
DDX39A
DDX39B
DDX39B
DDX3X
DDX3Y
DDX41
DDX42
DDX46
DDX47
DDX49
DDX5
DDX50
DDX52
DDX54
DDX55
DDX56
DDX6
DDX60
DECR1
DECR2
DEF6
DEFA1;DEFA
3
DEFA5
DEFB1
DEGS1
DEK
DENND10
DENND1A
DENND2B
DENND2D
DENND3
DENND4C
DENND5A
DENR
DEPTOR
DERA
DERL1
DERL2
DERPC
DES
DFFA
DFFB
DGAT1
DGAT2
DGKA
DGKB
DGKG
DGKQ
DGLUCY
DGLUCY
DHCR24
DHCR7
DHODH
DHRS1
DHRS11
DHRS2
DHRS3
DHRS4
DHRS7
DHRS7B
DHTKD1
DHX15
DHX16
DHX29
DHX30
DHX32
DHX36
DHX37
DHX38
DHX40
DHX57
DHX58
DHX8
DHX9
DIABLO
DIAPH1
DIAPH2
DICER1
DIDO1
DIMT1
DIP2A
DIP2B
DIP2C
DIPK2A
DIS3
DIS3L
DIS3L2
DKC1
DKK2
DKK3
DLAT
DLD
DLG1
DLG2
DLG3
DLGAP2
DLGAP4
DLST
DMAC1
DMAC2L
DMD
DMKN
DMKN
DMPK
DMTN
DMXL1
DNAAF10
DNAAF5
DNAJA1
DNAJA2
DNAJA3
DNAJA4
DNAJB1
DNAJB11
DNAJB12
DNAJB14
DNAJB2
DNAJB4
DNAJB5
DNAJB6
DNAJC1
DNAJC10
DNAJC11
DNAJC13
DNAJC16
DNAJC19
DNAJC2
DNAJC21
DNAJC25-
GNG10
DNAJC3
DNAJC5
DNAJC7
DNAJC8
DNAJC9
DNAL1
DNASE1L1
DNASE1L3
DNASE2
DNM1
DNM1L
DNM2
DNM3
DNMBP

DNPEP
DNPEP
DNPH1
DNTTIP1
DOCK1
DOCK11
DOCK2
DOCK4
DOCK5
DOCK6
DOCK7
DOCK8
DOCK9
DOHH
DOP1A
DOP1B
DPAGT1
DPCD
DPF2
DPH1
DPH2
DPH5
DPH6
DPM1
DPM3
DPP3
DPP7
DPP8
DPP9
DPT
DPY19L1
DPY30
DPYD
DPYSL2
DPYSL3
DR1
DRAP1
DRG1
DRG2
DSC1
DSC2
DSC2
DSC3
DSG1
DSG2
DSG3
DSN1
DSP
DST
DST
DSTN
DTD1
DTNA
DTNA
DTNBP1
DTX2
DTX3
DTX3L
DTYMK
DUOX1
DUOXA1
DUS4L-
BCAP29;DUS
4L-
BCAP29;BCA
P29
DUSP12
DUSP14
DUSP22
DUSP23
DUSP3
DUT
DVL1
DVL2
DYNC1H1
DYNC1I2
DYNC1I2
DYNC1LI1
DYNC1LI2
DYNC2H1
DYNLL1
DYNLL2
DYNLRB1
DYNLT1
DYNLT3
DYRK1A
DYSF
E2F4
E2F7
EARS2
EBF1
EBI3
EBNA1BP2
EBP
ECD
ECE1
ECH1
ECHDC1
ECHDC2
ECHDC3
ECHS1
ECI1
ECI2
ECM1
ECM2
ECPAS
EDC3
EDC4
EDEM3
EDF1
EDIL3
EEA1
EED
EEF1A1
EEF1A2
EEF1B2
EEF1D
EEF1E1
EEF1G
EEF2
EEF2K
EEFSEC
EEPD1
EFCAB14
EFEMP1
EFEMP2
EFHD1
EFHD2
EFL1
EFNB1
EFNB2
EFR3A
EFTUD2
EGFL7
EGFR
EHBP1
EHBP1L1
EHD1
EHD2
EHD3
EHD4
EHHADH
EHMT2
EI24
EIF1
EIF1AX
EIF1AY
EIF2A
EIF2AK2
EIF2AK4
EIF2B1
EIF2B2
EIF2B3
EIF2B4
EIF2B5
EIF2D
EIF2S1
EIF2S2
EIF2S3
EIF3A
EIF3B
EIF3C
EIF3D
EIF3E
EIF3F
EIF3G
EIF3H
EIF3I
EIF3I
EIF3J
EIF3K
EIF3L
EIF3M
EIF4A1
EIF4A2
EIF4A3
EIF4B
EIF4E
EIF4E2
EIF4E3
EIF4ENIF1
EIF4G1
EIF4G2
EIF4G3
EIF4H
EIF5
EIF5A
EIF5B
EIF6
EIPR1
ELAC2
ELANE
ELAVL1
ELF1
ELF3
ELMO1
ELMO2
ELMO3
ELMOD2
ELN
ELN
ELOB
ELOC
ELOVL1
ELOVL7
ELP1
ELP2
ELP3
ELP3
ELP4
ELP5
EMC1
EMC10
EMC2
EMC3
EMC4
EMC6
EMC7
EMC8
EMD
EMG1
EMID1
EMILIN1
EMILIN2
EMILIN3

EML1
EML2
EML2
EML3
EML4
EMP3
ENAH
ENDOD1
ENDOG
ENDOU
ENG
ENGASE
ENO1
ENO2
ENOPH1
ENOSF1
ENPP1
ENPP2
ENPP4
ENPP6
ENSA
ENTPD1
ENTPD2
ENTPD3
ENTPD5
ENTR1
ENY2
EOGT
EPB41
EPB41L1
EPB41L2
EPB41L3
EPB41L3
EPB41L4A
EPB42
EPDR1
EPG5
EPHA1
EPHA2
EPHA3
EPHA4
EPHB2
EPHB3
EPHB4
EPHX1
EPHX2
EPHX3
EPM2A
EPM2AIP1
EPN1
EPN2
EPN3
EPPK1
EPRS1
EPS15
EPS15L1
EPS8
EPS8L1
EPS8L2
EPX
ERAP1
ERAP2
ERBB2
ERBIN
ERC1
ERCC1
ERCC2
ERCC3
ERG
ERG28
ERGIC1
ERH
ERI1
ERI3
ERLEC1
ERLIN1
ERLIN2
ERMP1
ERO1A
ERP29
ERP44
ESAM
ESD
ESPN
ESR1
ESRP1
ESRP2
ESRRA
ESYT1
ESYT2
ETF1
ETFA
ETFB
ETFDH
ETFRF1
ETHE1
ETV6
EVA1C
EVL
EVPL
EWSR1
EXD2
EXOC1
EXOC2
EXOC3
EXOC4
EXOC5
EXOC6
EXOC6B
EXOC7
EXOC8
EXOG
EXOSC10
EXOSC2
EXOSC3
EXOSC5
EXOSC6
EXOSC7
EXOSC8
EXOSC9
EXT1
EXT2
EXTL2
EZR
F10
F11
F11R
F12
F12
F13A1
F13B
F2
F3
F7
F8A1
F9
FAAH
FABP3
FABP4
FABP5
FABP5
FADD
FADS2
FADS6
FAF1
FAF2
FAH
FAHD1
FAHD2A
FAIM
FAM114A1
FAM114A2
FAM120A
FAM120A
FAM120B
FAM120C
FAM135A
FAM162A
FAM168A
FAM177A1
FAM180A
FAM180B
FAM210B
FAM234A
FAM241A
FAM25A;FAM25C;FAM25G
FAM3A
FAM3B
FAM3C
FAM3D
FAM50A
FAM83A
FAM83B
FAM83C
FAM83G
FAM83H
FAM8A1
FAM91A1
FAM98A
FAM98B
FAM98C
FAP
FAR1
FARP1
FARP2
FARS2
FARSA
FARSB
FAS
FASN
FAT2
FAU
FBH1
FBL
FBLIM1
FBLN1
FBLN1
FBLN2
FBLN5
FBLN7
FBN1
FBN2
FBP1
FBP2
FBXL18
FBXL20
FBXL22
FBXL8
FBXO17
FBXO2
FBXO21
FBXO22
FBXO27
FBXO3
FBXO30
FBXO4
FBXO44
FBXO45
FBXO6
FBXO7
FBXO9
FBXW2
FCER1G
FCGBP

FCGR2B;FCGR2C
FCGRT
FCHO2
FCSK
FDFT1
FDPS
FDX1
FDX2
FDXR
FECH
FEM1B
FEN1
FER
FERMT1
FERMT2
FERMT3
FERRY3
FETUB
FGA
FGB
FGD4
FGD5
FGF10
FGF2
FGF22
FGF7
FGFBP1
FGFR1
FGFR2
FGFR3
FGG
FGGY
FGL2
FH
FHIP1A
FHIP2A
FHIP2B
FHL1
FHL2
FHL3
FHL5
FIBIN
FIBP
FIG4
FILIP1
FILIP1L
FIS1
FITM2
FKBP10
FKBP11
FKBP15
FKBP1A
FKBP2
FKBP3
FKBP4
FKBP5
FKBP7
FKBP8
FKBP9
FKRP
FLAD1
FLG
FLG2
FLI1
FLII
FLNA
FLNB
FLNC
FLOT1
FLOT2
FLT1
FLT3LG
FLYWCH2
FMNL1
FMNL2
FMO1
FMO3
FMOD
FMR1
FN1
FN3K
FN3KRP
FNBP1
FNBP1L
FNBP4
FNDC1
FNDC3A
FNDC3B
FNTA
FNTB
FOCAD
FOLR2
FOSL2
FOXK1
FOXP1
FPGT
FRAS1
FRG1
FRMD6
FRMD8
FRMPD2;FRMPD2B
FRRS1
FRS2
FRY
FRYL
FRZB
FSCN1
FST
FSTL1
FSTL3
FTH1
FTL
FTO
FTSJ3
FUBP1
FUBP3
FUCA1
FUCA2
FUNDC2
FUS
FUT2
FUT3
FXN
FXR1
FXR2
FXYD1
FXYD3
FYCO1
FYN
FZD7
G3BP1
G3BP2
G6PC3
G6PD
GAA
GAB1
GABARAP
GABARAPL2
GABBR2
GABPA
GAK
GAL
GAL3ST4
GALC
GALE
GALK1
GALK2
GALM
GALNT1
GALNT10
GALNT16
GALNT2
GALNT3
GALNT5
GALNT7
GALT
GAMT
GAN
GANAB
GANAB
GAP43
GAPDH
GAPVD1
GAR1
GARS1
GART
GAS2
GAS2L1
GAS6
GAS7
GATAD2A
GATAD2B
GATD1
GATD3
GATM
GBA1
GBE1
GBF1
GBP1
GBP2
GBP3
GBP4
GBP6
GC
GCA
GCC2
GCDH
GCHFR
GCLC
GCLM
GCN1
GCSH
GDAP1
GDAP2
GDE1
GDF11
GDI1
GDI2
GDI2
GDPD3
GEM
GEMIN2
GEMIN4
GEMIN5
GET1
GET3
GET4
GFAP
GFER
GFM1
GFM2
GFOD1
GFPT1
GFUS
GGA1
GGACT
GGCT
GGCX
GGH
GGPS1
GGT5
GGT6
GGT7

GHDC
GID8
GIGYF2
GIMAP1
GIMAP1-
GIMAP5
GIMAP4
GIMAP7
GIMAP8
GIPC1
GIPC2
GIPC3
GIT1
GIT2
GJA1
GK
GLA
GLB1
GLE1
GLG1
GLIPR2
GLMN
GLO1
GLOD4
GLOD4
GLRX
GLRX3
GLRX5
GLS
GLS
GLT8D1
GLT8D2
GLTP
GLUD1
GLUL
GLYR1
GM2A
GMDS
GMFB
GMFG
GMPPA
GMPPB
GMPR
GMPR2
GMPS
GNA11
GNA12
GNA13
GNA15
GNAI1
GNAI2
GNAI3
GNAO1
GNAQ
GNAS
GNAS
GNAZ
GNB1
GNB1L
GNB2
GNB3
GNB4
GNE
GNG12
GNG2
GNG4
GNG5
GNG7
GNL1
GNL3
GNPAT
GNPDA1
GNPDA2
GNPNAT1
GNS
GOLGA2
GOLGA3
GOLGA4
GOLGA5
GOLGA7
GOLGB1
GOLIM4
GOLM1
GOLPH3
GOLPH3L
GOLT1B
GON7
GOPC
GORASP1
GORASP2
GOSR1
GOSR2;GOS
R2;GOSR2;G
OSR2;GOSR2
;GOSR2;GOS
R2;GOSR2;G
OSR2;;GOSR
2;GOSR2
GOT1
GOT2
GP6
GPAA1
GPALPP1
GPC1
GPC4
GPC6
GPD1
GPD1L
GPD2
GPHN
GPI
GPKOW
GPLD1
GPM6A
GPM6B
GPN1
GPNMB
GPR107
GPR108
GPR155
GPR15LG
GPR89A;GPR
89B
GPS1
GPS1
GPX1
GPX2
GPX3
GPX4
GPX7
GPX8
GRAMD4
GRAP2
GRB10
GRB2
GRB7
GREM1
GREM2
GRHL1
GRHL2
GRHPR
GRIPAP1
GRK2
GRK5
GRM2
GRN
GRPEL1
GRSF1
GRTP1
GRWD1
GSDMA
GSDMC
GSDMD
GSDME
GSK3A
GSK3B
GSKIP
GSN
GSN
GSPT1
GSPT2
GSR
GSS
GSTA1
GSTA3
GSTA4
GSTK1
GSTM1
GSTM1
GSTM2
GSTM3
GSTM4
GSTM5
GSTO1
GSTP1
GSTP1
GSTT1
GSTT2;GSTT2
;GSTT2B;GST
T2B
GSTZ1
GSTZ1
GTF2A1
GTF2A2
GTF2B
GTF2F2
GTF2H1
GTF2H2;GTF
2H2C
GTF2H3
GTF2H4
GTF2I
GTF3C1
GTF3C2
GTF3C3
GTF3C4
GTF3C5
GTF3C6
GTPBP1
GTPBP10
GTPBP4
GUCY1A1
GUCY1A2
GUCY1B1
GUF1
GUK1
GUSB
GXYLT2
GYG1
GYPC
GYS1
GZMB
GZMK
H1-0
H1-1
H1-10
H1-2
H1-3
H1-4
H1-5
H2AC11;H2A
C12;H2AC14
;H2AJ
H2AC20;H2A
C18

H2AC4;H2AC
25
H2AX
H2AZ1;H2AZ
2
H2BC12;H2B
C4
H2BC20P;H2
BC19P
H2BK1
H3-3B;H3-3A
H3-7
H4C1
H6PD
HAAO
HABP2
HACD2
HACD3
HACE1
HADH
HADHA
HADHB
HAGH
HAL
HAPLN1
HAPLN3
HARS1
HARS2
HAT1
HAX1
HBA1
HBA1
HBB
HBD
HBE1
HBG2
HBQ1
HBS1L
HCAR3;HCA
R2
HCCS
HCFC1
hCG_204342
6
HCLS1

HDAC1
HDAC2
HDAC4
HDAC6
HDAC7
HDAC8;HDA
C8;;HDAC8;
HDAC8;HDA
C8;HDAC8;H
DAC8;HDAC
8;HDAC8;HD
AC8;HDAC8;
HDAC8
HDDC2
HDDC3
HDGF
HDGFL2
HDGFL3
HDHD2
HDHD3
HDHD5
HDLBP
HEATR1
HEATR3
HEATR5A
HEATR5B
HEATR6
HEBP1
HEBP2
HECTD1
HECTD3
HECTD4
HELZ2
HEPH
HERC4
HERC6
HEXA
HEXB
HEXIM1
HFE
HGFAC
HGH1
HGS
HGSNAT
HHAT

HHIPL2
HIBADH
HIBCH
HIBCH
HID1
HIKESHI
HINT1
HINT2
HINT3
HIP1
HIP1R
HIRA
HK1
HK2
HLA-A
HLA-A
HLA-B
HLA-B
HLA-B
HLA-B
HLA-B
HLA-C
HLA-C
HLA-C
HLA-C
HLA-C
HLA-DMA
HLA-DMB
HLA-DPA1
HLA-DPB1
HLA-DPB1
HLA-
DQA1;HLA-
DQA2
HLA-DQB1
HLA-DQB1
HLA-DRA
HLA-DRB1
HLA-DRB1
HLA-DRB1
HLA-DRB3
HLA-E
HLA-F
HM13
HMBS

HMCES
HMCN1
HMCN2
HMG20A
HMGA1
HMGB1
HMGB2
HMGCL
HMGCS1
HMGCS2
HMGN1
HMGN2
HMGN3
HMGN4
HMGN5
HMOX1
HMOX2
HNMT
HNRNPA0
HNRNPA1
HNRNPA2B1
HNRNPA3
HNRNPAB
HNRNPC
HNRNPD
HNRNPD
HNRNPDL
HNRNPF
HNRNPH1
HNRNPH2
HNRNPH3
HNRNPK
HNRNPK
HNRNPL
HNRNPLL
HNRNPM
HNRNPR
HNRNPU
HNRNPUL1
HNRNPUL2
HOMER2
HOOK1
HOOK2
HOOK3
HOPX

HP
HP1BP3
HPCAL1
HPF1
HPGD
HPGDS
HPR
HPRT1
HPSE
HPSE2
HPX
HRAS
HRC
HRG
HRNR
HS1BP3
HSBP1
HSCB
HSD11B1
HSD17B10
HSD17B11
HSD17B12
HSD17B13
HSD17B14
HSD17B4
HSD17B6
HSD17B8
HSD3B7
HSDL1
HSDL2
HSP90AA1
HSP90AA4P
HSP90AB1
HSP90B1
HSPA12A
HSPA12B
HSPA13
HSPA14
HSPA1B;HSP
A1A;HSPA1B
HSPA1L
HSPA2
HSPA4
HSPA4L
HSPA5

HSPA6
HSPA8
HSPA8
HSPA9
HSPB1
HSPB1
HSPB2-
C11orf52;HS
PB2
HSPB6
HSPB7
HSPB8
HSPBP1
HSPD1
HSPE1
HSPE1-
MOB4
HSPG2
HSPH1
HTATIP2
HTATSF1
HTRA1
HTRA2
HTRA3
HTT
HUWE1
HVCN1
HYCC1
HYDIN
HYI
HYOU1
HYPK
IAH1
IARS1
IARS1
IARS2
IBA57
ICAM1
ICAM2
ICAM3
ICMT
ICOSLG
IDE
IDH1
IDH2
IDH3A
IDH3B
IDH3G
IDI1
IDUA
IER3IP1
IFFO1;IFFO1;
IFFO1;IFFO2
IFI16
IFI30
IFI35
IFI44
IFIH1
IFIT1
IFIT2
IFIT3
IFIT5
IFITM1
IFITM3
IFT122
IFT20
IFT25
IFT27
IFT74
IFT81
IFT88
IGBP1
IGF1
IGF2
IGF2BP2
IGF2R
IGFALS
IGFBP2
IGFBP3
IGFBP4
IGFBP5
IGFBP6
IGFBP7
IGHA1
IGHA2
IGHA2
IGHD
IGHG1
IGHG2
IGHG3
IGHG4
IGHM
IGHV1-18
IGHV1-2
IGHV1-24
IGHV1-3
IGHV1-45
IGHV1OR15-
1;
IGHV2-26
IGHV2-70
IGHV2-70D
IGHV3-13
IGHV3-35
IGHV3-38
IGHV3-38-3
IGHV3-43
IGHV3-49
IGHV3-64
IGHV3-64D
IGHV3-7
IGHV3-72
IGHV3-73
IGHV3OR16-
12
IGHV4-28
IGHV4-34
IGHV4-4
IGHV4OR15-
8
IGHV5-51
IGHV6-1
IGKC
IGKJ1
IGKJ4
IGKV1-16
IGKV1-17
IGKV1-27
IGKV1-33
IGKV1-
37;IGKV1D-
37
IGKV1-5
IGKV1-6
IGKV1D-
13;IGKV1-13
IGKV1D-16
IGKV2-
28;IGKV2D-
28
IGKV2-29
IGKV2-
40;IGKV2-
40;IGKV2-
40;IGKV2D-
40
IGKV2D-24
IGKV3-20
IGKV3-
7;IGKV3D-
7;IGKV3OR2-
268
IGKV3D-
11;IGKV3-11
IGKV3D-15
IGKV3D-20
IGKV4-1
IGLC2;IGLC3
IGLC7
IGLL1
IGLL5
IGLV1-40
IGLV1-47
IGLV1-51
IGLV2-14
IGLV2-18
IGLV3-19
IGLV3-21
IGLV3-25
IGLV3-27
IGLV3-9
IGLV4-60
IGLV4-69
IGLV5-45
IGLV6-57
IGLV7-
46;IGLV7-43
IGLV8-61
IGLV9-49
IGSF1
IGSF3
IGSF8
IK
IKBIP
IKBKB
IKBKG
IL16
IL17D
IL18
IL1RN
IL33
IL36RN
IL4I1
IL6ST
ILF2
ILF3
ILK
ILKAP
ILVBL
IMMT
IMMT
IMP3
IMP4
IMPA1
IMPA2
IMPACT
IMPDH
IMPDH1
IMUP
INF2
ING5;ING5;I
NG5;ING4
INHBE
INMT
INPP1
INPP4A
INPP4B
INPP5A
INPP5D
INPP5F
INPP5K
INPPL1
INSR
INTS1

INTS10
INTS11
INTS13
INTS2
INTS3
INTS4
INTS5
INTS8
INTS9
IPO11
IPO13
IPO4
IPO5
IPO7
IPO8
IPO9
IQCA1
IQGAP1
IQGAP2
IQSEC1
IQSEC2
IRAG1
IRAG2
IRAK1
IRAK3
IRAK4
IREB2
IRF2
IRF2BP1
IRF2BP2
IRF2BPL
IRF3
IRF9
IRGQ
ISCA1
ISCA2
ISCU
ISG15
ISLR
ISOC1
ISOC2
IST1
ISYNA1
ITCH
ITFG1
ITGA1
ITGA2
ITGA2B
ITGA3
ITGA4
ITGA5
ITGA6
ITGA7
ITGA7
ITGA8
ITGA9
ITGAM
ITGAV
ITGB1
ITGB2
ITGB3
ITGB4
ITGB4
ITGB5
ITGBL1
ITIH1
ITIH2
ITIH3
ITIH4
ITIH5
ITLN1
ITM2B
ITM2C
ITPA
ITPK1
ITPKB
ITPKC
ITPR1
ITPR2
ITPR3
ITPRID2
ITPRIP
ITSN1
ITSN2
IVD
IVL
IVNS1ABP
IWS1
JAGN1
JAK1
JAM2
JAM3
JCAD
JCHAIN
JMJD7
JOSD2
JPH2
JPT1
JPT2
JUN
JUP
KALRN
KANK1
KANK2
KANK3
KARS1
KAT6B
KAT7
KATNA1
KATNAL2
KBTBD11
KCMF1
KCNAB1
KCNAB2
KCNJ10
KCNMA1
KCNU1
KCTD10
KCTD12
KCTD15
KCTD21
KDELR1
KDM1A
KDM2A
KDM3B
KDM4B
KDM5A
KDM5B
KDSR
KEAP1
KERA
KGD4
KHDRBS1
KHDRBS3
KHNYN
KHSRP
KIAA0319L
KIAA0408
KIAA1217
KIAA1671
KIAA2013
KIDINS220
KIF13A
KIF13B
KIF1B
KIF1C
KIF21A
KIF2A
KIF3A
KIF5A
KIF5B
KIF5C
KIFAP3
KIFBP
KIFC3
KIN
KIRREL1
KIT
KLC3
KLC4
KLF4
KLF5
KLHL15
KLHL22
KLHL28
KLHL42
KLK10
KLK11
KLK12
KLK13
KLK14
KLK5
KLK6
KLK7
KLK8
KMT2A
KMT2C
KMT2D
KNG1
KNTC1
KPLCE
KPNA1
KPNA2
KPNA3
KPNA4
KPNA6
KPNB1
KRAS
KRI1
KRR1
KRT1
KRT10
KRT13
KRT13
KRT14
KRT15
KRT16
KRT17
KRT18
KRT19
KRT2
KRT23
KRT24
KRT3
KRT32
KRT4
KRT5
KRT6A
KRT6B
KRT6C
KRT7
KRT71
KRT72
KRT73
KRT75
KRT76
KRT77
KRT78
KRT8
KRT80
KRT9
KRTCAP2
KRTDAP
KSR1
KTN1

KXD1
KYAT3
L1CAM
L2HGDH
L3HYPDH
LACC1
LACTB
LACTB2
LAD1
LAGE3
LAMA2
LAMA3
LAMA4
LAMA5
LAMB1
LAMB2
LAMB3
LAMC1
LAMC2
LAMC3
LAMP1
LAMP2
LAMTOR1
LAMTOR2
LAMTOR3
LAMTOR4
LAMTOR5
LANCL1
LANCL2
LAP3
LARP1
LARP4
LARP4B
LARP7
LARS1
LARS2
LAS1L
LASP1
LBH
LBP
LBR
LCK
LCLAT1
LCMT1
LCN2
LCP1
LDAH
LDB1
LDB3
LDB3
LDHA
LDHB
LECT2
LEFTY2;LEFTY1
LEMD2
LEMD3
LEO1
LETM1
LETMD1
LFNG
LGALS1
LGALS3
LGALS3BP
LGALS7
LGALS8
LGALS9C;LGALS9B;LGALS9C
LGALSL
LGI3
LGI4
LGMN
LHFPL2
LHPP
LIG1
LIG3
LIG4
LILRB4
LILRB5
LIMA1
LIMCH1
LIMD1
LIMK1
LIMK2
LIMS1
LIMS1
LIMS1
LIMS2
LIMS3
LIN7C
LIN9
LIPA
LLGL1
LLGL2
LMAN1
LMAN2
LMAN2L
LMBRD1
LMCD1
LMF2
LMNA
LMNA
LMNB1
LMNB2
LMO7
LMOD1
LNPEP
LNPK
LOC100653049
LOC122394732
LONP1
LOX
LOXL1
LOXL3
LOXL4
LPA
LPAR1
LPCAT1
LPCAT2
LPCAT3
LPCAT4
LPGAT1
LPP
LPXN
LRBA
LRCH1
LRCH3
LRG1
LRP1
LRPAP1
LRPPRC
LRRC1
LRRC14
LRRC15
LRRC17
LRRC20
LRRC25
LRRC32
LRRC40
LRRC41
LRRC47
LRRC59
LRRC8A
LRRC8C
LRRFIP1
LRRFIP1
LRRFIP2
LRRK2
LRRN4CL
LRSAM1
LRWD1
LSAMP
LSG1
LSM1
LSM12
LSM14A
LSM14B
LSM2
LSM3
LSM4
LSM5
LSM6
LSM7
LSM8
LSP1
LSR
LSS
LTA4H
LTBP1
LTBP2
LTBP3
LTBP4
LTC4S
LTF
LTN1
LUC7L2
LUC7L3
LUM
LUZP1
LXN
LY6D
LY6G6C
LY6G6C
LY75
LY96
LYAR
LYN
LYNX1;LYNX1;LYNX1-SLURP2;LYNX1
LYPD2
LYPD3
LYPD5
LYPLA1
LYPLA2
LYPLAL1
LYRM4
LYRM7
LYSMD4
LYST
LYVE1
LYZ
LZIC
LZTFL1
LZTR1
M6PR
MAB21L4
MAB21L4
MACF1
MACF1
MACROD1
MACROD2
MACROH2A1
MACROH2A1
MACROH2A1
MACROH2A2
MAD2L1
MAEA
MAGED2
MAGOH
MAGOHB

MAGT1
MAIP1
MAL2
MALSU1
MALT1
MAMDC2
MAN1A1
MAN1A2
MAN1B1
MAN2A1
MAN2A2
MAN2B1
MAN2B2
MAN2C1
MANBA
MANF
MAOA
MAOB
MAP1A
MAP1B
MAP1LC3A
MAP1LC3B2;
MAP1LC3B;M
AP1LC3B
MAP1S
MAP2
MAP2K1
MAP2K2
MAP2K3
MAP2K4
MAP2K6
MAP2K7
MAP3K20
MAP3K4
MAP3K5
MAP3K7
MAP4
MAP4
MAP4K4
MAP7
MAP7D1
MAPK1
MAPK13
MAPK14
MAPK1IP1L
MAPK3
MAPK9
MAPKAP1
MAPKAPK2
MAPKAPK3
MAPRE1
MAPRE2
MAPRE3
MAPT
MARCHF5
MARCKS
MARCKSL1
MARK2
MARK3
MASP1
MASP2
MAST2
MAST4
MAT2A
MAT2B
MATN2
MATN2
MATR3
MAU2
MAVS
MAX
MB
MB21D2
MBD1
MBD2
MBD3
MBLAC1
MBLAC2
MBNL1
MBOAT2
MBOAT7
MBP
MBP
MCAM
MCAT
MCCC1
MCCC2
MCEE
MCFD2
MCM2
MCM3
MCM4
MCM5
MCM6
MCM7
MCMBP
MCRIP1
MCTS1
MCU
MDH1
MDH2
MDK
MDN1
MDP1
ME1
ME2
ME3
MEAK7
MECP2
MECP2
MECR
MED12
MED14
MED15
MED16
MED18
MED20
MED23
MED24
MED27
MED30
MED4
MEF2A
MEF2D
MEGF6
MEIS1
MELTF
MEMO1
MEN1
MESD
MEST
METAP1
METAP2
METRN
METRNL
METTL1
METTL13
METTL16
METTL26
METTL3
MFAP2
MFAP4
MFAP5
MFAP5
MFF
MFGE8
MFN1
MFN2
MFNG
MFSD1
MFSD10
MFSD5
MGAT1
MGLL
MGMT
MGP
MGST1
MGST2
MGST3
MIA;MIA-
RAB4B
MIA2
MIA3
MICAL1
MICAL2
MICAL3
MICALL1
MICOS10;MI
COS10;MICO
S10-
NBL1;MICOS
10-NBL1
MICOS13
MICU1
MICU2
MICU3
MID1
MIDEAS
MIEN1
MIF
MIF4GD
MINDY2
MINDY3
MINK1
MIOS
MIPEP
MITF
MIX23
MKKS
MKRN2
MLEC
MLF2
MLKL
MLST8
MLYCD
MMAA
MMAB
MME
MMGT1
MMP10
MMP11
MMP14
MMP19
MMP2
MMP23B
MMP28
MMRN1
MMRN2
MMS19
MMUT
MOB1B
MOB2
MOB3A
MOB3C
MOCS1
MOCS2
MOCS2
MOCS3
MOGS
MON1B
MON2
MORC3
MORF4L1
MOSPD2
MOV10

| | | | | |
|---|---|---|---|---|
| MOXD1 | MRPL27 | MRTFB | MTMR14 | MYH11 |
| MPC1 | MRPL28 | MRTO4 | MTMR2 | MYH14 |
| MPC2 | MRPL3 | MSH2 | MTMR6 | MYH7 |
| MPDU1 | MRPL34 | MSH3 | MTMR9 | MYH9 |
| MPDZ | MRPL37 | MSH6 | MTNAP1 | MYL1 |
| MPG | MRPL38 | MSI2 | MT-ND2 | MYL10 |
| MPHOSPH10 | MRPL39 | MSMO1 | MT-ND3 | MYL12A;MYL |
| MPHOSPH6 | MRPL4 | MSN | MT-ND4 | 12B;MYL12A |
| MPHOSPH8 | MRPL41 | MSR1 | MT-ND5 | MYL4 |
| MPI | MRPL43 | MSRA | MT-ND6 | MYL6 |
| MPO | MRPL44 | MSRB2 | MTOR | MYL6 |
| MPP1 | MRPL45 | MSRB3 | MTPAP | MYL6B |
| MPP7 | MRPL46 | MST1 | MTPN | MYL9 |
| MPPED2 | MRPL47 | MST1R | MTR | MYLK |
| MPRIP | MRPL48 | MSTO1 | MTRES1 | MYLK |
| MPST | MRPL49 | MT1E;MT1F; | MTREX | MYO18A |
| MPV17 | MRPL53 | MT1B;MT1L | MTSS2 | MYO19 |
| MPZ | MRPL55 | MT1X | MTURN | MYO1B |
| MPZL1 | MRPL55 | MTA1 | MTX1 | MYO1C |
| MPZL2 | MRPL58 | MTA2 | MTX2 | MYO1C |
| MRC1 | MRPL9 | MTA3 | MTX3 | MYO1D |
| MRC2 | MRPS11 | MTAP | MUC15 | MYO1E |
| MRE11 | MRPS12 | MTARC1 | MUC21 | MYO1F |
| MREG | MRPS14 | MTARC2 | MUC4 | MYO1G |
| MRGBP | MRPS17;hCG | MT-ATP6 | MUC5B | MYO5A |
| MRGPRF | _1984214;MR | MT-ATP8 | MUSTN1 | MYO5B |
| MRI1 | PS17 | MTCH1 | MVB12A | MYO6 |
| MRM2 | MRPS18A | MTCH2 | MVD | MYO9A |
| MRM3 | MRPS18B | MT-CO1 | MVK | MYO9B |
| MRNIP | MRPS2 | MT-CO2 | MVP | MYOC |
| MROH1 | MRPS21 | MT-CO3 | MX1 | MYOF |
| MRPL1 | MRPS22 | MT-CYB | MX2 | MYOZ1 |
| MRPL10 | MRPS23 | MTDH | MXRA5 | MYZAP |
| MRPL11 | MRPS24 | MTFMT | MXRA7 | MZB1 |
| MRPL13 | MRPS25 | MTFR1L | MXRA7 | MZT1 |
| MRPL14 | MRPS27 | MTG1 | MXRA8 | MZT2B;MZT2 |
| MRPL15 | MRPS28 | MTHFD1 | MYADM | B;MZT2A;MZT |
| MRPL16 | MRPS31 | MTHFD1L | MYBBP1A | 2B;MZT2A |
| MRPL17 | MRPS33 | MTHFD2 | MYCBP2 | N4BP1 |
| MRPL18 | MRPS34 | MTHFR | MYCT1 | N6AMT1 |
| MRPL19 | MRPS35 | MTIF2 | MYD88 | NAA10 |
| MRPL20 | MRPS5 | MTM1 | MYDGF | NAA15 |
| MRPL21 | MRPS6 | MTMR1 | MYEF2 | NAA15 |
| MRPL22 | MRPS7 | MTMR10 | MYG1 | NAA20 |
| MRPL24 | MRPS9 | MTMR12 | MYH10 | NAA25 |

NAA30
NAA35
NAA38
NAA40
NAA50
NAAA
NAALAD2
NACA
NADK2
NADSYN1
NAE1
NAGA
NAGK
NAGLU
NAIP
NAMPT
NANS
NAP1L1
NAP1L1
NAP1L4
NAPA
NAPG
NAPRT
NARS1
NARS2
NASP
NAT1
NAT10
NAV1
NAXD
NAXE
NBAS
NBEA
NBEAL1
NBEAL2
NBL1
NBN
NCALD
NCAM1
NCAM2
NCAPD3
NCBP1
NCBP2
NCCRP1
NCDN
NCEH1
NCF1B;NCF1
NCF2
NCF4
NCK1
NCK1
NCK2
NCKAP1
NCKAP1L
NCL
NCLN
NCOA5
NCOA7
NCOR1
NCOR2
NCSTN
NDE1
NDP
NDRG1
NDRG2
NDRG3
NDRG4
NDST1;NDST
2;
NDUFA1
NDUFA10
NDUFA11
NDUFA12
NDUFA13
NDUFA2
NDUFA3
NDUFA4
NDUFA4L2
NDUFA5
NDUFA6
NDUFA7
NDUFA8
NDUFA9
NDUFAB1
NDUFAF1
NDUFAF2
NDUFAF3
NDUFAF4
NDUFAF5
NDUFAF6
NDUFAF7
NDUFB1
NDUFB10
NDUFB11
NDUFB2
NDUFB3
NDUFB4
NDUFB5
NDUFB7
NDUFB8
NDUFB9
NDUFC2
NDUFS1
NDUFS2
NDUFS3
NDUFS4
NDUFS5
NDUFS6
NDUFS7
NDUFS8
NDUFV1
NDUFV2
NEBL
NECAP2
NECTIN1
NECTIN2
NECTIN4
NEDD1
NEDD4
NEDD4L
NEDD8
NEDD8-
MDP1
NEFH
NEFL
NEFM
NEGR1
NEK7
NEK9
NELFA
NELFB
NELFCD
NELFE
NEMF
NEMP1
NENF
NES
NEXN
NF1
NF2
NFATC2
NFATC4
NFIA
NFIB
NFIC
NFIX
NFKB1
NFKB2
NFKBIA
NFKBIB
NFS1
NFU1
NFYB
NFYC
NGF
NGFR
NGLY1
NHEJ1
NHERF1
NHERF2
NHLRC2
NHP2
NHSL3
NIBAN1
NIBAN2
NID1
NID2
NIF3L1
NIFK
NIP7
NIPBL
NIPSNAP1
NIPSNAP2
NIPSNAP2
NIPSNAP3A
NISCH
NIT1
NIT2
NKIRAS1
NKIRAS2
NLGN4Y;NLG
N4Y;NLGN4X
;NLGN4Y;NL
GN2
NLN
NLRP2
NLRX1
NMB
NMD3
NME1
NME1-NME2
NME3
NME4
NME7
NMES1
NMI
NMNAT1
NMNAT3
NMRAL1
NMRK1
NMT1
NMT2
NNMT
NNT
NNT
NOB1
NOC2L
NOC4L
NOG
NOL10
NOL11
NOL3
NOL6
NOL7
NOLC1
NOMO3;NO
MO3;NOMO3
;NOMO2
NONO
NOP10
NOP14
NOP2
NOP56
NOP58
NOP9

NOS1AP
NOS3
NOSIP
NOSTRIN
NOTCH2
NOTCH3
NOVA1
NOVA2
NPC1
NPC2
NPEPL1
NPEPPS
NPLOC4
NPLOC4
NPM1
NPM1
NPM2
NPM3
NPNT
NPNT
NPR2
NPTN
NPY
NQO1
NQO2
NR1H3;NR1
H3;NR1H3;N
R1H3;NR1H2
;NR1H2;NR1
H2;NR1H3
NR2C2AP
NR2F2
NR3C1
NRAS
NRBP1
NRBP2
NRDC
NRF1
NRM
NRP1
NRP2
NSDHL
NSF
NSFL1C
NSL1
NSMCE2
NSUN2
NSUN5
NT5C
NT5C2
NT5C3A
NT5C3A
NT5DC1
NT5DC2
NT5DC3
NT5E
NTAN1
NTHL1
NTM
NTMT1
NTN1
NTN4
NTPCR
NTRK2
NUB1
NUBP1
NUBP2
NUBPL
NUCB1
NUCB2
NUCKS1
NUDC
NUDCD1
NUDCD2
NUDCD3
NUDT1
NUDT16
NUDT16L1
NUDT19
NUDT2
NUDT21
NUDT3
NUDT4
NUDT5
NUDT9
NUMA1
NUMB
NUMBL
NUP107
NUP133
NUP153
NUP155
NUP160
NUP188
NUP205
NUP210
NUP214
NUP35
NUP37
NUP43
NUP50
NUP54
NUP58
NUP62
NUP85
NUP88
NUP93
NUP98
NUTF2
NXF1
NXN
NXT2;NXT1
NYNRIN
OAF
OARD1
OAS1
OAS2
OAS3
OAT
OBSL1
OCIAD1
OCIAD2
OCRL
ODF2
ODR4
OGA
OGDH
OGDH
OGFR
OGFRL1
OGN
OGT
OLA1
OLFM4
OLFML1
OLFML2B
OLFML3
OMD
OPA1
OPHN1
OPLAH
OPTN
OR52M1
OR5K1;OR5K
2
ORC2
ORC3
ORC4
ORM1
ORM2
ORMDL2
ORMDL3;OR
MDL3;ORMD
L1
OS9
OSBP
OSBPL10
OSBPL11
OSBPL1A
OSBPL2
OSBPL5
OSBPL6
OSBPL8
OSBPL9
OSGEP
OSGEPL1
OSTC
OSTF1
OSTM1
OTC
OTUB1
OTUD4
OTUD6B
OTUD7B
OVCA2
OXA1L
OXCT1
OXR1
OXSM
OXSR1
P2RX1
P2RX4
P2RX7
P3H1
P3H3
P3H4
P4HA1
P4HA2
P4HA3
P4HB
PA2G4
PAAF1
PABPC1
PABPC4
PABPN1
PACS1
PACS2
PACSIN2
PACSIN3
PAF1
PAFAH1B1
PAFAH1B2
PAFAH1B3
PAICS
PAIP1
PAIP2
PAK1
PAK1IP1
PAK2
PAK4
PALD1
PALLD
PALM
PALM2AKAP2
PALMD
PALS2
PAM
PAM16;COR
O7
PAMR1
PANK2
PANK4
PANX1
PAPLN
PAPOLA

PAPPA
PAPSS1
PAPSS2
PARD3
PARD3B
PARK7
PARN
PARP1
PARP10
PARP12
PARP14
PARP16
PARP4
PARP9
PARVA
PARVB
PARVG
PATJ
PAWR
PAXBP1
PAXX
PBDC1
PBLD
PBRM1
PBX1
PBXIP1
PC
PCBD1
PCBD2
PCBP1
PCBP2
PCCA
PCCB
PCDH1
PCDH7
PCF11
PCID2
PCIF1
PCK2
PCMT1
PCNA
PCNP
PCNT
PCOLCE
PCOLCE2
PCP4
PCP4L1
PCSK5
PCSK9
PCYOX1
PCYOX1L
PCYT1A
PCYT2
PDAP1
PDCD10
PDCD11
PDCD4
PDCD5
PDCD6
PDCD6IP
PDCL
PDCL3
PDE12
PDE1B
PDE2A
PDE3A
PDE4D
PDE5A
PDE6D
PDGFA
PDGFB
PDGFD
PDGFRA
PDGFRB
PDGFRL
PDHA1
PDHB
PDHX
PDIA3
PDIA4
PDIA5
PDIA6
PDK1
PDK2
PDK3
PDLIM1
PDLIM2
PDLIM3
PDLIM4
PDLIM5
PDLIM7
PDP1
PDPK1
PDPR
PDS5A
PDS5B
PDXDC1
PDXK
PDZD11
PDZD8
PDZK1IP1
PEA15
PEAK1
PEBP1
PECAM1
PEDS1-
UBE2V1
PEF1
PELO
PELP1
PELP1
PEPD
PERP
PES1
PEX1
PEX11B
PEX14
PEX16
PEX19
PEX3
PEX5
PEX6
PFAS
PFDN1
PFDN2
PFDN4
PFDN5
PFDN6
PFKFB2
PFKL
PFKM
PFKM
PFKP
PFN1
PFN2
PFN2
PGAM1
PGAM5
PGAP1
PGD
PGGT1B
PGK1
PGLS
PGLYRP2
PGM1
PGM2
PGM2L1
PGM3
PGM5
PGP
PGR
PGRMC1
PGRMC2
PHAF1
PHB1
PHB2
PHC2
PHC3
PHF1
PHF10
PHF14
PHF23
PHF5A
PHF6
PHGDH
PHIP
PHKA1
PHKB
PHKG2
PHLDA1
PHLDA3
PHLDB1
PHLDB2
PHLPP2
PHOX2B
PHPT1
PHYHD1
PHYHIP
PHYKPL
PI15
PI16
PI3
PI4K2A
PI4K2B
PI4KA
PI4KB
PICALM
PICALM
PICK1
PIGBOS1
PIGG
PIGK
PIGN
PIGO
PIGQ
PIGR
PIGS
PIGT
PIGU
PIGX
PIK3C2A
PIK3C2B
PIK3C3
PIK3CA
PIK3CB
PIK3CD
PIK3R1
PIK3R2
PIK3R4
PIKFYVE
PILRA
PIN1
PIN4
PIP4K2A
PIP4K2B
PIP4K2C
PIP4P1
PIP4P2
PIP5K1A
PIP5K1C
PIR
PITHD1
PITPNA
PITPNB
PITRM1

PKIG
PKM
PKM
PKN1
PKN2
PKP1
PKP2
PKP3
PKP3
PKP4
PLA1A
PLA2G2A
PLA2G4A
PLA2G4B
PLA2G4D
PLA2G4E
PLA2G4F
PLAA
PLAC9
PLAT
PLBD1
PLBD2
PLCB1
PLCB1
PLCB3
PLCB4
PLCD1
PLCD3
PLCD4
PLCG1
PLCG2
PLCH2
PLCH2
PLCL1
PLD1
PLD2
PLD3
PLEC
PLEC
PLEC
PLEK
PLEKHA1
PLEKHA2
PLEKHA5
PLEKHA8
PLEKHF1
PLEKHF2
PLEKHG5
PLEKHM2
PLEKHN1
PLEKHO2
PLG
PLGRKT
PLIN1
PLIN3
PLIN4
PLN
PLOD1
PLOD2
PLOD3
PLP1
PLP2
PLPBP
PLPP1
PLPP3
PLPP6
PLRG1
PLS1
PLS3
PLSCR1
PLSCR3
PLSCR4
PLTP
PLVAP
PLXDC1
PLXDC2
PLXNA1
PLXNA3
PLXNB2
PLXNC1
PLXND1
PM20D2
PMEL
PMF1;PMF1-
BGLAP
PML
PMM2
PMP2
PMPCA
PMPCB
PMVK
PNKP
PNMA1
PNN
PNO1
PNP
PNPLA4
PNPLA6
PNPLA8
PNPO
PNPT1
PODN
PODNL1
PODXL
POF1B
POFUT1
POFUT2
POGLUT1
POGLUT2
POGLUT3
POGZ
POLB
POLD1
POLD2
POLD3
POLDIP2
POLDIP3
POLG
POLI
POLR1A
POLR1B
POLR1C
POLR1D
POLR2A
POLR2B
POLR2C
POLR2D
POLR2E
POLR2F
POLR2G
POLR2H
POLR2I
POLR2L
POLR2M
POLR3A
POLR3B
POLR3F
POM121
POMGNT2
POMP
POMT1
PON1
PON2
POP4
POPDC2
POR
POSTN
POTEI
POTEJ
POU2F1;POU
2F2;POU2F2;
POU2F2;POU
2F2;POU2F2;
POU2F2;POU
2F1;POU2F3;
POU2F2;POU
2F2
PPA1
PPA2
PPA2
PPAT
PPCDC
PPCS
PPFIA1
PPFIBP1
PPFIBP2
PPIA
PPIA
PPIB
PPIC
PPID
PPIE
PPIF
PPIG
PPIH
PPIL1
PPIL3
PPIL4
PPIP5K1
PPIP5K2
PPL
PPM1A
PPM1B
PPM1F
PPM1G
PPM1K
PPM1L
PPME1
PPOX
PPP1CA
PPP1CB
PPP1CC
PPP1CC
PPP1R11
PPP1R12A
PPP1R12B
PPP1R12C
PPP1R13B;TP
53BP2;PPP1
R13B
PPP1R13L
PPP1R14A
PPP1R14B
PPP1R14C
PPP1R18
PPP1R2
PPP1R21
PPP1R3G
PPP1R7
PPP1R8
PPP1R9B
PPP2CB
PPP2R1A
PPP2R1B
PPP2R2A
PPP2R2D
PPP2R5A
PPP2R5B
PPP2R5C
PPP2R5D
PPP2R5E
PPP3CA
PPP3CB
PPP3R1
PPP4C

PPP4R1
PPP4R2
PPP4R3A
PPP5C
PPP6C
PPP6R2
PPP6R3
PPT1
PPT2;PPT2;P
PT2;PPT2;PP
T2;PPT2;PPT2
;PPT2;;;PPT2;
PPT2;PPT2;P
PT2;PPT2
PPTC7
PPWD1
PQBP1
PRAF2
PRAMEF18;P
RAMEF18;PR
AMEF22;PRA
MEF22;PRAM
EF19;PRAME
F18
PRCP
PRCP
PRDX1
PRDX2
PRDX3
PRDX4
PRDX5
PRDX6
PREB
PRELP
PREP
PREPL
PREX1
PRG2
PRG3
PRG4
PRKAA1
PRKAA2
PRKAB1
PRKAB2
PRKACA

PRKACB
PRKAG1
PRKAG2
PRKAR1A
PRKAR2A
PRKAR2B
PRKCA
PRKCB
PRKCD
PRKCE
PRKCI
PRKCSH
PRKD1
PRKD2
PRKD2
PRKDC
PRKG1
PRKG1
PRKRA
PRMT1
PRMT5
PRNP
PROC
PROCR
PROM2
PRORP
PROS1
PROZ
PRP4K
PRPF18
PRPF19
PRPF3
PRPF31
PRPF38A
PRPF38B
PRPF4
PRPF40A
PRPF6
PRPF8
PRPH
PRPS1
PRPS2
PRPSAP1
PRPSAP2
PRR5

PRRC1
PRRC2A
PRRC2B
PRRC2C
PRRX1
PRSS1;PRSS
2;PRSS2;PRS
S1;PRSS1;PR
SS1;PRSS2;P
RSS3P2
PRSS12
PRSS23
PRSS3
PRSS35
PRSS8
PRTFDC1
PRTN3
PRUNE1
PRUNE2
PRX
PRXL2A
PRXL2B
PSAP
PSAPL1
PSAT1
PSCA
PSEN1
PSIP1
PSMA1
PSMA2
PSMA3
PSMA4
PSMA5
PSMA6
PSMA7
PSMB1
PSMB10
PSMB2
PSMB2
PSMB3
PSMB4
PSMB5
PSMB6
PSMB7
PSMB8

PSMB9
PSMC1
PSMC1
PSMC2
PSMC3
PSMC4
PSMC5
PSMC6
PSMD1
PSMD10
PSMD11
PSMD12
PSMD13
PSMD14
PSMD2
PSMD3
PSMD4
PSMD5
PSMD6
PSMD7
PSMD8
PSMD9
PSME1
PSME2
PSME3
PSME4
PSMF1
PSMG1
PSMG2
PSMG3
PSMG4
PSPC1
PSPH
PSTPIP1
PTBP1
PTBP2
PTBP3
PTCD3
PTDSS1
PTDSS2
PTEN
PTER
PTGDS
PTGES
PTGES2

PTGES3
PTGES3L-
AARSD1
PTGFRN
PTGIS
PTGR1
PTGR2
PTGR3
PTGS1
PTGS1
PTK2
PTK2
PTK2B
PTK6
PTK7
PTMA
PTMS
PTN
PTP4A2
PTPA
PTPMT1
PTPN1
PTPN11
PTPN12
PTPN13
PTPN23
PTPN3
PTPN6
PTPN9
PTPRA
PTPRB
PTPRC
PTPRCAP
PTPRE
PTPRF
PTPRG
PTPRK
PTPRM
PTPRS
PTPRU
PTPRZ1
PTRH1
PTRH2
PTRHD1
PTTG1IP

PTX3
PUDP
PUF60
PUM1
PUM2
PUM3
PURA
PURB
PUS1
PUS7
PVR
PWP1
PWP2
PXDN
PXK
PXMP4
PXN
PYCARD
PYCR1
PYCR2
PYCR3
PYGB
PYGL
PYGM
PYGO2
PYM1
PYROXD2
QARS1
QDPR
QKI
QNG1
QPCT
QPCTL
QPRT
QSOX1
QTRT1
RAB10
RAB11B
RAB11FIP1
RAB11FIP5
RAB12
RAB13
RAB14
RAB18
RAB1A
RAB1B
RAB21
RAB22A
RAB23
RAB24
RAB25
RAB27A
RAB27B
RAB29
RAB2A
RAB2B
RAB30
RAB31
RAB32
RAB33B
RAB34
RAB35
RAB38
RAB3C
RAB3D
RAB3GAP1
RAB3GAP2
RAB3IL1
RAB41
RAB43
RAB4A
RAB4B
RAB5A
RAB5B
RAB5B
RAB5C
RAB6A
RAB6D
RAB7A
RAB7B
RAB8A
RAB8B
RAB9A
RABEP1
RABEP2
RABGAP1
RABGAP1L
RABGEF1
RABGGTA
RABGGTB
RABIF
RABL2A;RAB
L2B
RABL3
RABL6
RAC1
RAC2
RAC3
RACK1
RAD21
RAD23A
RAD23B
RAD50
RAE1
RAF1
RAI14
RALA
RALB
RALGAPA1
RALGAPB
RALY
RAMP3
RAN
RANBP1
RANBP10
RANBP2
RANBP3
RANBP9
RANGAP1
RAP1A
RAP1B
RAP1B
RAP1GDS1
RAP2A
RAP2B
RAPGEF1
RAPGEFL1
RAPH1
RARA;RARB;
RARA;RARA;
RARB
RARRES2
RARS1
RARS2
RASA1
RASA3
RASA4
RASAL1
RASAL2
RASAL3
RASGRP2
RASIP1
RASL12
RASSF5
RAVER1
RB1
RB1CC1
RBBP4
RBBP5
RBBP6
RBBP7
RBBP9
RBCK1
RBFOX1;RBF
OX2;RBFOX1
;RBFOX1;RBF
OX1;RBFOX1
;RBFOX1;RBF
OX1;RBFOX1
;RBFOX2;RBF
OX2;RBFOX2
;RBFOX1;RBF
OX2;RBFOX2
;RBFOX1;RBF
OX1;RBFOX1
;RBFOX1;RBF
OX1;RBFOX2
;RBFOX1;RBF
OX2
RBM10
RBM12
RBM12B
RBM14
RBM15
RBM17
RBM22
RBM25
RBM26
RBM27
RBM28
RBM3
RBM39
RBM4
RBM42
RBM7;RBM7;
RBM7;;RBM7
;RBM7
RBM8A
RBMS1
RBMX
RBMXL1
RBP1
RBP4
RBP7
RBPJ
RBPMS
RBPMS2
RBSN
RBX1
RCBTB2
RCC1
RCC2
RCE1
RCL1
RCN1
RCN2
RCN3
RCOR3
RCSD1
RDH10
RDH11
RDH12
RDH13
RDH14
RDX
RECK
RECQL
RECQL5
REEP3
REEP4
REEP5
REL
RELA
RELCH
REM1

RENBP
REPS1
RER1
RERG
RETN
RETREG2
RETREG3
RETSAT
REXO2
RFC1
RFC2
RFC3
RFC4
RFC5
RFFL
RFT1
RFTN1
RFX5
RGN
RGS12
RHAG
RHBDD1
RHBDF1
RHBDF2
RHCE
RHCG
RHEB
RHOA
RHOB
RHOC
RHOG
RHOJ
RHOT1
RHOT1
RHOT2
RIC1
RIC8A
RICTOR
RIDA
RIGI
RIIAD1
RILPL1
RIMOC1
RIN1
RIN3
RING1
RINT1
RIOK3
RIOX1
RIPK1
RIPOR1
RIT1
RLIG1
RMC1
RMDN1
RMDN2
RMDN3
RMND5A
RNASE1
RNASE3
RNASE4
RNASE6
RNASEH2B
RNASEH2C
RNASEK
RNASEL
RNASET2
RND3
RNF114
RNF121;RNF
121;RNF175;
RNF121
RNF123
RNF126
RNF14
RNF170
RNF181
RNF2
RNF213
RNF31
RNF39
RNF7
RNH1
RNMT
RNPEP
RNPS1
RO60
ROBO1
ROCK1
ROCK2
ROR2
RP2
RPA1
RPA2
RPA3
RPAP1
RPE
RPF2
RPIA
RPL10
RPL10A
RPL11
RPL12
RPL13
RPL13A
RPL14
RPL15
RPL15
RPL17;RPL17
-
C18orf32;RP
L17;RPL17;R
PL17
RPL18
RPL18A
RPL19
RPL21
RPL22
RPL22L1
RPL23
RPL23A
RPL24
RPL26
RPL27
RPL27A
RPL28
RPL29
RPL3
RPL30
RPL31
RPL32
RPL34
RPL35
RPL35A
RPL36
RPL36A;RPL3
6A-
HNRNPH2;R
PL36A;RPL36
A
RPL36AL
RPL37A
RPL38
RPL39;RPL39
P5
RPL4
RPL5
RPL6
RPL7
RPL7A
RPL7L1
RPL8
RPL9
RPLP0
RPLP1
RPLP2
RPN1
RPN2
RPP25L
RPP30
RPP40
RPRD1A
RPRD1B
RPRD2
RPS10
RPS10-
NUDT3
RPS11
RPS12
RPS13
RPS14
RPS15
RPS15A
RPS16
RPS18
RPS19
RPS2
RPS20
RPS21
RPS23
RPS24
RPS25
RPS26
RPS27
RPS27A
RPS27L
RPS28
RPS29
RPS3
RPS3A
RPS4X
RPS4Y1
RPS5
RPS6
RPS6KA1
RPS6KA3
RPS6KA4
RPS6KA5
RPS6KB1
RPS6KB2
RPS7
RPS7
RPS8
RPS9
RPSA
RPTN
RPTOR
RRAD
RRAGA
RRAGC
RRAS
RRAS2
RRBP1
RRM1
RRM2B
RRN3
RRP1
RRP12
RRP1B
RRP7A
RRP9
RRS1
RSBN1
RSKR;;RSKR;
RSKR;RSKR

RSL1D1
RSL24D1
RSPRY1
RSU1
RTCA
RTCB
RTF1
RTF2
RTKN
RTL8C
RTN1
RTN2
RTN3
RTN3
RTN4
RTRAF
RUFY1
RUFY2
RUFY3
RUNX2
RUSC1
RUSF1
RUVBL1
RUVBL2
RWDD1
RWDD4
RXRA
RXRB
RYR2
S100A1
S100A10
S100A11
S100A12
S100A13
S100A14
S100A16
S100A2
S100A3
S100A4
S100A6
S100A7
S100A7L2
S100A8
S100A9
S100B
S100P
S1PR2
SAA2-SAA4
SACM1L
SACS
SAE1
SAFB
SAFB2
SAMD4B
SAMD9
SAMD9L
SAMHD1
SAMM50
SAP18
SAP30BP
SAR1A
SAR1B
SARG
SARM1
SARNP
SARS1
SARS2
SART1
SART3
SASH1
SAT2
SATB2
SAV1
SBDS
SBF1
SBF2
SBSN
SBSPON
SCAF4
SCAI
SCAMP1
SCAMP2
SCAMP3
SCAMP4
SCARA5
SCARB2
SCARF2
SCCPDH
SCD5
SCEL
SCEL
SCFD1
SCFD2
SCGB1A1
SCIN
SCLY
SCN7A
SCN8A
SCN9A
SCO1
SCOC
SCP2
SCPEP1
SCRIB
SCRIB
SCRIB
SCRN1
SCRN2
SCRN3
SCUBE1
SCUBE3
SCYL1
SCYL2
SCYL3
SDAD1
SDC1
SDC2
SDCBP
SDCBP
SDCBP2
SDF2
SDF2L1
SDF4
SDHA
SDHB
SDHC
SDHD
SDK1
SDR16C5
SDR39U1
SDR9C7
SEC11A
SEC11C
SEC13
SEC14L2
SEC16A
SEC22A
SEC22B
SEC23A
SEC23A
SEC23B
SEC23IP
SEC24A
SEC24B
SEC24C
SEC24D
SEC31A
SEC61A1
SEC61B
SEC61G
SEC62
SEC63
SEH1L
SEL1L
SELENBP1
SELENOF
SELENOH
SELENOI
SELENOM
SELENOO
SELENOP
SELENOS
SELENOT
SELP
SEMA3B
SEMA3C
SEMA4B
SEMG1
SENP3-EIF4A1;SENP3
SEPHS1
SEPHS2
SEPSECS
SEPTIN1
SEPTIN10
SEPTIN11
SEPTIN2
SEPTIN2
SEPTIN3
SEPTIN4
SEPTIN5
SEPTIN5
SEPTIN6
SEPTIN7
SEPTIN8
SEPTIN9
SERAC1
SERBP1
SERF2
SERINC1
SERINC3
SERINC5
SERPINA1
SERPINA1
SERPINA10
SERPINA12
SERPINA3
SERPINA4
SERPINA5
SERPINA6
SERPINA7
SERPINB1
SERPINB10
SERPINB11
SERPINB12
SERPINB13
SERPINB13
SERPINB2
SERPINB3
SERPINB4
SERPINB5
SERPINB6
SERPINB7
SERPINB8
SERPINB9
SERPINC1
SERPIND1
SERPINE2
SERPINF1
SERPINF2
SERPING1
SERPINH1
SESTD1
SET

SETD1A
SETD3
SETD7
SETMAR
SF1
SF3A1
SF3A2
SF3A3
SF3B1
SF3B2
SF3B3
SF3B4
SF3B5
SF3B6
SFN
SFPQ
SFRP1
SFRP2
SFRP4
SFT2D2
SFXN1
SFXN2
SFXN3
SFXN4
SGCB
SGCD
SGCD
SGCE
SGK3
SGPL1
SGSH
SGTA
SGTB
SH3BGRL
SH3BGRL2
SH3BGRL3
SH3BP1
SH3D19
SH3GL1
SH3GLB1
SH3GLB2
SH3KBP1
SH3PXD2B
SH3RF2
SHANK1
SHANK3
SHBG
SHC1
SHISA4
SHMT1
SHMT2
SHOC2
SHROOM3
SHTN1
SIDT2
SIGLEC1
SIGMAR1
SIK3
SIL1
SIN3A
SIPA1
SIRPA
SIRT2
SIRT3
SIRT5
SKAP1
SKIC2
SKIC3
SKIC8
SLAIN2
SLC11A1
SLC11A2
SLC12A2
SLC12A4
SLC12A7
SLC12A9
SLC15A3
SLC16A1
SLC16A3
SLC17A5
SLC18B1
SLC1A3
SLC1A4
SLC1A5
SLC20A1
SLC22A18
SLC22A3
SLC25A1
SLC25A1
SLC25A11
SLC25A12
SLC25A13
SLC25A17
SLC25A19
SLC25A20
SLC25A22
SLC25A24
SLC25A25
SLC25A3
SLC25A3
SLC25A32
SLC25A4
SLC25A40
SLC25A42
SLC25A44
SLC25A5
SLC25A52;SLC25A52;SLC25A51
SLC25A6
SLC26A2
SLC27A1
SLC27A3
SLC27A4
SLC27A6
SLC29A1
SLC2A1
SLC2A10
SLC2A4
SLC30A1
SLC30A5
SLC30A6
SLC30A7
SLC31A1
SLC33A1
SLC35A1
SLC35A2
SLC35A3
SLC35A4
SLC35B1
SLC35B2
SLC35C1
SLC35E3
SLC35F6
SLC37A2
SLC38A7
SLC39A11
SLC39A14
SLC39A2
SLC39A7
SLC3A2
SLC41A3
SLC43A3
SLC44A1
SLC44A2
SLC4A1
SLC4A1AP
SLC4A2
SLC4A7
SLC52A2
SLC52A3
SLC66A3
SLC6A2
SLC7A1
SLC8A1
SLC9A1
SLC9A6
SLC9A9
SLCO2A1
SLFN5
SLIRP
SLIT2
SLK
SLMAP
SLPI
SLTM
SLURP1
SLURP2
SMAD1
SMAD2
SMAD3
SMAD4
SMAP1
SMAP2
SMARCA1
SMARCA2
SMARCA4
SMARCA5
SMARCAL1
SMARCB1
SMARCC1
SMARCC2
SMARCD1
SMARCD2
SMARCE1
SMC1A
SMC2
SMC3
SMCHD1
SMDT1
SMG1
SMG8
SMG9
SMIM12
SMIM20
SMIM5
SMNDC1
SMOC1
SMPD1
SMPD4
SMS
SMTN
SMTN
SMU1
SMUG1
SMURF1
SMYD5
SNAP23
SNAP29
SNAPIN
SNCA
SNCG
SND1
SNF8
SNRNP200
SNRNP40
SNRNP70
SNRPA
SNRPA1
SNRPB2
SNRPC
SNRPD1
SNRPD2
SNRPD3
SNRPE

SNRPF
SNRPGP15;S
NRPG
SNRPN;SNR
PB;SNRPN
SNTA1
SNTB1
SNTB2
SNU13
SNU13
SNW1
SNX1
SNX12
SNX14
SNX15
SNX17
SNX18
SNX2
SNX21
SNX24
SNX27
SNX3
SNX33
SNX4
SNX5
SNX6
SNX7
SNX8
SNX9
SOAT1
SOD1
SOD2
SOD3
SON
SORBS1
SORBS1
SORBS1
SORBS1
SORBS2
SORBS2
SORBS2
SORBS3
SORD
SORT1
SOS1
SOSTDC1
SOWAHC
SP1
SP100
SP3
SPAG7
SPAG9
SPARC
SPARCL1
SPART
SPATA20
SPATS2L
SPCS1
SPCS2
SPCS3
SPECC1
SPECC1L
SPG11
SPG21
SPG7
SPHK1
SPINK5
SPINK7
SPINT1
SPINT2
SPNS1
SPON1
SPON2
SPP2
SPPL2A
SPPL2B
SPR
SPRR1A
SPRR1B
SPRR2A
SPRR2D
SPRR2E
SPRR3
SPRYD3
SPRYD4
SPTA1
SPTAN1
SPTAN1
SPTAN1
SPTB
SPTBN1
SPTBN2
SPTLC1
SPTLC2
SPTLC3
SQLE
SQOR
SQSTM1
SRC
SRD5A2
SRGAP2
SRI
SRM
SRP14
SRP19
SRP54
SRP68
SRP72
SRP9
SRPK1
SRPK2
SRPRA
SRPRB
SRPX
SRPX2
SRRM1
SRRM2
SRRT
SRSF1
SRSF10
SRSF11
SRSF2
SRSF3
SRSF4
SRSF7
SRSF9
SRXN1
SSB
SSB
SSBP1
SSBP3
SSC5D
SSH3
SSNA1
SSPN
SSR1
SSR2
SSR3
SSR4
SSRP1
SSU72
ST13
ST3GAL4
ST3GAL6
ST6GALNAC4
ST7
ST8SIA3
STAB1
STAG1
STAG2
STAG3
STAM
STAM2
STAMBP
STAP2
STARD10
STARD13
STARD5
STAT1
STAT2
STAT3
STAT3
STAT5A
STAT5B
STAT6
STAU1
STBD1
STC1
STEAP3
STEAP4
STIM1
STING1
STIP1
STK10
STK11IP
STK17B
STK24
STK25
STK26
STK3
STK32A
STK38
STK38L
STK39
STK4
STMN1
STOM
STOML2
STON1-
GTF2A1L
STON2
STRAP
STRIP1
STRN
STRN3
STRN4
STS
STT3A
STT3B
STUB1
STUM
STX10
STX12
STX16-
NPEPL1;STX1
6
STX17
STX18
STX2
STX3
STX4
STX5
STX7
STXBP1
STXBP1
STXBP2
STXBP3
STXBP4
STXBP5
SUB1
SUCLA2
SUCLG1
SUCLG2
SUFU
SUGP1

SUGT1
SULF1
SULT1A2
SULT1A4;SUL
T1A3;SULT1A
4
SULT1E1
SULT2B1
SUMF2
SUMO1
SUMO2
SUN1
SUN1
SUN1
SUN2
SUOX
SUPT16H
SUPT4H1
SUPT5H
SUPT6H
SUPV3L1
SURF1
SURF4
SUSD2
SUZ12
SVIL
SWAP70
SYAP1
SYDE1
SYK
SYMPK
SYN1
SYN2
SYNC
SYNCRIP
SYNE1
SYNE2
SYNE3
SYNGR1
SYNGR2
SYNGR3
SYNJ1
SYNJ2BP
SYNJ2BP-
COX16
SYNM
SYNPO
SYNPO2
SYNPO2
SYP
SYPL1
SYPL1
SYPL2
SYT15B;SYT1
5;SYT15;SYT1
5B
SYTL1
SYTL2
SYVN1
SZT2
TAB1
TACC1
TACC2
TACO1
TACSTD2
TAF15
TAF9;TAF9;TA
F9;TAF9;TAF9
;TAF9;TAF9B
TAGLN
TAGLN2
TAGLN3
TALDO1
TAMM41
TANC1
TANGO2
TAOK1
TAOK2
TAOK3
TAP1
TAP2
TAP2
TAPBP
TARBP1
TARDBP
TARS1
TARS2
TARS3
TATDN1
TAX1BP3
TBC1D1
TBC1D10A
TBC1D10B
TBC1D13
TBC1D15
TBC1D17
TBC1D22A
TBC1D23
TBC1D24
TBC1D2B
TBC1D4
TBC1D5
TBC1D8B
TBC1D9B
TBCA
TBCB
TBCC
TBCD
TBCE
TBCEL
TBK1
TBL1XR1
TBL2
TBL3
TBP
TBRG4
TBXA2R
TBXAS1
TCAF2
TCEA1
TCEA3
TCEAL3
TCEAL4
TCERG1
TCF25
TCIRG1
TCOF1
TCP1
TCP11L1
TEAD3
TECR
TEP1
TERF2
TERF2IP
TES
TEX10
TEX2
TEX264
TF
TFAM
TFAP2A
TFB1M
TFCP2
TFG
TFRC
TGFB1I1
TGFBI
TGFBI
TGFBR3
TGFBRAP1
TGM1
TGM2
TGM2
TGM5
TGOLN2
TH
THADA
THAP11
THAP12
THBD
THBS1
THBS2
THBS3
THBS4
THEM6
THG1L
THNSL1
THOC1
THOC2
THOC3
THOC6
THOC7
THOP1
THRAP3
THSD4
THSD7A
THTPA
THUMPD1
THUMPD3
THY1
THYN1
TIA1
TIAL1
TIGAR
TIMM10
TIMM10B
TIMM13
TIMM17B
TIMM21
TIMM22
TIMM23
TIMM29
TIMM44
TIMM50
TIMM8A
TIMM8B
TIMM9
TIMMDC1
TIMP1
TIMP2
TIMP3
TINAGL1
TINF2
TIPRL
TJP1
TJP1
TJP2
TK1
TK2
TKFC
TKT
TLCD1
TLCD3A
TLE3
TLE4;TLE1;TL
E4
TLE5
TLK2
TLN1
TLN2
TLR3
TM2D1
TM2D3
TM4SF1
TM4SF18

TM7SF2
TM9SF2
TM9SF3
TM9SF4
TMA16
TMBIM1
TMCO1
TMCO4
TMED1
TMED10
TMED2
TMED3
TMED4
TMED5
TMED7-
TICAM2;TME
D7
TMED9
TMEM106B
TMEM109
TMEM11
TMEM119
TMEM120A
TMEM123
TMEM126A
TMEM128
TMEM134
TMEM147
TMEM14C
TMEM154
TMEM161A
TMEM165
TMEM167A
TMEM167B
TMEM168
TMEM179B
TMEM184B
TMEM19
TMEM192
TMEM201
TMEM205
TMEM214
TMEM222
TMEM223
TMEM231
TMEM245
TMEM256-
PLSCR3;TME
M256-
PLSCR3;TME
M256
TMEM258
TMEM259
TMEM263
TMEM30A
TMEM33
TMEM35A
TMEM40
TMEM41A
TMEM43
TMEM45A
TMEM47
TMEM50A
TMEM50B
TMEM63A
TMEM63B
TMEM68
TMEM70
TMEM79
TMEM87A
TMEM87B
TMEM94
TMEM97
TMF1
TMLHE
TMOD1
TMOD2
TMOD2
TMOD3
TMPO
TMPO
TMPPE
TMPRSS11A
TMPRSS11B
TMPRSS11D
TMPRSS11E
TMSB10
TMSB4X
TMT1A
TMTC3
TMUB1
TMX1
TMX2
TMX3
TMX4
TNC
TNFAIP2
TNFAIP6
TNFAIP8
TNFAIP8L3
TNFRSF6B
TNFSF12
TNFSF12-
TNFSF13;TNF
SF13;TNFSF1
3
TNFSF13
TNIP1
TNK1
TNKS1BP1
TNMD
TNNI2
TNPO1
TNPO2
TNPO3
TNS1
TNS1
TNS2
TNS3
TNXB
TOLLIP
TOM1
TOM1L1
TOM1L2
TOMM20
TOMM22
TOMM34
TOMM40
TOMM5
TOMM6
TOMM70
TOP1
TOP2B
TOP3B
TOR1A
TOR1AIP1
TOR1AIP1
TOR1AIP2
TOR1B
TOR2A
TOR3A
TP53BP1
TP53I11
TP53I13
TP53I3
TP53RK
TP63
TPBG
TPCN1
TPD52
TPD52L1
TPD52L2
TPI1
TPK1
TPM1
TPM1
TPM2
TPM2
TPM3
TPM4
TPM4
TPMT
TPP1
TPP2
TPPP
TPPP3
TPR
TPRG1
TPRG1L
TPRKB
TPSAB1
TPSB2
TPT1
TRA2A
TRA2B
TRADD
TRAF3IP3
TRAF6
TRAF7
TRAFD1
TRAM1
TRAP1
TRAPPC1
TRAPPC10
TRAPPC11
TRAPPC12
TRAPPC13
TRAPPC2;TR
APPC2B
TRAPPC2L
TRAPPC3
TRAPPC4
TRAPPC5
TRAPPC6A
TRAPPC6B
TRAPPC8
TRAPPC9
TREX1
TREX2
TRIAP1
TRIM14
TRIM16
TRIM2;;TRIM
2;TRIM2;TRI
M2;TRIM2;TR
IM2;TRIM2;;;T
RIM2
TRIM21
TRIM22
TRIM25
TRIM26
TRIM28
TRIM29
TRIM3
TRIM32
TRIM33
TRIM38
TRIM4
TRIM47
TRIM5
TRIM56
TRIM65
TRIM66
TRIO
TRIOBP

TRIP10
TRIP11
TRIP12
TRIP4
TRIP6
TRIR
TRMT1
TRMT10C
TRMT112
TRMT2A
TRMT6
TRMT61A
TRNT1
TRPC4AP
TRPM4
TRPM7
TRPT1
TRPV1
TRPV2
TRRAP
TSC1
TSC2
TSC22D1
TSC22D3
TSC22D4
TSEN34
TSEN54
TSFM
TSG101
TSKU
TSN
TSNAX
TSPAN13
TSPAN14
TSPAN15
TSPAN18
TSPAN3
TSPAN31
TSPAN4
TSPAN6
TSPAN7;;TSP
AN7
TSPAN8
TSPAN9
TSPO
TSR1
TSR2
TSSC4
TST
TSTD1
TTC1
TTC19
TTC21B
TTC22
TTC27
TTC28
TTC33
TTC38
TTC39A
TTC39B
TTC39C
TTC7A
TTC7B
TTI2
TTLL12
TTPAL
TTR
TUBA1B
TUBA1C
TUBA3E
TUBA4A
TUBAL3
TUBB
TUBB1
TUBB2A
TUBB2B
TUBB3
TUBB4A
TUBB4B
TUBB6
TUBG1
TUBGCP2
TUBGCP3
TUBGCP4
TUFM
TUFT1
TULP1;TUB;T
ULP1;TUB
TUT1
TUT7
TWF1
TWF2
TWSG1
TXLNA
TXN
TXN2
TXNDC12
TXNDC15
TXNDC17
TXNDC5
TXNDC9
TXNIP
TXNL1
TXNL4A
TXNL4B
TXNRD1
TXNRD2
TYMP
TYRO3
TYRP1
U2AF1
U2AF2
U2SURP
UACA
UAP1
UAP1L1
UBA1
UBA2
UBA2
UBA3
UBA5
UBA6
UBA7
UBAC1
UBAC2
UBAP1
UBAP2
UBAP2L
UBE2A
UBE2D2
UBE2E2
UBE2F
UBE2G1
UBE2G2
UBE2H
UBE2I
UBE2K
UBE2L3
UBE2L6
UBE2M
UBE2N
UBE2O
UBE2R2
UBE2V1
UBE2V2
UBE2Z
UBE3A
UBE3C
UBE4A
UBE4B
UBFD1
UBL3
UBL4A
UBL5
UBL7
UBLCP1
UBP1
UBQLN1
UBQLN2
UBR1
UBR2
UBR3
UBR4
UBR5
UBR7
UBTF
UBXN1
UBXN4
UBXN6
UBXN7
UCHL1
UCHL3
UCHL5
UCKL1
UEVLD
UFC1
UFD1
UFL1
UFM1
UFSP2
UGDH
UGGT1
UGP2
UGP2
UGT1A6
UGT1A9
ULK3
UMPS
UNC119B
UNC13B
UNC13D
UNC45A
UNG
UNK
UPF1
UPF2
UPK1A
UPK3BL1;UP
K3BL2
UPP1
UQCC1
UQCC2
UQCC3
UQCR10
UQCR11
UQCRB
UQCRB
UQCRC1
UQCRC2
UQCRFS1
UQCRH
UQCRQ
URB1
URGCP
URI1
URM1
UROD
USF1
USO1
USP10
USP11
USP13
USP14
USP15
USP16

| | | | | |
|---|---|---|---|---|
| USP19 | VIP | VSNL1 | WDR70 | XRCC5 |
| USP24 | VIRMA | VTA1 | WDR74 | XRCC6 |
| USP34 | VIT | VTI1A | WDR75 | XRN1 |
| USP39 | VKORC1L1 | VTI1B | WDR77 | XRN2 |
| USP4 | VKORC1L1 | VTN | WDR81 | XXYLT1 |
| USP40 | VMA21 | VWA1 | WDR82 | XYLB |
| USP46 | VOPP1 | VWA5A | WDR91 | YAP1 |
| USP47 | VPS11 | VWA8 | WFDC12 | YARS1 |
| USP48 | VPS11 | VWF | WFDC2 | YARS2 |
| USP5 | VPS13A | WAC | WFDC5 | YBX1 |
| USP7 | VPS13C | WAPL | WFS1 | YBX3 |
| USP8 | VPS13D | WARS1 | WIPF1 | YEATS2 |
| USP9X | VPS16 | WARS2 | WIPF2 | YES1 |
| UTP15 | VPS18 | WAS | WIPI2 | YIF1A |
| UTP20 | VPS25 | WASF2 | WLS | YIF1B |
| UTP4 | VPS26A | WASH3P;WA | WNK1 | YIPF3 |
| UTRN | VPS26B | SH2P | WNT10A | YIPF4 |
| UVRAG | VPS26C | WASHC2A | WNT10B | YIPF5 |
| VAC14 | VPS28 | WASHC3 | WNT11 | YKT6 |
| VAMP2;VAMP | VPS29 | WASHC4 | WNT16 | YLPM1 |
| 2;;VAMP2 | VPS33A;VPS | WASHC5 | WNT2 | YME1L1 |
| VAMP3 | 33A;VPS33A; | WASL | WNT2B | YOD1 |
| VAMP5 | ;VPS33A | WBP2 | WNT3 | YRDC |
| VAMP7 | VPS33B | WDFY1 | WNT4 | YTHDC1 |
| VAMP8 | VPS35 | WDR1 | WNT5A | YTHDC2 |
| VANGL1 | VPS35L | WDR11 | WNT5B | YTHDF1 |
| VANGL2 | VPS36 | WDR12 | WNT6 | YTHDF2 |
| VAPA | VPS37A | WDR13 | WNT7B | YTHDF3 |
| VAPB | VPS37B | WDR18 | WNT9A | YWHAB |
| VARS1 | VPS37C | WDR20 | WNT9B | YWHAE |
| VASN | VPS39 | WDR24 | WRAP73 | YWHAG |
| VASP | VPS41 | WDR26 | WRNIP1 | YWHAH |
| VAT1 | VPS45 | WDR3 | WSCD1 | YWHAQ |
| VAT1L | VPS4A | WDR36 | WTAP | YWHAZ |
| VBP1 | VPS4B | WDR37 | XAB2 | YWHAZ |
| VCAN | VPS50 | WDR41 | XG | YY1 |
| VCL | VPS51 | WDR43 | XPC | ZBED1 |
| VCP | VPS52 | WDR44 | XPNPEP1 | ZBTB20 |
| VCPIP1 | VPS53 | WDR45 | XPNPEP2 | ZBTB47 |
| VDAC1 | VPS54 | WDR45B | XPO1 | ZBTB7A |
| VDAC2 | VPS8 | WDR47 | XPO4 | ZBTB8OS |
| VDAC3 | VRK1 | WDR48 | XPO5 | ZC2HC1A |
| VEGFB | VRK2 | WDR5 | XPO7 | ZC3H11A |
| VEZF1 | VRK3 | WDR55 | XPOT | ZC3H14 |
| VIM | VSIG10L | WDR6 | XRCC1 | ZC3H15 |

ZC3H18
ZC3H4
ZC3H7A
ZC3H7B
ZC3HAV1
ZC3HAV1L
ZC3HC1
ZCCHC3
ZCCHC4
ZDHHC13
ZDHHC20
ZDHHC21
ZDHHC3
ZDHHC5
ZEB2
ZER1
ZFAND1
ZFAND2B
ZFP36L1
ZFPL1
ZFR
ZFYVE1
ZFYVE16
ZFYVE19
ZFYVE21
ZFYVE26
ZG16
ZG16B
ZHX3
ZMIZ1
ZMPSTE24
ZMYND11
ZMYND8
ZNF143
ZNF185
ZNF185
ZNF207
ZNF232
ZNF326
ZNF384
ZNF385A
ZNF426
ZNF496
ZNF512
ZNF512B
ZNF573
ZNF598
ZNF638
ZNF677
ZNF687
ZNF706
ZNF75A
ZNF814;ZNF417;ZNF417;ZNF587
ZNFX1
ZNG1B;ZNG1A
ZNRD2
ZPR1
ZRANB2
ZSCAN18
ZSCAN30
ZSWIM8
ZW10
ZYG11B
ZYX
ZZEF1
